# Integrated multimodal cell atlas of Alzheimer’s disease

**DOI:** 10.1101/2023.05.08.539485

**Authors:** Mariano I. Gabitto, Kyle J. Travaglini, Victoria M. Rachleff, Eitan S. Kaplan, Brian Long, Jeanelle Ariza, Yi Ding, Joseph T. Mahoney, Nick Dee, Jeff Goldy, Erica J. Melief, Anamika Agrawal, Omar Kana, Xingjian Zhen, Samuel T. Barlow, Krissy Brouner, Jazmin Campos, John Campos, Ambrose J. Carr, Tamara Casper, Rushil Chakrabarty, Michael Clark, Jonah Cool, Rachel Dalley, Martin Darvas, Song-Lin Ding, Tim Dolbeare, Tom Egdorf, Luke Esposito, Rebecca Ferrer, Lynn E. Fleckenstein, Rohan Gala, Amanda Gary, Emily Gelfand, Jessica Gloe, Nathan Guilford, Junitta Guzman, Daniel Hirschstein, Windy Ho, Madison Hupp, Tim Jarksy, Nelson Johansen, Brian E. Kalmbach, Lisa M. Keene, Sarah Khawand, Mitch Kilgore, Amanda Kirkland, Michael Kunst, Brian R. Lee, Mckaila Leytze, Christine L. Mac Donald, Jocelin Malone, Zoe Maltzer, Naomi Martin, Rachel McCue, Delissa McMillen, Gonzalo Mena, Emma Meyerdierks, Kelly P. Meyers, Tyler Mollenkopf, Mark Montine, Amber L. Nolan, Julie Nyhus, Paul A. Olsen, Maiya Pacleb, Chelsea M. Pagan, Nicholas Peña, Trangthanh Pham, Christina Alice Pom, Nadia Postupna, Christine Rimorin, Augustin Ruiz, Giuseppe A. Saldi, Aimee M. Schantz, Nadiya V. Shapovalova, Staci A. Sorensen, Brian Staats, Matt Sullivan, Susan M. Sunkin, Carol Thompson, Michael Tieu, Jonathan Ting, Amy Torkelson, Tracy Tran, Nasmil J. Valera Cuevas, Sarah Walling-Bell, Ming-Qiang Wang, Jack Waters, Angela M. Wilson, David Haynor, Nicole M. Gatto, Suman Jayadev, Shoaib Mufti, Lydia Ng, Shubhabrata Mukherjee, Paul K. Crane, Caitlin S. Latimer, Boaz P. Levi, Kimberly Smith, Jennie L. Close, Jeremy A. Miller, Rebecca D. Hodge, Eric B. Larson, Thomas J. Grabowski, Michael Hawrylycz, C. Dirk Keene, Ed S. Lein

## Abstract

Alzheimer’s disease (AD) is the most common cause of dementia in older adults. Neuropathological and imaging studies have demonstrated a progressive and stereotyped accumulation of protein aggregates, but the underlying molecular and cellular mechanisms driving AD progression and vulnerable cell populations affected by disease remain coarsely understood. The current study harnesses single cell and spatial genomics tools and knowledge from the BRAIN Initiative Cell Census Network to understand the impact of disease progression on middle temporal gyrus cell types. We used image-based quantitative neuropathology to place 84 donors spanning the spectrum of AD pathology along a continuous disease pseudoprogression score and multiomic technologies to profile single nuclei from each donor, mapping their transcriptomes, epigenomes, and spatial coordinates to a common cell type reference with unprecedented resolution. Pseudo-progression analysis showed two major epochs corresponding with a slow early increase in pathology and a later exponential increase that correlated with cognitive decline. The early phase included inflammatory microglial and reactive astrocyte component, as well as a selective loss of Sst+ inhibitory neuron types in superficial cortical layers, loss of myelinating oligodendrocytes, and up-regulation of a re-myelination program by OPCs. The later phase involved loss of excitatory neurons and Pvalb and Vip neuron subtypes also predominantly in superficial layers. These cell vulnerabilities were also seen in prefrontal cortex and replicated by other independent studies when integrated with the BRAIN Initiative reference. Study data and exploratory tools are freely available to accelerate progress in AD research at <u>SEA-AD.org</u>.

## Introduction

Alzheimer’s Disease (AD) is a complex etiology disease characterized by deposition of hallmark pathological peptides and neurodegeneration that progress across partially overlapping neuroanatomical and temporal axes^1,2^. This process is generally believed to follow a stereotyped progression with Amyloid Beta (Aβ) plaques starting in the cerebral cortex^3^ and hyperphosphorylated tau (pTau) aggregation (neurofibrillary tangles) starting in the brainstem/limbic system^4^. Despite being important biomarkers of AD^5^ and notwithstanding decades of efforts, treatment strategies aiming to reduce the burden of these pathological peptides have resulted in, at best, a modest impact on pathology accompanied by significant side effects^6,7^. Single cell and spatial genomics technologies now offer a dramatically higher resolution analysis of complex brain tissues in health and disease that build on decades of observations in the field^8^, and the first studies applying them to AD have begun to identify cellular vulnerabilities and molecular changes with disease^9–17,9,18^. These include molecular characterization of long-noted changes to microglia^12^ and astrocytes^10,14,19^ partly induced by Aβ plaques^20^, identification of excitatory neurons that bear neurofibrillary tangles^21^ and that are selectively vulnerable in disease^11^, a greater appreciation for the role of endothelial and perivascular cells^15^, and a potential role for the strongly disease-associated *APOE4* allele^22^ in regulating oligodendrocyte cholesterol biosynthesis^23^. However, discoveries from each of these studies have yet to be synthesized into a robust and complete understanding of the mechanistic underpinnings of AD because they lack a framework to relate to one another and to understand when, in the decades long course of AD, they occur. A comprehensive assessment of the key cellular events requires integration of multiple data modalities capable of capturing and relating changes in pathology, gene expression, epigenomic changes, and spatial alterations in the tissue at the single cell level with high-resolution characterization of the affected cell types.

Recent work catalyzed by the BRAIN Initiative Cell Census Network (BICCN) and Cell Atlas Network (BICAN) has established best practices in experimental and quantitative analyses of mouse, non-human primate and human brain using single cell genomics, spatial transcriptomics, and Patch-seq methods to characterize cellular properties and build a knowledge base of brain cell types^24–29^. Over 100 cell types can be reliably identified using single nucleus RNA-seq in any cortical area, and alignment across species shows strong conservation of cellular architecture^30,31^ that allows inference of human cellular properties from studies in experimentally tractable mouse and non-human primate models. Systematic BICCN and BICAN efforts are now producing the first brain-wide cell atlases of the mouse^28,32,33^ and human brain^25–27,34,35^, providing robust and highly curated genomically-based reference cell classifications, spatial maps of cellular distributions, and characterization of cellular properties in normal brain. These reference classifications provide an extremely powerful foundational reference, akin to the human genome, to understand the cellular, molecular, and epigenomic underpinnings of Alzheimer’s disease. Furthermore, mapping to this reference allows integration across data modalities and across independent studies to validate findings and leverage a growing knowledge base on the properties and function of cell types that are affected in disease.

The Seattle Alzheimer’s Disease Cell Atlas (SEA-AD) consortium aims to utilize the advanced technologies and best practices for studying human brain from the BRAIN Initiative to produce the highest-resolution, multimodal brain-wide cell atlas of AD mapped to the BICCN foundational references. Once completed, the SEA-AD Atlas will enable systematic characterization and interpretation of the cellular and molecular correlates of AD neuropathology across brain regions. Keys to achieving this goal are 1) the selection of a high quality donor cohort, spanning the full spectrum of AD pathology, chosen from prospective longitudinal cohort studies with well-characterized participants; 2) the use of improved tissue preparation methods that have been shown extensively to produce high quality single nucleus transcriptomics, epigenomics and spatial transcriptomics data^24–27,30,31,34,35^; 3) a deep donor characterization strategy with all analytical methods applied to the same donors, including quantitative image-based neuropathology, single nucleus multiome analysis with high coverage of nuclei and reads per sample, and targeted spatial transcriptomics; 4) mapping profiled cells to the highly granular and curated BICCN cell type reference; and 5) validating cellular phenotypes across cortical areas and independent studies by integrating and mapping to the same cellular taxonomy reference. By combining temporal modeling of disease severity or progression with single nucleus genomics and spatial analyses, this approach provides the most comprehensive understanding to date on the specific, highly granular cell types affected over the course of disease, where those affected cells are located in tissue microarchitecture, and when they are affected as disease progresses.

The current study focused on the middle temporal gyrus (MTG), an area involved in language and semantic memory processing^36^ and higher order visual processing^37^. MTG is the human cortical region currently with the best annotated BICCN cell classification^25,30^, including cellular phenotype data (morphological and physiological properties) available through Patch-seq analysis of neurosurgical specimens^29,38,39^. Numerous studies, from the seminal histopathology of Braak and Braak^4^ to modern longitudinal studies of tau PET imaging ^40–42^, demonstrate that MTG is a transition zone between aging- or preclinical AD-related medial temporal lobe pTau and more advanced stages of AD where neocortical pTau extends across the brain and is strongly correlated with dementia^4,43–50^. Optimized tissue collection and preparation methods produced high quality human brain tissues, and thereby high-quality single nucleus and spatial genomics data, across the range of age and AD pathology. These data were effectively mapped to the BICCN neurotypical reference classification and used to expand that classification to include disease cell states. Machine learning (ML)-based methods were used to quantitate the local burden of neuropathology in analyzed samples. These data were used for pseudo-trajectory analyses to model disease severity or progression and identify cellular and molecular correlates of disease progression.

This integrative framework around the BICCN reference and temporal modeling demonstrated robust and highly selective neuronal vulnerabilities and changes in non-neuronal disease states as a function of disease progression, along with a wide range of temporal molecular changes. Spatial analyses demonstrated the co-localization of vulnerable cell populations largely in supragranular cortical layers. Data integration to the BICCN reference allowed replication of major cellular changes across data modalities, cortical regions, and independent studies. Overall, these analyses suggest two major epochs with different cellular and molecular events in AD progression. The early phase involves low and linearly increasing levels of neuropathology and increasing inflammatory microglial and reactive astrocyte states, with a notably selective loss of Sst inhibitory neurons, loss of myelinating oligodendrocytes, and sharp up-regulation of an OPC differentiation and re-myelination program. Donors in this phase have no cognitive deficits, suggesting this may represent a preclinical stage of AD. The later phase involves exponentially increasing neuropathology, selective loss of both excitatory and inhibitory neuron types, and eventually a broad cellular pathology accompanying rapid cognitive decline in severely affected donors. This strategy and ability to integrate data across studies to a common reference is highly extensible and provides a unifying framework for the AD community.

## Results

### SEA-AD: Multimodal profiling Alzheimer’s disease progression across wide pathological stages

As summarized in **Figure 1**, to construct an integrated multimodal cellular atlas of AD and comorbid related disorders (AD/ADRD) we generated quantitative i) neuropathological measurements, ii) single nucleus RNAseq (snRNA-seq), ATACseq (snATAC-seq), and Multiome (snMultiome) and iii) cellularly resolved spatial transcriptomics (MERFISH) in the middle temporal gyrus (MTG) from a cohort of 84 aged donors that spanned the spectrum of AD pathology (including donors with no pathology and those with both AD pathology and common co-morbidities) (**Fig. 1a**, **Extended Data Fig. 1a**). We collectively profiled 3.4 million high quality nuclei across all modalities, mapping each to one of 139 molecular cell types from an expanded BRAIN Initiative MTG cellular taxonomy^26,30^ that included disease-associated states. A continuous pseudo-progression score (CPS) from the quantitative neuropathology measurements was constructed, which ordered donors along a neuropathological continuum, and increased discovery power to identify molecular and cellular changes. To validate and replicate these results, we generated a similar 1.2M nuclei snRNA-seq dataset from the dorsolateral prefrontal cortex (DLPFC) in the same 84 donors, mapping to a matched BRAIN initiative DLPFC taxonomy. This DLPFC data also allowed the use of a range of published DLPFC AD studies as replication cohorts. We uniformly re-processed 10 publicly available datasets that applied snRNA-seq to 4.3M high quality nuclei from the DLPFC of 707 additional donors that also spanned the spectrum of AD pathology to validate the principal experimental findings. These multimodal datasets (including both raw and processed data), tools to explore them, and tools to map new datasets to this new cellular taxonomy are all available at SEA-AD.org.

**Figure 1:**
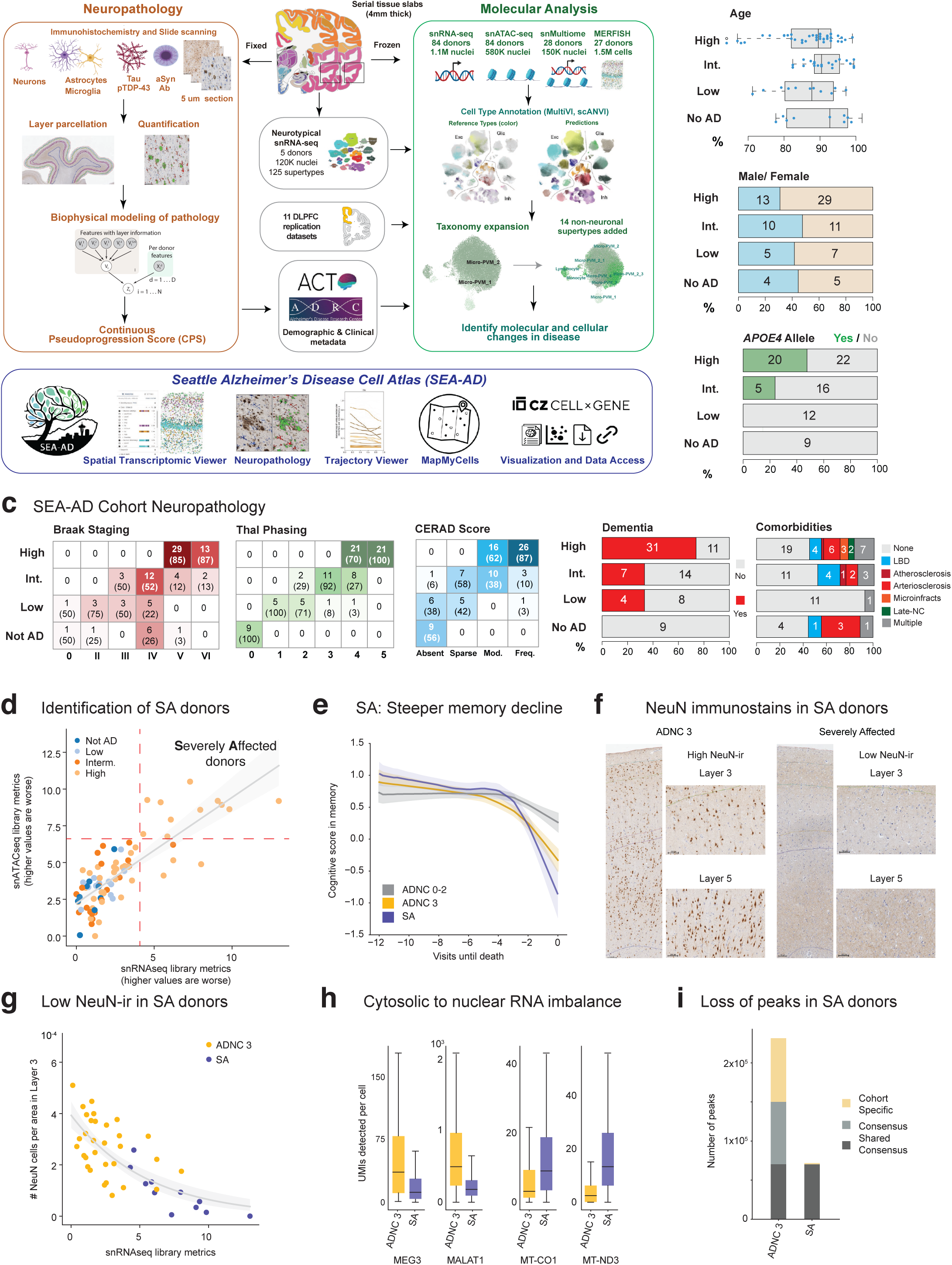
SEA-AD Study of the Middle Temporal Gyrus and Cohort Description. A) Schematic detailing experimental design for applying quantitative neuropathology, single nucleus RNA sequencing (snRNAseq), single nucleus ATAC sequencing (snATACseq), single nucleus Multiome (Multiome), and multiplexed error robust fluorescence in situ hybridization (MERFISH) to middle temporal gyrus (MTG) of SEA-AD donors as well as the analysis plan for construction of a pseudo-progression score from quantitative neuropathology, integration across -omics data modalities, common cell type mapping to the BRAIN initiative reference, use of demographic and clinical metadata to identify cellular and molecular changes in AD, and replication of results across 10 publicly available snRNAseq datasets on the dorsolateral prefrontal cortex (DLPFC) of cohorts that include sporadic AD donors and a snRNAseq dataset generated from the DLPFC of SEA-AD donors. B) SEA-AD cohort demographics stratified by AD neuropathological change (ADNC) score on left. Age at death is represented by box-and-whisker plots where the box represents the interquartile range (IQR) and the whiskers represent 1.5 times the IQR. The median is indicated by the solid line within the box and donor values are represented as points. The fraction of donors that have a biological sex of male or female and that have an APOE4 allele are shown as bar charts per ADNC level, with the number of donors in each group indicated. C) SEA-AD cohort composition stratified by ADNC on the left versus Braak stage (measuring distribution of neurofibrillary tangles across the brain), Thal Phase (measuring distribution of amyloid beta plaques across the brain), and CERAD score (measuring the distribution of neuritic plaques across the brain) as heatmaps. The number of donors in each box are indicated with the fraction in Braak, Thal, and CERAD stages in parentheses. Darker colors represent a higher fraction of donors. The fraction of donors across ADNC that have dementia (red) or not (grey) and any co-morbidities are shown as bar plots. Numbers indicate the number of donors in each group. D) First principal component (PC) with values shifted so the min is set to zero for snRNAseq quality control metrics versus snATACseq quality control metrics for each library color-coded by ADNC category. Dashed red lines indicate the point where values are above 1.5 times the interquartile range. Linear regression is shown in grey, Pearson R=0.80) E) LOESS regression on longitudinal cognitive scores in the memory domain across ADNC 0-2 (Not AD to Intermediate) in grey, ADNC 3 donors that were not severely affected in gold, and ADNC 3 donors that were in purple. Note significant decline in SA donor group. Uncertainty represents the standard error from 1000 LOESS fits with 80% of the data randomly selected in each iteration. F) Exemplar low power micrographs showing the entire cortical column and higher power micrographs of cortical layers 3 and 5 from (left) an ADNC3 (high) donor with NeuN immunoreactive cells and (right) an SA donor case that lacks NeuN-ir (right). Scale bars, 100 um. Thin lines represent cortical layer boundaries. G) Scatterplot showing the number of NeuN immunoreactive cells per area in cortical layer 3 for each donor versus the quality control metrics PC from snRNAseq. SA donors (purple) localize at the end of this trajectory and exhibit almost no immunoreactive cells. Logistic regression is shown in grey. H) Box and whisker plots showing the number of UMIs detected per cell for nuclear-localized RNA species, MEG3 and MALAT1, and mitochondrially encoded (so cytosolically localized) RNA species, MT-CO1 and MT-ND3, for ADNC 3 (high) donors or SA donors. I) Barplot showing the number of chromatin accessible regions in 11 randomly selected ADNC 3 (high) donors or SA donors. *Shared Consensus* (dark grey) accessible regions are regions shared across both groups. *Consensus* (light grey) regions denote regions shared across members of each group and *cohort-specific* (gold) depict peaks unique to some members of each cohort. Note the lack of consensus peaks within SA donors, indicating their peak universe is almost entirely a subset of that found in other ADNC 3 donors.

The SEA-AD cohort was derived from longitudinally characterized research brain donors (mean postmortem interval=7.0 hrs, **Extended Data Fig. 1b**) from the community-based Adult Changes in Thought (ACT) study and the University of Washington (UW) Alzheimer’s Disease Research Center (ADRC)^51–61^. Donors were included in the study if their death occurred within the specific time period of data collection (**Supplementary Table 1**). Brains were collected using highly optimized brain preparation methods that enable exceptionally high-quality snRNA-seq, snATAC-seq, and MERFISH profiling^25–27,34,35^. To assess donor neuropathology, AD has been classically staged by Aβ plaque distribution (Thal phase), neurofibrillary tau tangle distribution (Braak stage), or neuritic plaque density (CERAD score). To account for variability in the rate of plaque and tangle formation across individuals, a composite AD Neuropathological Change (ADNC) scale was developed by the National Institute on Aging – Alzheimer’s Association (NIA-AA) that uses Thal, Braak, and CERAD to categorize donors as either not having AD or having low, intermediate, or high levels of pathology. SEA-AD includes donors at each pathological level (9 Not AD, 12 Low, 21 Intermediate, 42 High ADNC) that were all aged (min. age at death=65, mean=88, **Fig. 1b**, **top**). The cohort contained more female donors (51 females, 33 males), particularly in those with high ADNC (29 females, 13 males), consistent with known prevalence of AD in females^62,63^ (**Fig. 1b**, **middle**). Donors with an *APOE4* allele made up nearly half (20 of 42) of High ADNC cases, nearly a quarter (5 of 21) of Intermediate cases, and no Low ADNC or Not AD cases (**Fig. 1b**, **bottom**), consistent with it being a primary risk factor for late onset AD^22^. Braak stage, Thal phase, and CERAD increased as expected with ADNC (**Fig. 1c**, **left**). Not everyone with AD progresses to develop dementia^64–66^, although the level of pathology is correlated with cognitive decline. Consistent with this, nearly three-quarters (31 of 42) of High ADNC cases had developed dementia prior to death, versus only a third in Intermediate (7 of 21) and Low (4 of 12) ADNC cases, and none in Not AD cases (**Fig. 1c**, **middle**). Donors with any level of Lewy body disease (LBD), vascular pathology, or limbic-predominant age related TDP-43 encephalopathy (LATE) were included as these conditions are common co-morbidities in AD patients^64,67^. Roughly half the cohort (42 of 84) had 1 or more co-pathology (**Fig. 1c**, **right**), enabling exploration of features common across the full spectrum of AD pathology with and without co-pathology.

Nearly all (82 of 84) SEA-AD donors had high pre-sequencing quality control metrics (e.g., brain pH, RIN scores, and sequencing library yield) across the whole range of disease severity (**Extended Data Fig. 1b**). The 2 outlier samples exhibited low RIN values and brain pH and were therefore excluded from further analysis. Basic post sequencing metrics, such as the number of genes and accessible chromatin regions detected per cell, were also uniformly high across disease severity (**Extended Data Fig. 1c**), suggesting there is no inherent tissue quality degradation related to advanced age and neuropathology in most donors. Principal component analysis (PCA) on the detailed snRNA-seq and snATAC-seq library-level metrics did, however, identify a subset of high pathology donors (11 of 42, 26.2%) that had lower quality data in both modalities (**Fig. 1d**, **Extended Data Fig. 2a**). These donors had steeper decline in memory function in their final years of life compared to other high pathology donors (slopes in memory decline = –0.15 in SA donors versus –0.11 in all other high ADNC donors, p-value with Not AD and Low ADNC donors as base outcome = 0.01 versus 0.15, **Fig. 1e**). Longitudinal cognitive testing in our cohort spanned four cognitive domains (memory, executive, language and visuospatial function^68,69^). The remaining cognitive domains showed a similar trajectory across groups (**Extended Data Fig. 2b**). As part of SEA-AD, we measure a battery of quantitative neuropathological variables (described in the next section). When comparing these quantitative measures, these 11 donors had a pronounced reduction in NeuN immunoreactivity (NeuN-ir) in neurons that was not due entirely to cell loss (**Fig. 1f, g**). This is notable as NeuN-ir has previously been shown to be anti-correlated with pTau pathology^70^. Given the steeper cognitive decline and effects on multiple data modalities, we flagged these donors as “Severely Affected” (SA).

Despite having more reads per nucleus, snRNA-seq libraries from these 11 SA donors had fewer unique molecular identifiers overall, genes detected, uniquely mapped reads (which mostly reflects increased ribosomal RNA content) and reads with introns (which reflects mRNA versus pre-mRNA content) (**Extended Data Fig. 2a**). This suggested that cells from the 11 SA donors had less nuclear-localized vs. cytosolic-localized RNA content. Consistent with this, nuclei from the 11 SA donors had lower levels of nuclear-localized RNA^71^ (e.g., *MALAT1* and *MEG3*) and higher levels of cytosolic localized RNA (e.g., RNA from mitochondrially encoded genes) when compared to other high ADNC donors (**Fig. 1h**). To disentangle whether reduced nuclear representation was due to global transcriptional shutdown or degradation, we studied the chromatin landscape in these donors. We computed peaks within each high pathology donor and assessed their similarity by Jaccard distance. The chromatin landscape also segregated the 11 SA donors from matching high pathology donors (**Extended Data Fig. 2c**). We computed consensus peaks across the 11 donors and across matching High ADNC donors (**Methods**) and saw no significant difference in peak-length distribution between groups (**Extended Data Fig. 2d**). However, the 11 SA donors showed many fewer peaks (**Fig. 1i**), which were almost entirely a subset of peaks seen in the high pathology donors. Notably, there were a small number of peaks (n=1,574) unique to the SA donors that were enriched for binding motifs for transcription factors associated with inflammation, de-differentiation and AD pathology (**Extended Data Fig. 2e**). Taken together, these results suggest that SA donors, which suffered from more rapid cognitive decline compared to other high pathology donors, possess nuclei that underwent global chromatin repression which led to shutdown of transcription, consistent with previous reports studying familial AD in which chromatin re-organization triggered neuronal identity repression and de-differentiation^72^. Since SA donors showed systematically lower data quality (**Extended Data Fig 2f**), we excluded these donors from analyses on gene expression changes (described below).

### Quantifying the progression of AD severity

Neuropathological staging is the gold standard for diagnosing AD and relies on semi-quantitative multi-regional assessments of select pathological proteins^3,73,74^. However, their semi-quantitative nature fails to capture the heterogeneity and regional burden of pathology present among donors at each stage. Similar to the concept of an aggregate score for AD staging (ADNC) established by the NIA-AA^64^, we aimed to create a quantitative metric of the local burden of pathology using neuropathological variables (QNP). By modeling disease severity as a continuum that orders the donors from low to high burden of pathology, we could identify earlier and later molecular and cellular events with respect to pathological burden. In addition, a continuous scale of pathological progression yields higher discovery power in statistical analyses, surpassing the effectiveness of traditional categorical staging metrics.

To quantitatively measure neuropathological features that accompany AD progression we used machine learning (ML) approaches to create a mask for each immunohistochemical stain and quantify neuropathological variables (**Extended Data Fig. 3a**). This included stains for well-established markers used for conventional AD neuropathologic staging, including pTau (AT8) for neurofibrillary tangles and Aβ (6e10) for amyloid plaques, as well as additional markers for associated comorbid pathologies and cellular changes. These included pTDP-43 for limbic-predominant, age-related TDP-43 encephalopathy (LATE), alpha synuclein (α-Syn) for Lewy body disease, IBA1 for microglia (including activated states), GFAP for astrocytes (including reactive states), NeuN for neurons, and hematoxylin and eosin (H&E) to assess cytopathology and white matter integrity (**Fig. 2a, b**, **Supplementary Table 2**). These latter cellular markers capture aspects of pathology not typically used in AD neuropathologic staging.

**Figure 2:**
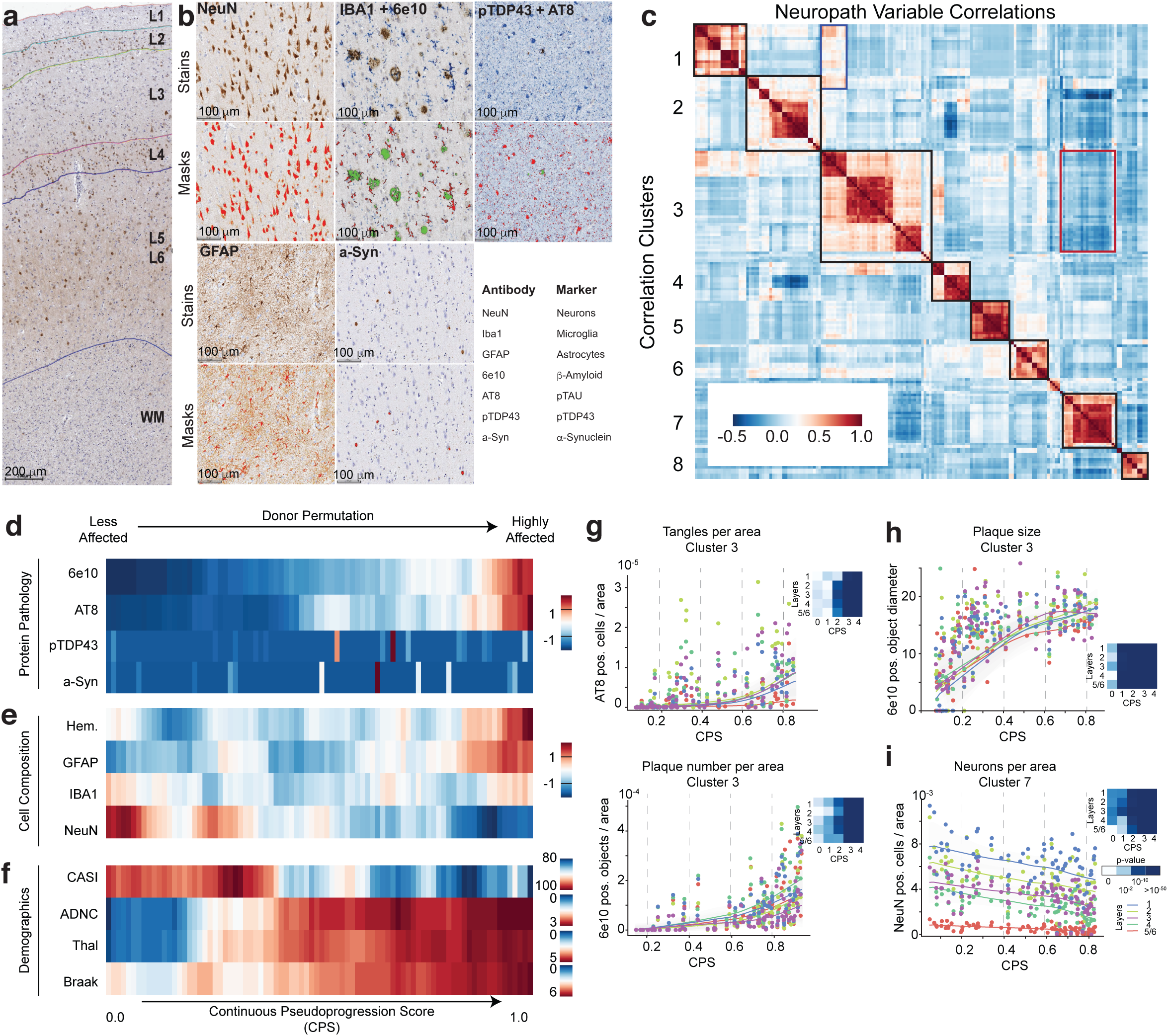
MTG quantitative neuropathology orders donors according to pseudo-progression of disease. A) Representative slide showing the whole cortical column with NeuN visualized with immunohistochemistry (IHC). Cortical layers (L1 to L6) and white matter (WM) are annotated using machine learning with adjustments made by expert neuropathologists. Layer 5 and 6 could not be distinguished by the model so were segmented together. Scale bar, 200 um. B) Higher powered representative micrographs showing IHC stains for protein aggregates (Aβ (6e10), pTau (AT8), α-Syn, and pTDP43) and cellular populations (neurons (NeuN), microglia (Iba1), and astrocytes (GFAP)). Bottom, masks showing positive voxels generated by HALO n red for single stains (NeuN, GFAP, and a-Syn) and both red (IBA1 and AT8) and green (6e10 and pTDP43) for duplex stains. Scale bars, 100 um. C) Heatmap showing hierarchically organized co-correlation matrix of quantitative neuropathology variables. Black boxes on the diagonal, seven clusters that were identified. Red box, anti-correlation between clusters 3 and 7, which represent AD protein pathologies and NeuN immunoreactivity (NeuN-ir), respectively. Blue box, correlation between variables related to neurofibrillary tangles in cluster 3 and pTDP43 variables in cluster 1. D) Heatmap showing the number of pathological protein objects detected per unit area across all cortical layers in each donor, ordered along the continuous pseudo-progression score (CPS). 6e10, Aβ objects; AT8, pTau bearing cells; α-Syn, α-synuclein bearing cells; and pTDP43, pTDP43 bearing cells. All values were converted to z-scores and adjusted by a moving average. E) Heatmap showing the number of cellular objects detected per unit area across all cortical layers in each donor, ordered along the continuous pseudo-progression score (CPS). Hem, all hematoxylin positive nuclei; GFAP, GFAP positive cells; IBA1, IBA1 positive cells; and NeuN, NeuN positive cells. All values were converted to z-scores and adjusted by a moving average. F) Heatmap showing cognitive scores at last visit (CASI) and brain-wide AD pathology stage (ADNC, Thal, Braak) in each donor, ordered along the continuous pseudo-progression score (CPS). All values were adjusted by a moving average. Note, none of these measurements were used to build CPS. G, H, I) Scatterplots showing how specific QNP variables within correlation clusters 3 and 7 of the co-correlation matrix depicted in (C) relate to CPS. Dots represent values from each donor in the cortical layer indicated, lines are LOESS regressions for measurements across donors within each layer. G, H) Cluster 3 is comprised of variables increasing along pseudo-progression, such as the number of AT8 positive (pos) cells per unit area, 6e10 positive objects per unit area, or the average 6e10 positive object diameter of the 6e10-ir Aβ plaques. (I) Cluster 7 comprised variables decreasing their value along CPS, such as the number of NeuN positive cells or percent NeuN-ir cell area. The heatmap on each QNP variable across layers represents the p-value from a general additive model in which pseudo-progression was binned into 5 equal intervals.

The number of Aβ plaques and pTau+ neurofibrillary tangle bearing neurons in each donor were consistent with traditional staging thresholds for Braak stage and Thal phases, respectively (**Extended Data Fig. 3b**). However, at higher Braak stages and Thal phases we observed high variability in pathological burden that underscored the limitation of classical staging (as has also been observed with biochemical methods^75^) (**Extended Data Fig. 3c**). pTDP-43 and α-Syn pathologies were detected in the relatively small number of donors with high stage LATE-NC^76^ and neocortical LBD, respectively (**Extended Data Fig. 3d**). Cross-correlation of the QNP variables followed by hierarchical clustering revealed 8 biologically coherent clusters (**Fig. 2c**), with 2 anti-correlated clusters: cluster 3, which contained measurements of AD-related pathological proteins (i.e. diameter of Aβ plaques, number of Aβ plaques or pTau-bearing cells) and cluster 7, which contained NeuN-ir in neurons related variables (i.e. number of NeuN-ir nuclei per area).

We were inspired by biophysical studies^77^ that suggested that pathology aggregates exponentially in AD to construct a Bayesian model to infer AD pathological burden from the trajectory of each QNP variable. The models assigned a continuous pseudo-progression score (CPS) to each donor that ranged from 0 to 1 (**Extended Data Fig. 4a**), accounting for measurement noise and inferring the posterior distribution of all parameters with exponential dynamics. Along CPS, the number of pathological pTau-bearing neurons and Aβ plaques increased exponentially across donors (**Fig. 2d**, **Extended Data Fig. 4b**), with only small increases early in CPS and larger increases later. There was no clear relationship to pTDP-43^78^ and α-Syn levels. Consistent with the anti-correlation across QNP variables mentioned before, the number of NeuN-ir nuclei decreased along CPS, but had linear dynamics, suggesting some neuronal loss may precede large-scale plaque deposition and neurofibrillary tangle formation. Later in CPS, in donors with the highest pathological burden, we observed increased cellularity (i.e. the number of nuclei detected per area) and increased number of GFAP-positive nuclei (**Fig. 2e**, **Extended Data Fig. 4b**), consistent with later-stage astrogliosis. Importantly, CPS correlated with independent clinical data not included in the model, comprising Braak stage, Thal phase, ADNC score, and cognitive scores (CASI), but not other covariates, such as age (**Fig. 2f**, **Extended Data Fig. 4c**).

The CPS score allowed us to revisit the complex cross-correlation structure seen in QNP variables and understand their dynamics (**Fig. 2c**). We divided CPS into 5 equal bins and determined whether significant changes occurred in each with a generalized additive model (GAM). Cluster 3 encompassed several variables related to plaque and tangle pathology that mostly had their first significant increases later in CPS (**Fig. 2g**). Specifically, CPS=0.4 to 0.6 (bins 2 and 3) appeared to be a critical point when pTau-bearing cells and Aβ plaques started accumulating more substantially. It was also the point in CPS when donors started exhibiting increasing cognitive deficits. Within cluster 3, the diameter of Aβ plaques increased much earlier than other variables (**Fig. 2h**), having a significant change starting at CPS=0.2 (bin 1). This suggested that while plaque number was still low in early CPS, other Aβ species such as peptides and oligomers may be present. NeuN immunoreactivity decreased significantly along CPS in all cortical layers (**Fig. 2i**). Further, we observed an interaction between clusters 1 and 3 (**Fig. 2c**, blue box, **Extended Data Fig. 4d**) that captures the accumulation and colocalization of pTDP43-inclusions in pTau-bearing cells, as previously described^79^. Most remaining variables displayed significant increases after CPS bin 3 (**Extended Data Fig. 4e**). These observations illustrate that CPS incorporates pathologic burdens and cytologic changes to effectively capture AD severity in a continuous quantitative metric. Two epochs of AD emerge along CPS: (1) one early epoch where donors have low levels of plaque and tangle pathology and are cognitive unaffected but do have some neuronal loss and evidence of early amyloid pathology and (2) one late epoch where donors have markedly increased levels of AD pathology, neuronal loss, and cognitive impairment.

### Constructing an integrated, multi-modal AD atlas in MTG

Prior BICCN efforts identified 151 transcriptionally distinct cell types and states in the MTG from young, neurotypical reference donors^80^, hierarchically organized into 24 highly separable subclasses (e.g., L2/3 intratelencephalic-projecting excitatory neurons or L2/3 IT) within 3 main classes (excitatory neurons, inhibitory neurons, and non-neuronal cells). We used this BICCN reference as a base to construct a cellular taxonomy for SEA-AD. In order to map SEA-AD data to cell types consistently across all 84 donors, we first defined robust transcriptional “supertypes” in the BICCN reference. These supertypes represent the cell types that could be reliably re-identified in reference data sets (mean F1 score=0.91) using hierarchical probabilistic Bayesian mapping methods^81–83^(**Extended Data Fig. 5a, b**). Cell types not re-identified in reference data were unlikely to be consistently mapped in SEA-AD data. The cellular resolution of supertypes surpasses the level of analysis traditionally conducted in AD research, which typically analyzes cellular and molecular changes at our subclass level. We then mapped SEA-AD snRNAseq and snMultiome nuclei to the 125 defined supertypes using the same mapping method above (**Fig. 3a**). After removing low quality nuclei (**Extended Data Fig. 5c**), we noted some non-neuronal nuclei that had systematically lower mapping scores. This suggested cell types or states may be present in the SEA-AD dataset that were not captured in the original BICCN reference. We used a clustering-based approach to identify and add 14 non-neuronal cell types or states to our final SEA-AD taxonomy, resulting in 139 supertypes (**Fig. 3a**, **Extended Data Fig. 5d,e; Methods**). DLPFC snRNAseq data from the same SEA-AD donors was mapped to a matched DLPFC BRAIN Initiative cellular taxonomy using identical methods. We then extended our transcriptionally defined supertypes across snRNA-seq, snATAC-seq, and snMultiome datasets to construct a joint multiomic representation^84^ from both neurotypical reference and diseased donors (**Extended Data Fig. 6a-e**).

**Figure 3:**
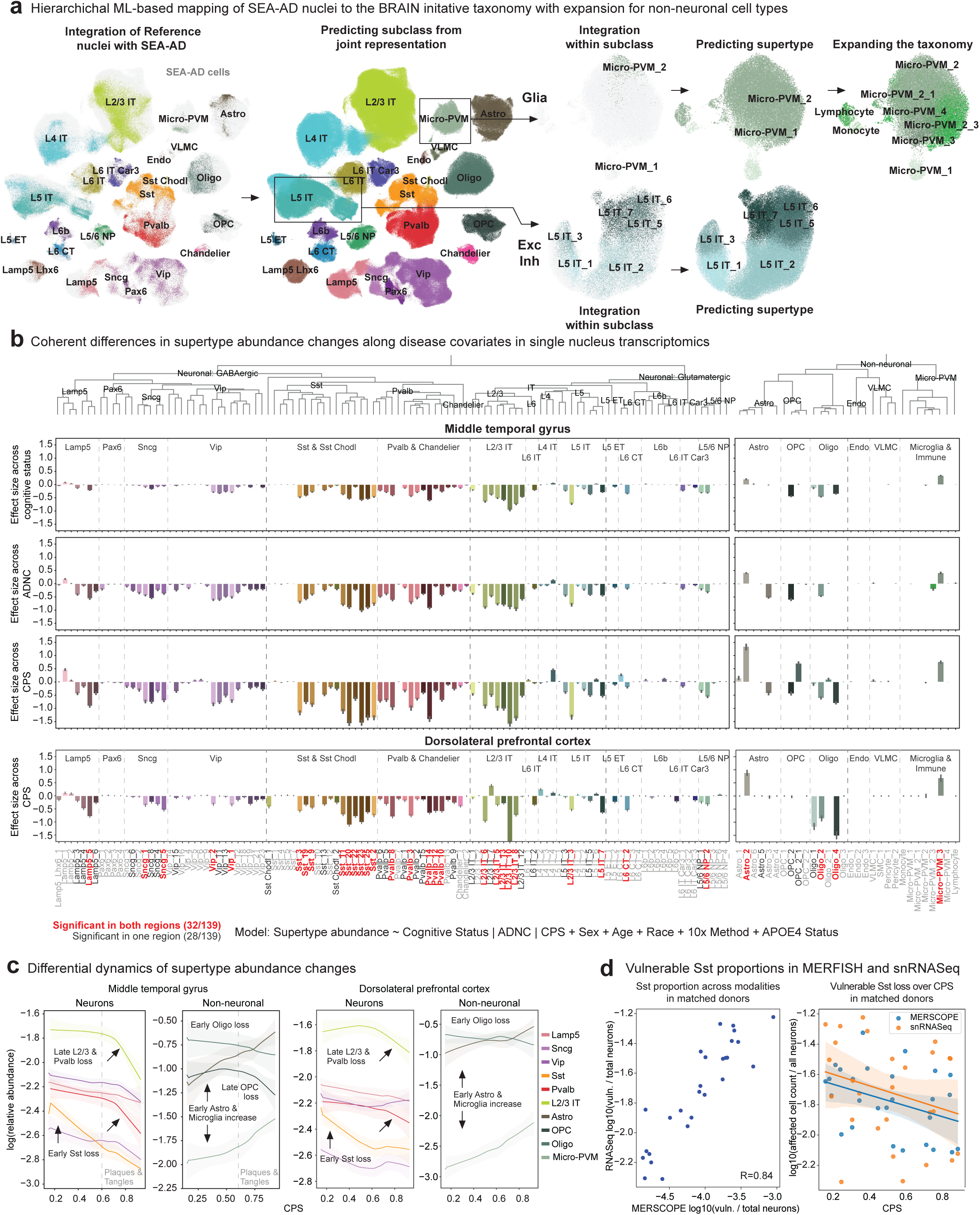
Vulnerable Populations in MTG concentrate around superficial supragranular layers. A) Schematic showing the hierarchical mapping procedure utilized to create SEA-AD taxonomy and annotate all SEA-AD cells. Reference MTG cells were used to define neuronal supertypes (see **Methods**). Then, reference and SEA-AD data sets were integrated, and cells were assigned to subclasses by using deep generative models (DGMs) of gene expression. Neuronal supertypes were annotated by subsetting the data to each neuronal subclass and using a new subclass-specific DGM for label transfer. Non-neuronal subclasses were annotated with the same procedure, but also underwent robust Leiden clustering (see **Methods**) to identify transcriptionally distinct populations not present in the reference taxonomy. Each scatterplot shows the UMAP coordinates computed from the nearest neighbor graph based on the latent representations from the DGMs for either all reference and SEA-AD nuclei or only those in the subclasses indicated from the middle temporal gyrus (MTG). In the plots focused on integration SEA-AD nuclei are colored light grey. Cell subclasses and supertypes are indicated. B) Barplots showing effect sizes for how each supertype changed in its relative abundance in the MTG across (Top) cognitive status, (Middle) ADNC, or (Bottom) continuous pseudo-progression (CPS) from the compositional model that also controlled for sex, age, single-cell technology, and *APOE4* status. Negative/positive values indicate that populations are decreasing/increasing across each covariate under analysis. Note, highly consistent changes across disease covariates with the greatest effect size along CPS. Below is an identical plot showing effect sizes for how each supertype changed in the DLPFC across CPS, controlling for sex, age at death, and race. Supertypes are indicated below all plots. Red, significantly changed in both cortical regions (MTG and DLPFC); dark grey significantly in one cortical region; light grey, not significantly changed in either brain region. Note, 26 of the 28 supertypes that significantly changed in only one brain region were specific to the earlier affected MTG. Subclasses are indicated in the top and bottom plots, with light grey dashed lines separating subclasses in the same cellular neighborhood and darker grey lines separating cellular neighborhoods. Supertypes are ordered by the dendrogram at top that shows how transcriptionally similar they are to one another. C) LOESS regression relating the log-normalized relative abundance (within all neuronal or all non-neuronal nuclei) of supertypes that were significantly changed in the MTG (left two plots) or DLPFC (right two plots) to the continuous pseudo-progression score (CPS). Supertypes were grouped by their subclasses to facilitate visualization of how each set of supertypes changed. Note, Sst supertypes decrease in their relative abundance early in CPS, before an exponential increase in the number of plaques and tangles present (indicated on each plot with a dashed light grey line). In contrast, L2/3 IT and Pvalb supertypes decrease as AD pathology increases. Uncertainty in each line represents the standard error from 1000 LOESS fits with 80% of the data randomly selected in each iteration. D) Left, scatterplot showing the correlation of vulnerable Sst supertype’s relative abundance in snRNA-seq and MERFISH data from matched donors (R=0.84). Right, scatterplot relating the relative abundance of vulnerable Sst supertypes to CPS in the snRNA-seq (orange) and MERFISH (blue) datasets from the same donors. Linear regressions with similar slopes are shown for both modalities.

To define the spatial distribution of supertypes in the MTG across AD and to enable orthogonal validation of cellular changes in matched SEA-AD donors, we generated a large-scale, cellularly resolved spatial transcriptomic dataset that contained 69 sections from a subset of SEA-AD donors, sampled across levels of AD pathology (n=28, **Extended Data Fig. 7a**). Tissue sections were profiled using a 140 gene panel (**Supplementary Table 3**) designed to capture a published MTG reference taxonomy^30^ and using a highly reproducible MERFISH-based data collection and analysis pipeline (**Extended Data Fig. 7b**). High quality data was obtained across disease severity, solving challenges present in spatial transcriptomic platforms profiling human tissue^85^, such as the removal of autofluorescence artifacts that are exacerbated with age and disease. To QC our spatial data, we compared spatial transcript counts across sections to bulk RNA-seq from brain samples from a subset of donors (mean correlation=0.62, **Extended Data Fig. 7c**), and correlated transcript counts across whole tissue sections with those within segmented cells (mean correlation=0.85, **Extended Data Fig. 7d**). This high correlation across quality control metrics was also present when assessing within donor technical reproducibility (mean correlation=0.98, **Extended Data Fig. 7e**). Next, we benchmarked supertype mapping ability using only the 140 gene panel in reference snRNA-seq data. Subclasses were readily identified (with F1 scores uniformly near 1, **Extended Data Fig. 7f**) and supertype mapping accuracy decreased slightly (134 of 139 with an F1 score above 0.7). The MERFISH gene panel failed to resolve 5 non-neuronal types absent in the original MTG reference. After mapping each cell in the spatial transcriptomic dataset to subclasses and supertypes, we found concordance between expected and mapped spatial distributions; for example, excitatory IT subclasses were restricted to expected cortical layers, and matched proportions observed in previous studies of neurotypical MTG tissue^80,86^ (**Extended Data Fig. 7g, h**). There was also high qualitative correspondence in gene expression across subclasses between donor matched snRNA-seq and MERFISH data (**Extended Data Fig. 7i**). The consistency seen across quality control and mapping metrics provided confidence in the application of our MERFISH platform across SEA-AD donors, even in those with higher levels of pathology.

### Vulnerable and disease associated supertypes

Inspired by decades of observations in the field^12,87–95^, we first asked whether specific supertypes are changed in their relative abundance through AD progression. We used scCODA, a Bayesian method^96^ that accounts for the compositional nature of relative abundances (e.g. if one cell type increases all others decrease in relative abundance), to test for changes across cognitive status, ADNC and CPS in the MTG snRNA-seq, snATAC-seq, and snMultiome datasets. This analysis was conducted separately for neuronal and non-neuronal cells as they are sorted at different ratios (70% neurons, 30% non-neurons). Using this approach, we identified multiple neuronal and non-neuronal supertypes that decrease in relative abundance as a function of disease severity, while a few highly specific non-neuronal supertypes increased **(Fig. 3b)**. Furthermore, a similar pattern of supertype abundance changes is seen for all three metrics of disease, with 36 of 139 (26%) supertypes significantly affected (mean inclusion probability > 0.8) across each disease-related covariate. While there was overlap in affected supertypes across disease-related covariates, effect sizes across CPS were greater. The number and effect size of affected supertypes were significantly less in other covariates and we observed consistent results with and without the SA donors and in other single nucleus data modalities (**Extended Data Fig. 8a,b, Supplementary Table 4**).

The extensive annotation of the BICCN reference (which SEA-AD is built upon) enabled meaningful interpretation of the types of cells affected in AD. We found that only a subset of supertypes were affected from most subclasses, highlighting the necessity of analyzing transcriptomic datasets at greater cellular resolution. The vulnerable neuronal supertypes (defined as those with statistically significant proportion decreases) include a subset of excitatory intratelencephalic (IT) neuron types largely in layer 2/3 (L2/3 IT), a subset of GABAergic interneuron types derived from the medial ganglionic eminence (MGE; Sst and Pvalb) and caudal ganglionic eminence (CGE; Vip, Lamp5 and Sncg) **(Fig. 3b**, left). Among non-neuronal affected populations, we observed increases in one microglial and one astrocytic supertype and decreases in one oligodendrocyte and one OPC supertype **(Fig. 3b**, right). We related the loss of vulnerable neurons and emergence of disease-associated non-neuronal states to the neuropathological changes (noted above) using CPS. Somatostatin inhibitory (Sst) interneurons and oligodendrocytes supertypes decreased early and continuously with CPS, with their changes occurring concurrent with the early increase in plaque size but before the exponential ramp in the number of plaques and tangles, followed by microglial and astrocyte supertypes increases (**Fig. 3c**, left). Multiple affected neuronal subclasses decreased later in CPS, concurrent with the exponential increment in plaque and tangle pathology. L2/3 IT neurons and Pvalb interneurons (which are common synaptic partners) decrease sharply together at high CPS.

To determine whether these cellular changes were replicated in other cortical areas, we tested for abundance changes with CPS in our snRNA-seq dataset from DLPFC in identical donors. DLPFC has a lower pathological burden than MTG at the same disease stage, so we expected both the number and effect sizes of affected types to be lower. More than half the supertypes affected in MTG were also changed in DLPFC (32/58), with these supertypes accounting for nearly all those undergoing changes in DLPFC (32/34) (**Fig. 3b**). When we related the dynamics of each supertypes change in the DLPFC with CPS they showed remarkable similarity across regions (**Fig. 3c**, left). Finally, we used our spatial transcriptomics dataset, collected from a subset of SEA-AD donors, to investigate changes in neuronal populations affected in AD, corroborating the vulnerability of specific supertypes. The relative abundance of affected Sst neurons were highly correlated (correlation = 0.84) between snRNAseq and MERFISH datasets (using 28 donors for which both modalities were profiled, **Fig. 3d**, left). Together, these analyses revealed a consistent, early, and continuous decline in these Sst supertypes across modalities (**Fig. 3d**, right) and demonstrate that there is a robust and consistent cellular signature of AD severity that involves selective cell populations affected differentially over disease progression.

### An integrated atlas of community AD data

Previous studies have described AD-associated molecular and cellular changes ^9–11,13–18,97–99^, but diversity in cohort selection, experimental design, and data processing techniques have made cross-study comparisons challenging. To further replicate our results using multiple studies, we obtained snRNA-seq data and associated donor metadata from DLPFC from 10 additional AD studies spanning 707 donors^9–11,13–17,97,98^. Harmonizing metadata across donors enabled direct cohort comparison, revealing that most studies span the spectrum of plaque and tangle pathology (**Fig. 4a**, top and **Extended Data Fig. 9a**). SEA-AD contained a similar distribution of donors with respect to neuritic plaques but a greater fraction of donors with larger neurofibrillary tangle spread (Braak stages V and VI). Donors in these Braak stages have neurofibrillary tangle pathology that extends to the DLPFC (as well as other cortical areas outside of the temporal pole), making them critical to understanding AD progression in these regions. Also notable, with rare exceptions^11^, the fraction of donors in each cohort with an *APOE4* allele, clinically diagnosed dementia, and severe co-morbidities were similar (**Fig. 4a**, bottom and **Extended Data Fig. 9a**).

**Figure 4:**
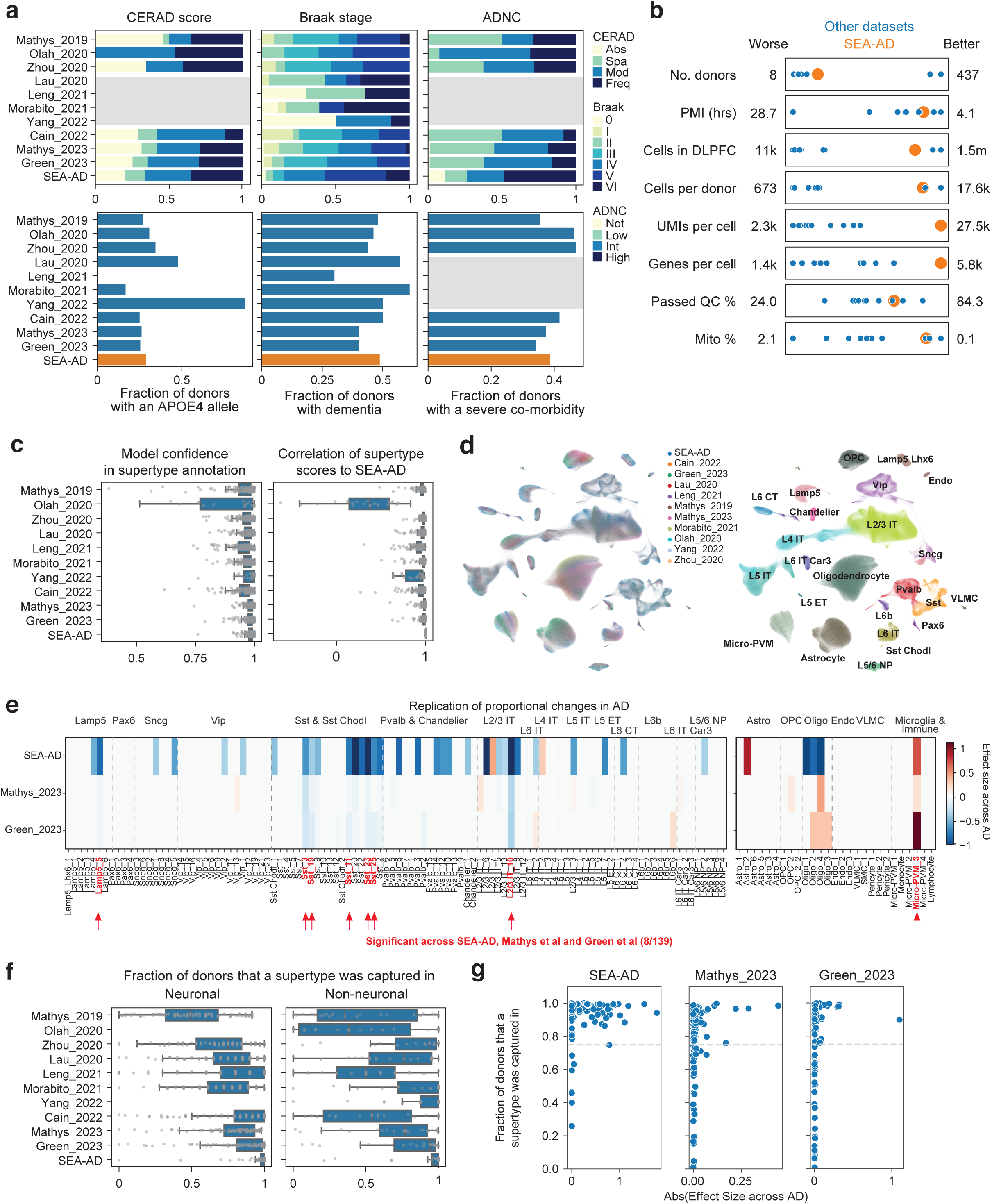
DLPFC single nucleus data integration replicates MTG vulnerable populations with AD. A) Barplots showing the fraction of donors in each of the publicly available snRNA-seq datasets that we harmonized metadata for and integrated classified in neuropathological stages (Top) or possessing *APOE4* alleles, dementia or a severe co-morbidity (Bottom). Grey boxes, metadata that was unavailable. Neuropathological staging included CERAD score for the distribution of neuritic plaques (ranging from Absent (Abs) to Sparse (Spa) to Moderate (Mod) to Frequent (Freq), Braak stage for the distribution of neurofibrillary tangles (ranging from Braak 0 to Braak VI), and the NIA-AA composite score AD Neuropathological Change (ADNC) (ranging from Not AD (Not) to Low to Intermediate (Int) to High). SEA-AD in indicated at bottom, in orange for variables that had only one category. All datasets applied snRNA-seq to the dorsolateral prefrontal cortex (or immediately adjacent region) in human donors that contained sporadic AD cases. B) Scatterplots showing the relative study size, dataset depth, and quality control metrics across publicly available snRNA-seq datasets (shown as blue dots) and SEA-AD (shown as a larger orange dot). The worst value for each metric is indicated on the left and the best value indicated on the right. Note, SEA-AD is the only study to perform consistently well (3^rd^ or better of the 11 datasets) across all metrics. C) Left, box and whisker plot showing the mapping confidence across datasets for each supertype from the deep generative model (DGM) that was used to annotate the publicly available snRNA-seq datasets to the SEA-AD cellular taxonomy. Right, box and whisker plot showing the spearman correlation of each supertype’s signature score across all nuclei in each dataset compared to SEA-AD (e.g. highly similar Sst_20, Sst_22, Sst_25, Sst_23 and Sst_11 supertypes would all have a high Sst_25 signature score, but the order from highest to lowest should be retained across datasets if they were mapped consistently). By definition SEA-AD supertypes all have a correlation score of 1. Note, consistently high model confidence and spearman correlations across datasets, except for Olah et al. D) Scatterplot showing the UMAP coordinates computed from the integrated latent representation of cells and nuclei from SEA-AD’s snRNAseq dataset on the DLPFC and each publicly available dataset color-coded by data set of origin (left) or subclass (right). E) Heatmap comparing the effect size of the relative abundance change of each supertype in the DLPFC across CPS (for SEA-AD) or ADNC (for Mathys_2023 and Green_2023) from a compositional model that also controlled for sex, age at death, and race in SEA-AD or sex, age, and *APOE4* status in Mathys_2023 and Green_2023 studies. Red text and arrows, supertypes that were significantly changed in abundance across all 3 studies. Note, some supertypes that were not significant in Mathys_2023 or Green_2023 had trend level changes similar to SEA-AD (e.g. Sst_20). Subclasses are indicated at the top of the heatmap, with light grey dashed lines separating subclasses in the same cellular neighborhood and darker grey lines separating cellular neighborhoods. F) Box and whisker plots showing the fraction of donors that each supertype was captured in across all 11 integrated datasets. Note, SEA-AD was the most deeply profiled dataset in the field with nearly every supertype being captured across all donors. No neuronal supertypes were identified in the Olah_2020 or Yang_2022 datasets due to their study design. G) Scatterplots relating the effect size for each supertype was changed in AD to the fraction of donors for which the supertype was captured in across the indicated datasets. Note, no populations captured in less than 70% of profiled donors (dashed grey line) were detected as significant across all studies. This represented extremely few supertypes for SEA-AD, but nearly a third of supertypes in Mathys_2023 and Green_2023 (including many that were significantly changed in SEA-AD).

SEA-AD aimed to profile many nuclei per donor with deep sequencing of libraries to drive high gene detection. Common re-processing of raw sequencing reads and quality control enabled direct comparison of data quality in the combined 5.5 million cells/nuclei across studies. SEA-AD was successful in simultaneously profiling a relatively large number of donors, number of overall nuclei, and number of nuclei per donor, while also having high sequencing depth and gene detection per nucleus (**Fig. 4b**, **Extended Data Fig. 9b**). To place these datasets into a single cellular nomenclature, we mapped them to the DLPFC BRAIN Initiative cellular taxonomy using the same hierarchical approach as above and computed marker-based signature scores for each supertype in each dataset (**Extended Data Fig. 9c**). In all but 1 dataset (Olah et al), both model confidences and supertype signature scores were uniformly high across types (**Fig. 4c**), indicating strong consistency in mapping across datasets. We qualitatively visualized these results by constructing integrated representations across all cells and across cells in each cell type neighborhood (**Fig. 4d** and **Extended Data Fig. 9d**).

Next, we tested for changes in supertype abundance along ADNC in two publicly available studies (Mathys et al and Green et al) with enough donors to reveal statistically significant changes^96^. Of the 34 supertypes that were significantly changed in the DLPFC in SEA-AD along CPS, 8 were also changed in these studies (**Fig. 4e**, **Extended Data Figs. 8a, 9e, Supplementary Table 4**). This included 5 of the Sst interneuron supertypes and the 1 Microglia supertype that were changed early in CPS, as well as Lamp5 interneuron and L2/3 IT excitatory neuron supertypes that were decreased later. Only changes in oligodendrocytes had contradictory significant effect sizes (decreasing in both the MTG and DLPFC of SEA-AD and increasing in both Mathys et al and Green et al), which will require deeper collaboration and investigation to resolve and may be related to differences in tissue dissection. Effect sizes were consistently lower in these datasets, more so than could be explained by using ADNC versus CPS alone (compared to **Fig. 3b**). The difference may relate to both studies having a lower fraction of donors with DLPFC tangle pathology (i.e. Braak stages V and VI), but also to sampling fewer nuclei per donor. The reduced nuclei per donor limited capture of each supertype consistently across donors in both studies (roughly 30% of supertypes were missing in at least a quarter of donors compared to only 4% in SEA-AD) (**Fig. 4f**). Significant changes were only detected in supertypes present in at least 75% of donors across SEA-AD and both publicly available datasets (**Fig. 4g**), suggesting this sparsity was particularly detrimental. Notably, some of the 34 supertypes that were not replicated had non-significant effect sizes that were directionally consistent with SEA-AD, such as Sst_20.

### Gene expression dynamics across supertypes and AD pseudo-progression

Numerous studies have implicated changes in specific molecular processes and pathways with AD, including mitochondrial function^100–103^, lipid biosynthesis^104–106^, proteostasis^107–109^, intracellular trafficking^110–112^, inflammation^113–115^, and more. To study the dysregulation of these processes by disease in a supertype-specific fashion, we tested for gene expression changes along CPS across each of the 139 supertypes (**Extended Data Fig. 10a, Supplementary Tale 5**). We found that the number of genes with significantly altered expression per supertype ranged from roughly 6,000 (in highly abundant IT excitatory neurons) to 180 (Endothelial and VLMC) (**Fig. 5a**), the latter being close to the expected false discovery rate. Most of the changes called significant were decreases in expression and had relatively small effect sizes, though a handful of genes had dramatically larger changes (**Extended Data Fig. 10b,** left). There was modest correlation (Pearson=0.62) between the number of nuclei in a supertype and the number of genes called significant (**Extended Data Fig. 10b,** right), suggesting the noise inherent in snRNA-seq from zero inflation is the limiting power for less abundant supertypes. Next, to characterize the dynamic molecular changes occurring with disease, we divided CPS into the two phases of AD noted above: an earlier phase in which pathology accumulates slowly and linearly and corresponds with a preclinical stage of disease (most donors exhibit unimpaired cognitive performance), and a later phase in which pathology accumulates exponentially and cognitive performance starts its decline. Comparing the average effect sizes for how each gene changed along CPS earlier and later revealed complex temporal dynamics (**Fig. 5b**).

**Figure 5:**
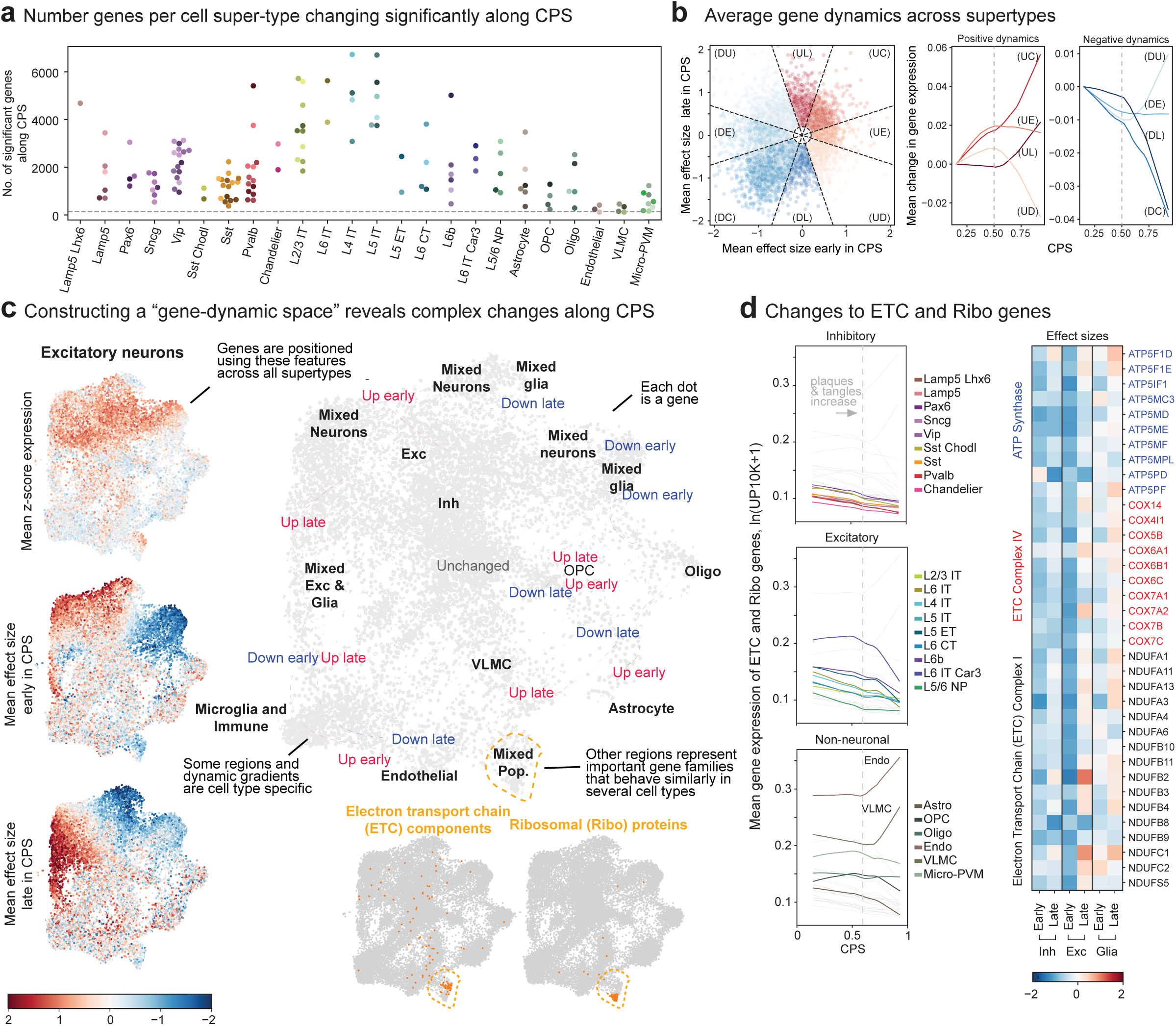
Gene expression changes along pseudo-progression exhibit complex cell type-specific dynamical patterns. A) Swarmplot showing the number of genes significantly changed with the continuous pseudo-progression score (CPS) in each supertype, organized by subclass. Grey dashed line, expected false discovery rate based on the average number of genes tested and the alpha threshold. B) Scatterplot relating the mean effect size across supertypes of each gene estimated using donors from the early versus late disease epochs along CPS. Genes were categorized into 8 bins given their early and late effect sizes: DU, down up. DE, down early. DC, down consistently. DL, down late. UD, up down. UE, up early. UC, up consistently. UL, up late. Right, LOESS regression relating the mean expression of all genes in each category to CPS. Note, the expected dynamics are observed in each gene set. Grey dashed line separates disease epochs (CPS=0.5) C) Framework to explore gene expression changes in an unsupervised manner. For each gene, early and late effect sizes and z-scored mean expression values were collected across supertypes. Next, an unsupervised low dimensional representation is built for all genes. Left, genes in the low dimensional representation are color coded by their mean expression values and early and late effect sizes in excitatory neurons as an example. Right, low dimensional representation of all genes qualitatively annotated to show areas of genes with cell type specific expression (black labels) and gene expression dynamics with CPS (blue to red labels and dashed lines). Right and bottom, region of the gene dynamic representation where electron transport chain (ETC) and ribosomal (Ribo) genes are clustered together. D) Left, LOESS regression relating the mean expression of electron transport chain (ETC) and ribosomal (Ribo)genes to CPS, color coded by inhibitory (top), excitatory (middle), and non-neuronal (bottom) subclasses. Dashed grey line, point in CPS when plaque and tangle pathology is definitively increasing (CPS=0.6). Note, nearly every neuronal type exhibited down-regulation of these gene families in the early disease epoch. Right, heatmap displaying mean effect sizes across cell class for genes within the ATP synthase complex (blue) and complexes 1 (black) and 4 (red) from the electron transport chain.

To visualize the landscape of gene expression dynamics across cell types, including their “baseline” expression patterns, we constructed a “gene-dynamic space” encompassing each gene mean expression and earlier and later effect sizes across CPS in all supertypes (**Fig. 5c**, **Extended Data Fig. 10c**). The organization of the gene-dynamic space suggested that many genes changed most strongly in the cell types they were specifically expressed in, with multiple dynamics along CPS. We discuss these cell type-specific changes further in the next three sections below. Other parts of the gene-dynamic space were more broadly expressed across cell subclasses and had similar dynamics across them. To better understand these regions, we curated 31 gene sets related to molecular processes implicated in AD (**Supplementary Table 6**) and determined if any were specifically enriched. One notable region contained nearly every nuclear encoded gene of complexes I (NADPH reductase) and IV (cytochrome oxidase) in the electron transport chain (ETC) as well as several ribosomal genes (**Fig. 5c**, bottom). Remarkably, across nearly every type of neuron these genes decreased in expression along CPS, particularly before the buildup of plaque and tangle pathology (**Fig. 5d**). Such a broad response across neurons may represent a protective mechanism in an environment where production of reactive oxygen^100,116,117^ species is particularly detrimental.

### Vulnerable Sst neurons in early AD

While L2/3 IT excitatory neurons are known to develop neurofibrillary tangles and are selectively lost in AD^73,94,118–120^, loss of other neuronal populations has been less studied until recently^9,9,16,17,121^. It remains unclear when in AD progression vulnerable interneuron populations are lost and characterization of their molecular identities, morphologies, tissue locations, and electrophysiological properties are incomplete. Here we focused specifically on vulnerable medial ganglionic eminence (MGE)-derived interneurons, as they encompass the Sst interneuron supertypes that were the earliest and most consistently vulnerable group of neurons across datasets. MGE interneurons form a transcriptional continuum that includes two major subclasses: Sst-positive and Pvalb-positive interneurons (**Fig. 6a**, left). Within this continuum, vulnerable Sst and Pvalb supertypes were as transcriptionally similar to each other as they were to other, unaffected supertypes within their respective subclasses (**Fig. 6a**, right and **Extended Data Fig. 11a**). As such, hundreds of genes were expressed in both vulnerable Sst and Pvalb supertypes but not unaffected supertypes from these subclasses (**Extended Data Fig. 11b, Supplementary Table 7**). Further, while Sst and Pvalb neurons are found throughout all layers of the cortex, vulnerable supertypes from both subclasses localized primarily to supragranular layers 2 and 3 in our spatial transcriptomics dataset (**Fig. 6b,c**). All Caudal ganglionic eminence (CGE)-derived interneurons (e.g. Lamp5-, Vip-, Sncg-, and Pax6-positive neurons) and L2/3 IT neurons are also only found in upper, supragranular layers (**Fig. 6b**, **Extended Data Fig. 11c,d**) so nearly all of the vulnerable supertypes that we identified resided in upper cortical layers (**Fig. 6b**, **Extended Data Fig. 11c,d**).

**Figure 6:**
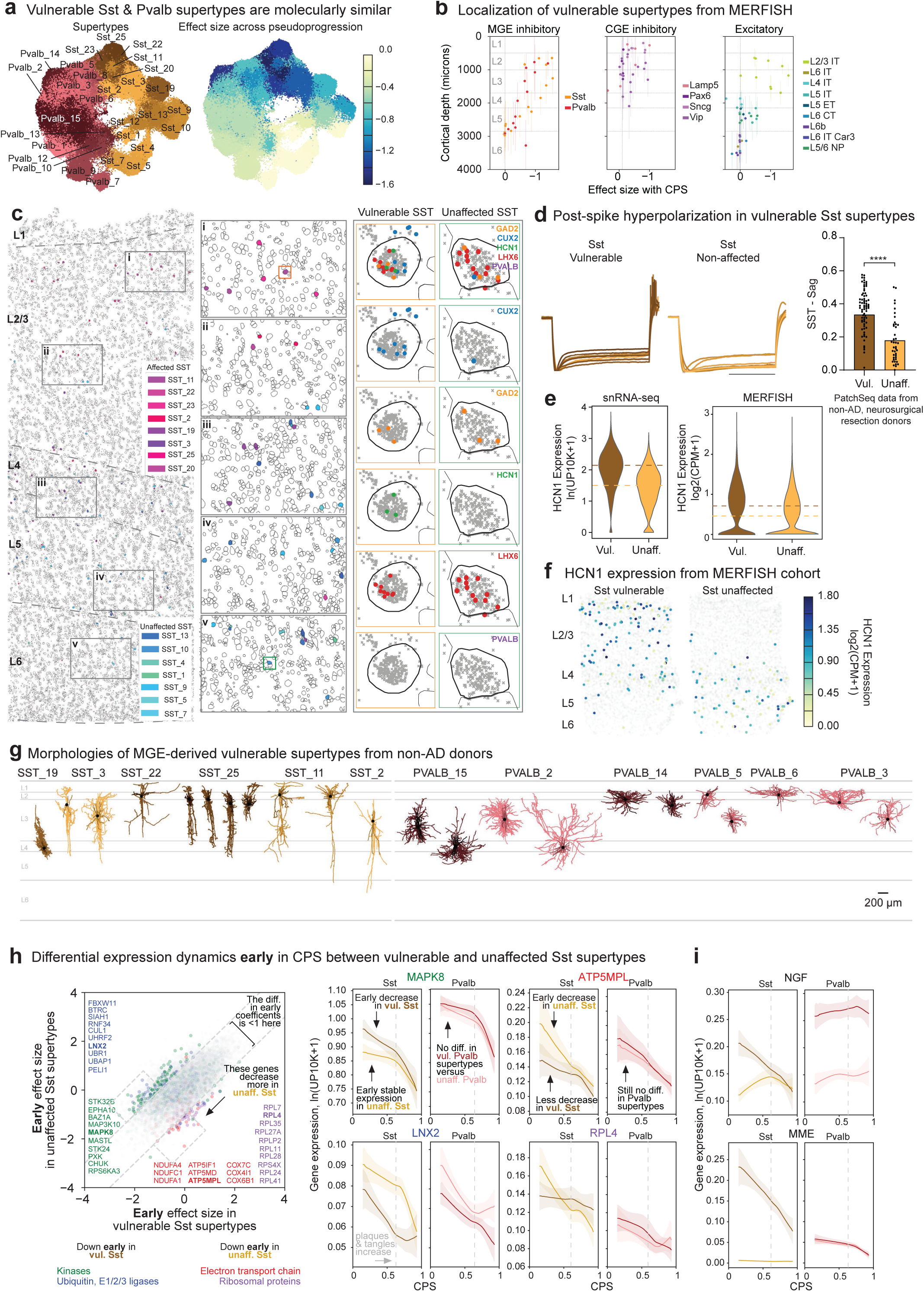
Superficial vulnerable MGE-derived inhibitory interneurons exhibit similar transcriptional profiles and common electrophysiological feature) A) Scatterplot showing the UMAP coordinates of MGE-derived neurons (Sst, Pvalb) from the middle temporal gyrus (MTG) color coded by supertype (left), or effect size of relative abundance changes in each supertype along CPS from scCODA (right). Note, the most strongly affected supertypes in both Sst and Pvalb subclasses are close together. B) Scatterplots relating the effect size of relative abundance changes of each supertype in the MTG from scCODA (colored by subclass) to their cortical depth from MERFISH experiments organized into CGE-, MGE-derived inhibitory neurons or excitatory neurons. Note, larger effect sizes are found for supertypes in superficial cortical layers (indicated and separated by horizontal grey lines) of MTG regardless of developmental origin. Each point represents the mean depth of the supertype in the MERFISH dataset, with error bars representing the standard deviation. C) Left, MERFISH-profiled brain slice in early CPS donor (CPS=0.23) showing each cells location and boundaries defined by the cell segmentation, with cortical layers indicated (L1-L6) and separated by dashed grey lines. Vulnerable and unaffected Sst neurons are color-coded with pink-purple (vulnerable) and green-blue (unaffected), respectively. Note, Sst cells that decline significantly during AD progression tend to localize in cortical layers 2 and 3 (L2/3), whereas Sst cells located in deep layers do not decline in abundance with disease progression. Insets: i) Detail of Layer 2 Sst interneuron supertypes. Most Sst types in this layer are affected. Orange box indicates cell shown in detail on the right, bounded by orange border. ii) Detail of Layer 3 Sst interneuron supertypes. Most Sst types in this layer are affected. Iii) Both affected and unaffected Sst supertypes are localized to L4 in roughly even proportions. iv) Layer 5 detail indicates that most Sst supertypes in this layer are unaffected. V) Layer 6 detail indicates that most Sst Ssupertypes in this layer are unaffected. Green box indicates unaffected Sst cell shown in detail in M-R, bounded by green border. Right, Example expression pattern of key genes (shown as grey or colored dots) in a vulnerable (orange box) and unaffected (green box) Sst neuron. Sst neurons should be positive for GAD2 (orange) and LHX6 (red), but negative for Pvalb (purple). Neurons in L2/3 have enriched expression for CUX2 (blue) and HCN1 (green). D) Left, electrophysiological traces showing post-spike hyperpolarization of membrane potential (y-axis) over time in all vulnerable Sst neurons but not the vast majority of unaffected Sst neurons recorded from tissue of non-AD human donors that underwent surgical resection. Right, bar and swarmplot showing sag distributions in individual vulnerable (Vul) and unaffected (Unaff) Sst neurons. P-values for all differential electrophysiological features are in **Supplementary Table 8**. E) Violin plots showing *HCN1* expression in vulnerable (Vul) and unaffected Sst neurons in both snRNA-seq (left) and MERFISH (right) datasets. Note, *HCN1* expression is significantly higher in vulnerable Sst neurons in both datasets (p-values from NEBULA are near 0). Mean expression in each group and each dataset are noted by colored dashed lines. ln(UP10K+1), natural log of UMIs per 10,000 plus 1. log2(CPM+1), log base 2 of counts per million plus 1. F) Exemplar scatterplot showing the positions of all cells (as dot) in the cortical column an early CPS donor (CPS=0.23). Sst cells are colored by their *HCN1* expression, with superficial supertypes having higher expression on average. All other cells are colored light grey, cortical layers are indicated (L1-L6) G) Exemplar morphological reconstructions of vulnerable MGE-derived interneurons from PatchSeq data obtained from non-AD donors. Cell dendrites are colored by their supertype annotation (indicated above, cell somas are indicated with black dots, and cell axons are darkened versions of the supertype colors. Cortical layers are indicated (L1-L6). Scale bar, 200 um. H) Left, scatterplot relating the mean early effect size of each gene (dots) in vulnerable versus unaffected Sst supertypes. Gene families that decrease preferentially in vulnerable Sst supertypes are color-coded blue (ubiquitin ligases, p-value=0.036) and green (kinases, p-value=8.92e-11). Gene families that decrease preferentially in unaffected Sst supertypes are color-coded red (electron transport chain, p-value near 0) and purple (ribosomal proteins, p-value near 0). The vast majority of genes fall within dashed grey lines bounding the space where effect sizes are within 1 unit of each other. Exemplar genes from each family are indicated in the plot. Right, LOESS regression plots relating the mean expression of indicated genes from these families to CPS across vulnerable (dark orange) and unaffected (light orange) Sst and and vulnerable (dark red) and unaffected (light red) Pvalb supertypes. Uncertainty in each line represents the standard error from 1000 LOESS fits with 80% of the data randomly selected in each iteration. ln(UP10K+1), natural log UMIs per 10,000 plus 1. Dashed grey line, point in CPS when plaque and tangle pathology is definitively increasing (CPS=0.6). I) Same LOESS regression plots as in (H) for notable single gene examples (NGF and MME) that decrease preferentially in vulnerable Sst supertypes but which do not belong to the gene families indicated.

To describe the morpho-electrical properties of vulnerable MGE supertypes, we harnessed a recently released large-scale Patch-seq dataset profiling MTG interneurons in non-AD human donors that underwent surgical resection^38,39^ and remapped its cells to the SEA-AD taxonomy. Vulnerable Sst supertypes had higher post-spike hyperpolarization (Sag) and lower membrane polarization time constants (Tau) compared to unaffected Sst supertypes (**Fig. 6d**, **Extended Data Fig. 11e, Supplementary Table 8**), differences not seen between vulnerable and unaffected Pvalb supertypes (**Extended Data Fig. 11f,g**). HCN channel activity is involved in setting sag level and, consistent with this difference, we observed higher HCN1 expression in vulnerable Sst supertypes in both snRNA-seq and MERFISH (**Fig. 6c,e,f**). Finally, the morphological reconstructions of vulnerable Sst and Pvalb interneurons qualitatively confirmed their supragranular localization and demonstrated that, despite their molecular similarity, they span a wide morphological range that includes non-Martinotti cells, Sparse, Basket, Basket-like, and the Double Bouquet cells (**Fig. 6g**). This molecular, electrophysiological, anatomical, and morphological characterization provided the deepest characterization to date of selectively vulnerable interneurons, enabling their study across several different modalities in future studies. More importantly, this level of resolution now allows consideration of AD progression as selective circuit dysfunction, and the consequences on cortical function that would ensue from perturbation of specific circuit elements (as discussed below).

We compared the early molecular changes across vulnerable and unaffected Sst supertypes (**Fig. 6h**). While the vast majority of genes changed similarly in both groups, specific gene families were significantly differentially down-regulated. Most notably, the vulnerable Sst supertypes collectively down-regulated specific kinases (from the TK^122^ and CAMK^123^ families) (**Fig 6h**, green) and E3 ubiquitin ligases (from the HECT family^124^) (**Fig 6h**, blue). They did not, however, down-regulated components of ETC (**Fig 6h**, red) and ribosomal genes (**Fig. 6h**, purple) as every other neuronal supertype did (including unaffected Sst supertypes). In contrast, vulnerable and unaffected Pvalb supertypes had no gene families affected differentially between them early in CPS (**Extended Data Fig. 11h**). In addition to the differentially affected gene families, still other notable genes were differentially changed early in CPS, including sharp down-regulation of neuronal growth factor (*NGF*) and *MME* specifically in vulnerable Sst supertypes (**Fig. 6i**). The cognate receptor for NGF, NGFR, is expressed specifically in Oligodendrocytes and OPCs, suggesting partial disruption in communication with vulnerable Sst supertypes that may impact myelination^125–127^. *MME* encodes the protease Neprilysin that may break down Aβ peptides^128^, suggesting vulnerable Sst supertypes’ may lose a critical mechanism for preventing later Aβ plaque formation near them.

### Microglia and Astrocyte activation in early AD

The discovery of disease associated microglia (DAMs^88,97^) has energized research to understand and ultimately modulate protective (e.g. clearing plaques^129^) and detrimental (e.g. driving neuroinflammation^113,130^) microglia function in AD^131,132^. Diverse direct and modulatory inflammatory roles suggest that many molecular microglial subtypes may exist; several cellular taxonomies for myeloid immune cells in the brain have been proposed using healthy and disease snRNA-seq data in human^12,16,17,97,133,134^. These taxonomies generally agree on three major types of monocyte-lineage cells in brain: monocytes, CNS-associated macrophages (CAMs)/perivascular macrophages (PVM), and a heterogenous group of microglia, which has proven difficult to reconcile across taxonomies. Comparing SEA-AD microglia taxonomy (**Fig. 7a**) to the most comprehensive effort to identify microglia subtypes to date^17^, revealed strong agreement across studies (**Fig. 7b**, top). Most notably, both taxonomies contained disease associated types (Micro-PVM_3 in SEA-AD and Mic.12 and Mic.13 in Green et al (2023)^17^) that were consistently increased in abundance with AD across datasets and have a common molecular signature (**Fig. 7c**). Also, both studies identify homeostatic and proliferative types and two other subtypes with no functional data or tissue localization information, which is necessary for properly assigning their molecular functions. While SEA-AD’s taxonomy is more conservative in splitting subtypes (with many 1-to-many relationships to those with Green et al (2023)), the same transcriptional continuum is captured in both datasets. Other studies have been even more conservative, describing only homeostatic and proliferative subtypes, despite their datasets containing other subtypes (**Fig. 7b**, bottom) such as a DAM subtype. Further collaboration and integration between studies, and localizing/functionally testing each proposed subtype will help resolve the true extent of molecular states. Broad agreement across studies on the AD transcriptional landscape and on certain consensus subtypes will greatly aid efforts to understand the role of microglia in disease initiation and progression.

**Figure 7:**
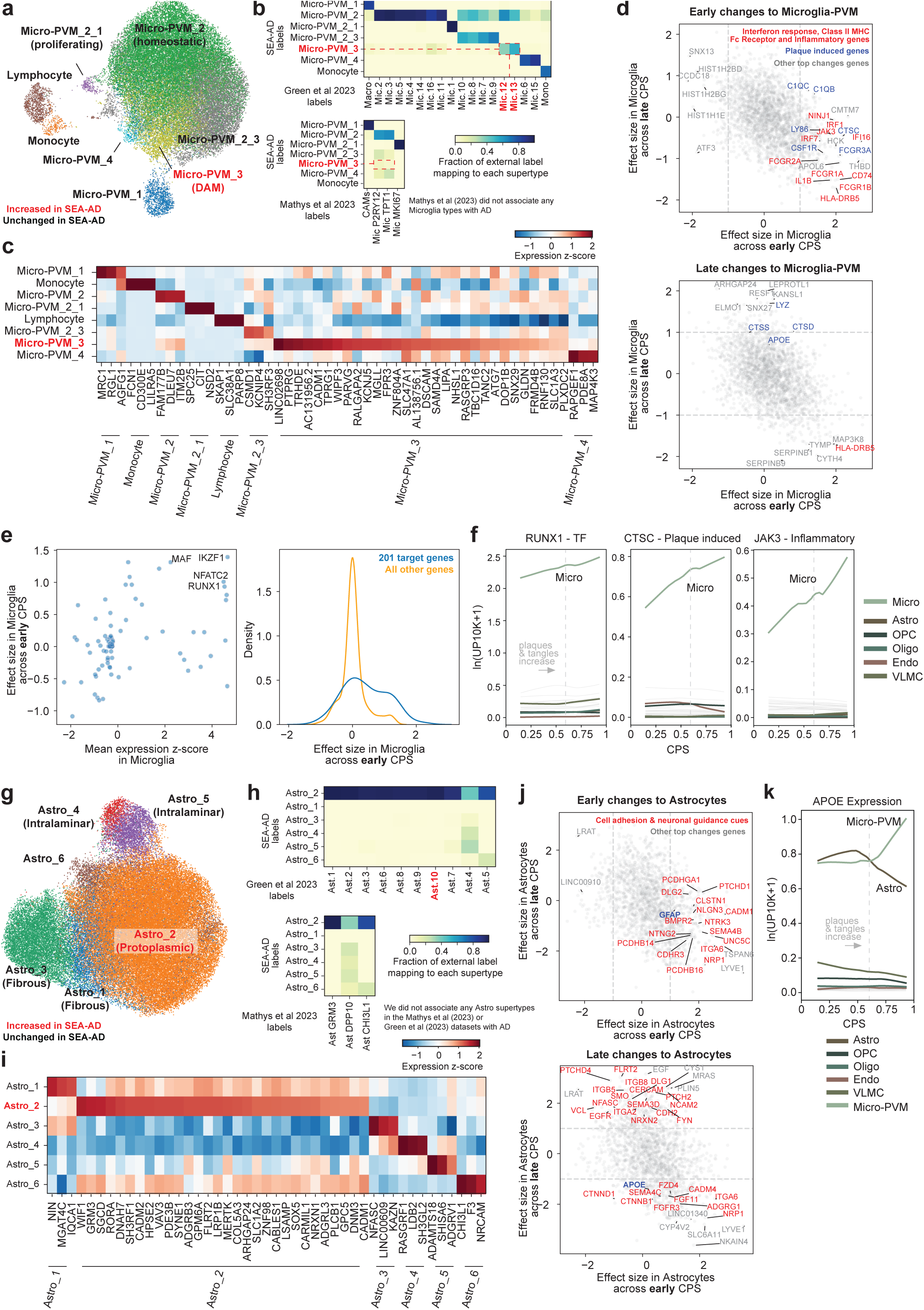
Early microglia and astrocyte activation and relationship of supertypes across publicly available data sets. A) Scatterplot showing UMAP coordinates for Micro-PVM and other immune supertypes from the middle temporal gyrus (MTG) SEA-AD dataset colored by supertype identity. Red text, disease associated microglia state. Other common states are indicated for supertypes (e.g. proliferating microglia). B) Heatmaps showing the confusion matrices comparing the annotations of microglia cells in the studies conducted by Mathys et al and Green et al 2023 with the annotations of the same cells labeled using the SEA-AD cellular taxonomy. Red text, SEA-AD supertypes that were significantly increased in AD in these datasets using scCODA. Also shows cell types that were associated with disease in the original studies. Note, Mathys_2023 did not associate a specific microglia cell type with disease, but we identified disease associated microglia in their dataset. The types called as disease associated in Green_2023 correspond to the disease associated SEA-AD supertype (dashed red lines). Also note, while some confusion exists, for the most part the taxonomies between Mathys_2023, Green_2023 and SEA-AD cover the same transcriptional landscape and more or less conservatively divide it. C) Heatmap showing the mean z-scored expression across microglia supertypes of marker genes identified by NEBULA. 3 marker genes are shown for all supertypes except for the disease associated Micro-PVM_3 (red text), which has 30 marker genes shown to establish correspondence with disease associated types in other datasets. D) Scatterplot relating the mean effect size of each gene across microglia supertypes in the early (x-axis) versus late (y-axis) epochs along CPS. Significant genes in the early phase (top) or late phase (bottom) that are interferon stimulated genes (p=0.040 early), class II major histocompatibility complex compnents (MHC) and Fc receptors (p=4.32e-6 early, p=9.86e-7 late), or pro-inflammatory genes are color-coded red, human plaque induced genes (p=1.22e-4 early) are color coded blue, and other top genes not in these families are color coded grey. The gene families indicated were significantly enriched among the up-regulated genes in the early and late disease epoch along. Grey dashed lines, denote effect sizes of 1 and -1 in the early (top) and late (bottom) AD epochs. E) Left, scatterplot relating the mean z-scored gene expression of transcription factors identified by the gene regulatory networks (GRNs) across non-neuronal cells in microglial supertypes versus their effect size in the early disease epoch along CPS. Comparing gene expression versus statistically significant effect size enabled the identification of candidate transcription factors (*MAF*, *IKZF1*, *NFATC2*, and *RUNX1*) that may be upstream of the microglia-specific early transcriptional response to AD pathology. Right, cumulative density plot depicting the effect sizes in the early disease epoch along CPS of genes downstream the transcription factors identified in (left) based on the GRNs (in blue) versus effect sizes all other genes (in yellow). Note, significant difference in mean effect size (p-value=1.36e-12 early) F) LOESS regression plots relating the mean expression of indicated genes from families noted in (D) to CPS across non-neuronal supertypes organized and colored by subclass. ln(UP10K+1), natural log UMIs per 10,000 plus 1. Dashed grey line, point in CPS when plaque and tangle pathology is definitively increasing (CPS=0.6) G) Scatterplot showing UMAP coordinates for Astrocyte supertypes from the middle temporal gyrus (MTG) SEA-AD dataset colored by supertype identity. Red text, disease associated protoplasmic astrocyte supertype. Other astrocyte supertypes are associated with their canonical morphological type (e.g. intralaminar in cortical layers 1 and 2 and fibrous in deeper cortical layers). H) Heatmaps showing the confusion matrices comparing the annotations of astrocyte cells in the studies conducted by Mathys et al and Green et al 2023 with the annotations of the same cells annotated with the SEA-AD cellular taxonomy. Note, Green et al associated Ast.10 with disease, which maps to the protoplasmic Astro_2 supertype in SEA-AD. We associated this type with disease in our MTG and DLPFC datasets, but not Green_2023. Mathys_2023 did not associate a specific astrocyte cell type with disease. Also note, most cell types in Mathys_2023 and Green_2023 correspond to protoplasmic and intralaminar astrocytes. The marker genes they use in their studies indicate they may not have captured a significant number of fibrous astrocytes, so are missing this cell type in their taxonomies. I) Heatmap showing the mean z-scored expression across astrocyte supertypes of marker genes identified by NEBULA. 3 marker genes are shown for all supertypes except for the disease associated Astro_2 (red text), which has 30 marker genes shown. J) Scatterplot relating the mean effect size of each gene across astrocyte supertypes in the early (x-axis) versus late (y-axis) epochs along CPS. Significant genes in the early phase (top) or late phase (bottom) that are cellular adhesion and neuronal guidance cues are color-coded red and other top genes not in these families are color coded grey. Grey dashed lines, denote effect sizes of 1 and -1 in the early (top) and late (bottom) AD epochs. K) Same LOESS regression plots as in (F) for the strongly disease-associated APOE gene, which decreases in expression in Astrocytes and increases in expression in Microglia in the late disease epoch along CPS.

In addition to confirming DAM’s existence in the SEA-AD dataset, broader molecular changes in microglia along CPS were consistent with previous studies. Early changes included significant up-regulation of gene sets involved in inflammatory processes (*IL1B*, *CSF1R*, *STAB1*, *NINJ1*, *JAK3*)^135–139^, interferon response (*IRF1*, *IRF7*, *IFI16*), Fc receptors (*FCGR1A*, *FCGR1B*, *FCGR2A*, *FCGR3B*), major histocompatibility complex II components (*CD74*, *HLA-DRB5*), and complement components (*C1QA*, *C1QB*) (**Fig. 7d**, top, red). Surprisingly, we also observed early up-regulation of several homologs of genes induced by Aβ plaques in AD (*CSF1R, CTSC, C1QA, C1QB, LY86, FCGR3A*)^20^ (**Fig. 7d**, top, blue), suggesting Microglia in individuals with low levels of Aβ plaques and tau tangles may be responding to Aβ peptides or oligomers^140^, or another pathological factor. Together, these changes indicate a highly stimulatory extracellular environment for microglia, early in AD. Other plaque induced genes were up-regulated later in CPS in donors with higher levels of pathology (**Fig. 7d**, bottom, blue), including more cathepsins (*CTSD* and *CTSS*) that may facilitate Aβ clearance^141,142^, the gene encoding lysozyme (*LYZ*), and *APOE*, which is by far the most strongly associated genetic risk factor for AD^22^. To identify the transcription factors driving early up-regulation of pro-inflammatory and plaque-induced genes, we leveraged SEA-AD’s snATAC-seq data to construct microglial gene regulatory networks (GRNs). GRNs allowed us to identify relevant microglia transcription factors (TFs). Next, we filtered these TFs by the specificity in their expression in microglia and their dynamic changes (requiring early upregulation), and identified 4 (*RUNX1*^143^, *IKZF1*, *NFATC2, MAF*) that are specifically expressed in Microglia, and are upregulated early in CPS (**Fig. 7e**, left). These transcription factors are predicted to co-regulate 201 genes, including genes noted above (**Fig. 7e**, right, **f**), and may be critical targets for modulating microglia responses to pathology.

Like microglia, astrocytes have been ascribed diverse roles in AD pathophysiology^19,90–92^, which makes understanding their molecular subtypes crucial. The SEA-AD taxonomy encompasses interlaminar, protoplasmic, fibrous, and a yet to be described astrocyte supertype (**Fig. 7g**). In contrast (**Fig. 7h**, top), Green et al (2023) split protoplasmic astrocytes into several subtypes, grouped interlaminar astrocytes into one subtype, and has few fibrous astrocytes (**Fig. 7i**). In both our MTG and DLPFC datasets, protoplasmic astrocytes (Astro_2) specifically increased early in CPS. While we could not replicate this association in Green et al (2023) (or Mathys et al (2023)), their original manuscript noted an increase in one protoplasmic subtype (Ast.10) with AD. This suggests agreement that at least a subset of astrocytes is increased with disease. Mathys et al (2023) had the fewest types, with one subtype for protoplasmic astrocytes, one subtype for fibrous and interlaminar astrocytes together, and one unknown subtype that was also similar to a type we identified (**Fig. 7h**, bottom). While consensus around certain subtypes provides a foundation to reconcile these taxonomies, the larger difference in the overall transcriptional landscape suggests more work is needed to compare tissue sampling to ensure consistent capture of all histologically established types.

Next, we sought to describe the molecular changes occurring in astrocyte supertypes, again dividing CPS into early and late epochs. Early changes included upregulation of cellular adhesion molecules (*CADM1*, *CDRH3*, *PCDHGA1*, *PCDHB14*, *PCDHB16, CLSTN1*, *ITGA6, NEO1, ANOS1*) and neuronal guidance cues (*NLGN3*, *NTRK3*, *SEMA4B, NTNG2*), signaling receptors (*PTCHD1*, *NRP1, BMPR2, UNC5C*)^144^, and *GFAP*, a known hallmark of AD and astrogliosis^145^ (**Fig. 7j**, top). Later in CPS, astrocytes continue to up-regulate molecules involved in cellular adhesion, axonal guidance, and signaling receptors, including *NCAM2* and *CERCAM*, additional hedgehog signaling receptors (*PTCHD4*, *PTCH2, SMO*) and their downstream target transcription factor *GLI1*, and both the *EGF* ligand and its receptor *EGFR* (**Fig. 7j**, bottom). Astrocytes also down-regulated *APOE* (**Fig. 7k**) and several genes involved in other signaling pathways, such as the Wnt-receptor *FZD4* and the FGF ligand and receptor, *FGF11* and *FGFR3*. Collectively, these molecular changes suggest a highly stimulatory extracellular environment occurring in early in disease, even in donors with relatively low levels of pathology.

### Oligodendrocyte loss and remyelination by OPCs

Multiple studies have indicated that the dysfunction of oligodendrocytes and myelin breakdown may be early events in AD^146–153^, possibly due to altered cholesterol localization and transport^23^, making the investigation of this cell type and their progenitor pool, OPCs, critical in understanding AD etiology. Among oligodendrocytes, two supertypes (Oligo_2 and Oligo_4) were decreased early in both MTG and DLPFC (**Fig. 8a**); both supertypes are found throughout the cortical column in the BRAIN Initiative reference dataset^26^. *CNP* was expressed in both (albeit higher in Oligo_4) (**Fig. 8b**), suggesting they are myelinating oligodendrocytes. We also observed a late decrease in one OPC supertype (OPC_2), which is found across cortical layers 2 through 6. When comparing against publicly available data sets, SEA-AD oligodendrocytes and OPCs types largely agreed with the fine-grained types described in Green et al (2023), with most supertypes having one-to-one or one-to-many relationships (**Fig. 8c**). There were a handful of many-to-many relationships that may represent different boundaries in the same transcriptional landscape and will require localization/functional data to resolve.

**Figure 8:**
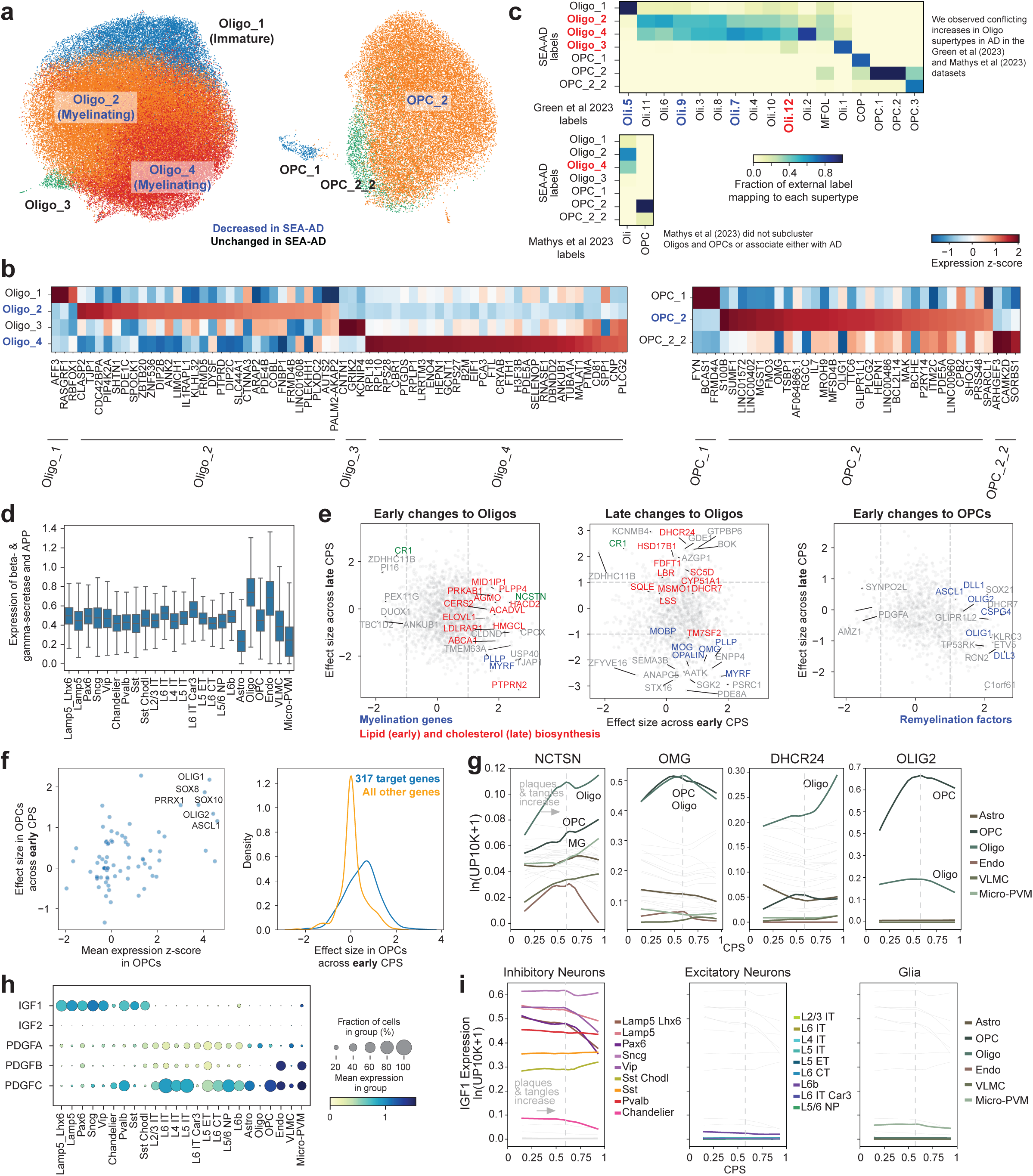
Early loss of oligodendrocytes with a re-myelination program in OPCs and relationship of supertypes across publicly available data sets. A) Scatterplots showing UMAP coordinates for Oligodendrocyte (left) and OPC (right) supertypes from the middle temporal gyrus (MTG) SEA-AD dataset colored by supertype identity. Blue text, vulnerable myelinating oligodendrocyte and OPC supertypes. An immature (non-myelinating) oligodendrocyte state is also indicated. B) Heatmap showing the mean z-scored expression across oligodendrocyte (left) and OPC (right) supertypes of marker genes identified by NEBULA. 3 marker genes are shown for all supertypes except for the vulnerable Oligo_2, Oligo_4, and OPC_2 supertypes (blue text), which all have 30 marker genes shown. C) Heatmaps showing the confusion matrices comparing the annotations of oligodendrocyte and OPC cells in the studies conducted by Mathys et al and Green et al 2023 with the annotations of the same cells labeled using the SEA-AD cellular taxonomy. Red text, SEA-AD supertypes that were significantly increased in AD in these datasets using scCODA (in contrast to the decrease seen in the SEA-AD data for these supertypes). Red and blue text also notes cell types that were associated or vulnerable, respectively, with disease in the original studies. Note, Mathys_2023 did not associate a specific microglia cell type with disease. Also note, while some confusion exists, for the most part the taxonomies between Mathys_2023, Green_2023 and SEA-AD cover the same transcriptional landscape and more or less conservatively divide it. D) Box and whisker plot showing the mean expression (natural log UMIs per 10,000 plus 1) of beta (*BACE1* and *BACE2*) and gamma (*PSEN1*, *PSEN2*, *APH1A*, *NCSTN,* and *PSENEN*) secretase components and the *APP* gene organized by subclass. E) Scatterplot relating the mean effect size of each gene across oligodendrocyte (left and middle) and OPC (right) supertypes in the early (x-axis) versus late (y-axis) epochs along CPS. Significant genes in the early phase (left) or late phase (middle) in oligodendrocytes that are involved in fatty acid biosynthesis (left) of cholesterol biosynthesis (middle, p=0.0040 late) are color-coded red and that are myelin components (p=0.006 late) are color coded blue. Significant genes in the early phase (right) in OPCs that are part of the re-myelination program (p=9.62e-5 early) are color coded blue. Other strongly changed genes not in these families are color coded grey across panels. These gene families indicated were significantly enriched among the up-regulated or down-regulated genes in the early or late disease epoch along Grey dashed lines, denote effect sizes of 1 and -1. F) Left, scatterplot relating the mean z-scored gene expression of transcription factors identified by the gene regulatory networks (GRNs) across non-neuronal cells in OPC supertypes versus their effect size in the early disease epoch along CPS. Comparing gene expression versus statistically significant effect size enabled the identification of candidate transcription factors (*OLIG1*, *OLIG2*, *SOX10*, *SOX8*, *PRRX1*, and *ASCL1*) that may be upstream of the OPC-specific early transcriptional response to AD pathology. Right, cumulative density plot depicting the effect sizes in the early disease epoch along CPS of genes downstream the transcription factors identified in (left) based on the GRNs (in blue) versus effect sizes all other genes (in yellow). Note, significant difference in mean effect size (p-value=3.14e-23 early) G) LOESS regression plots relating the mean expression of indicated genes from families noted in (E) to CPS across non-neuronal supertypes organized and colored by subclass. ln(UP10K+1), natural log UMIs per 10,000 plus 1. Dashed grey line, point in CPS when plaque and tangle pathology is definitively increasing (CPS=0.6) H) Dotplot depicting mean gene expression and fraction of cells in each group with non-zero expression in the SEA-AD MTG dataset organized by subclasses for the genes indicated. Expression is natural log UMIs per 10,000 plus 1. I) LOESS regression relating the mean expression of *IGF1* to CPS, color coded by inhibitory (left), excitatory (middle), and non-neuronal (right) subclasses. Dashed grey line, point in CPS when plaque and tangle pathology is definitively increasing (CPS=0.6).

Oligodendrocytes were recently described as a critical promoter of Aβ synthesis based on gene expression of beta (*BACE1* and *BACE2*) and gamma secretase (*PSEN1*, *PSEN2*, *PSENEN*, *APH1A*, *NCSTN*) components^18^. The mean expression of these genes was replicated in SEA-AD data (**Fig. 8d**), with oligodendrocytes having the highest levels of both *APP* and *PSEN1*. Therefore, the early loss of oligodendrocytes may be attributed to these higher levels of Aβ molecules that have known cytotoxicity, even prior to formation of extracellular plaques. Additionally, multiple gene expression programs are changing dynamically. There is an early up-regulation of a gamma secretase component (*NCSTN*), the transcription factor *MYRF* that regulates myelination^154^, and a structural component of myelin itself (*PLLP*) (**Fig. 8e**, left, **g**). Also up-regulated were lipid biosynthetic enzyme and transporter genes, including those that form secondary messengers (*PLPP4*, *PTPRN2, PRKAB1*), participate in beta-oxidation and carnitine biosynthesis in the mitochondria (*CPT1B, HMGCL, ACADVL*), ceramide/sphingolipid biosynthesis (*HACD2*, *ELOVL1*, *CERS2*), and cholesterol transport (*ABCA1*, *LDLRAP1*). Importantly, significant increases in expression of the cholesterol biosynthetic gene family, a proposed key process in AD etiology ^23^ occurs later in CPS (*DHCR24, LBR, FDFT, HSD17B1, SC5D, CYP51A1, SQLE, and DHCR7*) (**Fig. 8e**, middle, **g**). Further, late in CPS there is a down-regulation of *MYRF* and several components of myelin and myelination (*MOBP, MOG, OMG, PLLP, OPALIN*). The late change in both gene sets suggests they may represent a reaction to pathology rather than an early driver of dysfunction.

In OPCs, there was early up-regulation of several transcription factors (*OLIG1*, *OLIG2*, *SOX10*, *SOX8*, *PRRX1*, *ASCL1*) and Notch ligands (*DLL1*, *DLL3*) known to regulate differentiation^155–161^ to oligodendrocytes following loss of surrounding oligodendrocytes (**Fig. 8e**, right, **g**). Due to the overwhelming number of transcription factors involved in differentiation that changed early, we queried our OPC-specific GRN and identified 317 genes downstream of these factors (**Fig. 8f**, left). These genes were also upregulated early (**Fig. 8f**, right), compared to all other genes, and were predominantly involved in OPC differentiation. These data suggest a novel early event in AD may be damage and loss of oligodendrocytes that triggers a robust differentiation and remyelination response from OPCs in early-stage individuals. To identify cell types that may regulate the remyelination process, we examined expression of two signaling pathways that are important for OPC differentiation to oligodendrocytes: insulin-like growth factor (*IGF*)^162^ and platelet derived growth factor (*PDGF*)^163,164^. While expression of *PDGF* genes spanned several cellular subclasses, expression of *IGF* was restricted to inhibitory interneurons and a small subset of microglia (**Fig. 8h**). *IGF1* expression decreases later in CPS in several inhibitory interneuron populations, suggesting that these inhibitory populations may be the main source of IGF1 and the driver of myelination changes (**Fig. 8i**). These data suggest compensation of oligodendrocyte loss by OPCs may be limited later in AD, in part a result of downregulation of necessary extracellular factors in inhibitory neurons; future functional studies are necessary to validate and understand these findings.

## Discussion

The aggregated knowledge about cellular organization of the human brain from the BRAIN Initiative Cell Census Network (BICCN)^24,30,31^, coupled with single cell and spatial transcriptomics methods now provide the means to create a comprehensive understanding of the cellular and molecular phenotypes and temporal progression of Alzheimer’s disease (AD). Here, we created an integrated atlas of AD in the middle temporal gyrus (MTG), selected both as a transition area in AD pathology^4,73^ and the region with the greatest aggregated knowledge in BICCN about cell type phenotypes^29,38,39^. The atlas illustrates the utility of the BICCN reference as a unifying framework that can be used to map cell types at high resolution, incorporate cell types and states not included in the reference, and replicate results. The core results presented here replicated across data modalities, cortical regions and datasets from independent studies, and have now been associated with fine-grain cell types and phases of AD progression. The results in turn demonstrate the value of this integration in defining a robust and specific series of cellular and molecular events that show what cells are affected, where they are (co)-localized, and when these events happen as disease pathology increases. All data presented here are publicly accessible through a suite of data resources available through SEA-AD.org, including viewers for donor metadata and image-based quantitative neuropathology, high resolution single nucleus transcriptome data viewing and mining (also at CZI’s <u>cellxgene</u>), genome browser (through <u>UCSC browser</u>^165^), data download (through <u>Sage</u> <u>Bionetworks</u>) and a novel tool (<u>MapMyCells</u>) for the community to map against the highly annotated SEA-AD cell classification.

The major driving concepts for the modeling of disease severity or progression are that neuropathology drives cellular and molecular changes, and that the local quantitative burden of neuropathology (QNP) can be used to model disease severity and progression. Aggregate scores like Braak^4,74^, Thal^3^, CERAD^166^ and ADNC^64^ measure distribution of pTau, aβ, and neuritic plaques but rely on binary present/absent scores that do not capture the level of pathology in any given brain region. Quantitative analyses of pTau and aβ demonstrated this very clearly, with enormous variation across donors within the same Braak stage or Thal phase. The AD research field has recognized the need for additional biomarkers of disease progression^66^. Quantitative neuropathology can capture the aggregate local burden of pathology, including identification of various cell types including neurons and disease-associated glial types, and other pathological proteins (pTDP-43, alpha-synuclein). Pseudotrajectory analysis using this information produced an ordering of donors that is strongly correlated with traditional brain-wide staging metrics as well as astrogliosis (the appearance of reactive astrocytes and increase glial number), and, surprisingly, decreasing NeuN labeling. pTDP-43 and alpha-synuclein did not show relationships to the pseudotrajectory, indicating that despite including AD/ADRD donors that the pseudotrajectory was largely modeling Alzheimer’s disease phenotypes. This approach appears to have been justified, in that effect sizes for cell proportion changes were substantially larger with disease progression modeled on QNP compared to more qualitative brain-wide staging.

The core results indicate two major epochs in AD progression (**Fig. 9**), including an early phase with slowly increasing neuropathology and a late phase with exponentially increasing neuropathology, culminating in the terminal state observed for the severely affected donors. In the early epoch, donors have sparse Aβ plaques (albeit increasing in size) and pTau-positive tangle bearing neurons, accompanied by early increases in inflammatory or reactive microglial^97^ and astrocytic states^10,19^ and associated gene expression changes in relevant inflammatory^113^ and plaque induced genes (**Fig.9b**). This epoch also features losses of oligodendrocytes and a dramatic increase in OPC differentiation and re-myelination factors that may represent a compensatory response similar to that seen in models of oligodendrocyte loss^167–170^. Neuronal cells exhibited loss of particular Sst interneuron types that down-regulate kinases and E3 ubiquitin ligases, but not the electron transport chain and ribosomal pathways (which were down-regulated in other neuronal populations) (**Fig.9b**). While these vulnerable Sst types were molecularly similar, they were highly morphologically diverse and included double bouquet cells that are seen in primates but not mice^171^. These vulnerable Sst supertypes localized to superficial cortical layers, whereas deeper layer Sst supertypes were not affected **(Fig. 9a)**, and exhibited distinctive electrophysiological properties, such as higher Sag, compared to unaffected supertypes. Importantly, these neurons are lost well before the exponential phase of accumulation of plaques and tangles. They are also not the neurons that bear the greatest burden of intracellular neurofibrillary tangles in temporal cortex (the L2/3 IT excitatory types^79^). Therefore, they are likely be vulnerable to lower levels of pathology, and may represent the initial trigger for circuit dysfunction in AD. Several prior reports have implicated Sst neurons in AD pathology as well^93,95^, but not at the same level of molecular, morphological, and electrophysiological detail. Our main results are consistent and replicated after integrating our data set across two cortical areas.

**Figure 9:**
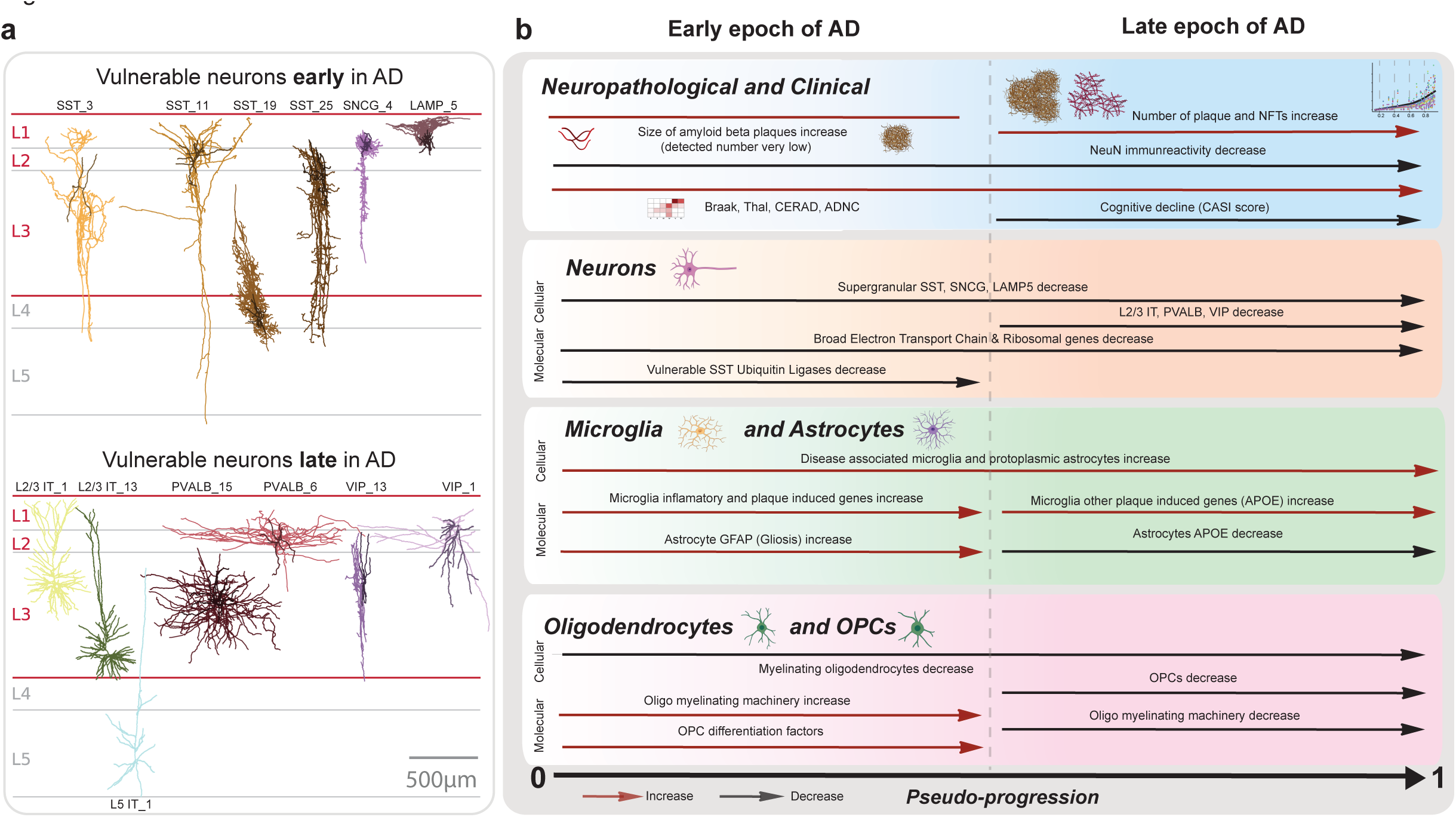
MTG cells impacted by AD, predominantly localizing to superficial layers, can be organized in two epochs: an early and a late phase. A) Diagram illustrating cortical columns with actual neuronal reconstruction from vulnerable populations (from non-AD donors) organized by the early (Top) and late (Bottom) disease epochs. During the early epoch, superficial Sst, Sncg and Lamp5 interneurons were lost. In the late epoch, most lost neurons localized superficially, (L2/3 IT, Pvalb and Vip), with the addition of deep cortical and striatum projecting L5 IT neurons. B) (first box) The dynamic changes associated with AD progression can be organized into an early and late epoch. In the early epoch, the first neuropathological event occurring is an increase in the size of sparse Aβ plaques, subsequently followed by an exponential aggregation of both pTau and plaque burden. A decrease in NeuN positive cells occurs throughout. (second box) Supragranular interneurons (Sst, Sncg, Lamp5) are lost early on. During this period, gene encoding the electron transport chain complex and ribosomal proteins are down-regulated broadly across neurons, except in the vulnerable Sst interneurons. In the latter cells, there is a strong downregulation of ubiquitin ligases and kinases. Later on, not only inhibitory cells (Pvalb and Vip) are lost but also long-range projecting pyramidal neurons (L2/3 IT and L5 IT). (third and fourth boxes) Nonneuronal cells accompany these changes with the early emergence of disease-associated microglia (DAM) and an increase protoplasmic astrocyte, while myelinating oligodendrocytes decrease their abundance. Concurrently, DAM upregulate inflammatory and plaque induced genes, while OPCs appear to attempt to compensate for oligodendrocyte loss by upregulating their OPC differentiating genes. Later, OPC cells are impacted and lost while myelination genes in oligodendrocytes are down-regulated.

What might be the consequences of an early loss of Sst neurons? Loss of inhibitory neurons would naturally be expected to disrupt excitatory/inhibitory balance, and impaired inhibition may therefore increase AD patient’s susceptibility to epilepsy, a clinical symptom found in more than 10% of patients^172^. This is supported by previous observations highlighting an anti-epileptic role of Sst+ interneurons^173^, and our observation that susceptible inhibitory interneurons express a high level of the HCN1 channel, dysfunction of which has been linked to several epileptogenesis pathways and the generation of hyperexcitability^174^. From a circuit perspective, Sst interneurons are uniquely positioned to exert control over both excitation and inhibition in the cortex, as they target excitatory and all other subclasses of inhibitory cortical neurons, but not themselves^175,176^. They also participate in a powerful disinhibitory loop via reciprocal connections with the VIP subclass^177^. Furthermore, they are known to mediate effects of arousal in cortical circuits^177,178^ under the effects of acetylcholine, which is also highly disrupted early in Alzheimer’s disease through the loss of cholinergic neurons in the basal forebrain^179^. SST neurons mediate context integration^180^ and exhibit responses to large-scale, high-intensity stimuli^181^, as well as contributing to cortical gamma-band oscillations^182^, which are important for inter-areal communication^183^. Indeed, functional properties of Sst neurons are consistent with a role in gating feedback between cortical areas^184,185^. Thus, reduction in numbers of Sst neurons is likely to have wide-ranging consequences beyond reduced network stability, affecting cognitive processes that rely on proper interactions in distributed brain areas, such as hierarchical integration of information, attention, or processing of novelty, all of which may result in deficits of learning and memory. Finally, it is possible that Sst neuron loss could also disrupt trophic support of connected neurons^186^, ultimately leading to the loss of long-range corticocortical connectivity that would be expected to affect cognitive function.

In the later epoch there is an exponential rise in Aβ and pTau pathology, continued increases in inflammatory microglia and astrocyte states, and a decrease in expression of both the OPC differentiation program and oligodendrocyte expression of myelin associated proteins (previously characterized by qPCR^187^). There is also broader loss of excitatory (L2/3 IT) and inhibitory (Pvalb and Vip neurons (**Fig.9b**). Vulnerable neuron types are specific too, including a subset of the supertypes within each broader subclass and are largely localized to the upper layers of the cortex (Fig.9a). For example, there was a selective loss of excitatory neurons in supragranular layers (L2/3 IT), as described previously based on cell counting of non-phosphorylated heavy chain neurofilament protein (labeling with SMI-32 antibody) positive neurons^94^, and more recent single cell analyses^188^. SMI-32 predominantly labels long-range ipsilateral-projecting corticocortical neurons in monkey^189^, and selectively labels human L2/3 IT types in layer 3, including the largest neurons that do not appear to have mouse homologues^29^.

Putting these two epochs together, the overall progression suggests a sequence of events in which early microglial activation at low levels of pathology triggers reactive astrocytes and potentially oligodendrocyte loss^190^. Further, the early loss of Sst neurons in upper cortical layers could lead to excitatory/inhibitory circuit imbalance (**Fig. 9b**) that could in turn lead to loss of other co-localized (and thus likely connected) excitatory and inhibitory neurons, including long range corticocortical (L2/3 IT) neurons that contribute to cognitive decline. Donors with the steepest memory cognitive decline late in life showed particularly broad cellular dysfunction, suggesting that this was not due to poor quality samples but rather a biological outcome of AD pathology and subsequent cognitive decline. These severely affected donors had lower transcription and reduced chromatin accessibility that may correspond to senescent states^191^, or global epigenome dysregulation indicative of cell identity loss^143^.

The results presented here in MTG demonstrate that systematic application of single cell genomic and spatial technologies coupled with quantitative neuropathology can effectively model disease progression across the spectrum of AD severity. Importantly, the BICCN reference now allows integration and direct comparison across many studies to use common annotation of the same cell types and states, and to cross-validate results to demonstrate their robustness and consistency. The remarkably similar cell vulnerabilities in MTG and DLPFC (and across studies) indicates there may be a common pattern of cellular and circuit dysfunction in response to neuropathology progression across the brain as well. The annotation of the BICCN reference now allows interpretation of AD cellular phenotypes at the finest level of circuit neuroscience, and to consider AD as a circuit disorder that ultimately affects cognitive function. These molecular phenotypes may help develop new biomarkers, while vulnerable cell populations may present new targets for therapeutic intervention using tools that can now be reliably developed for genetic targeting of specific cell populations^192,193^. This strategy can now be extended to understand disease progression across more diverse cohorts and across brain regions within individuals, to identify commonalities across brain regions and the earliest events in AD pathology when therapeutic interventions may be most effective.

## Acknowledgements

The Seattle Alzheimer’s Disease Brain Cell Atlas (SEA-AD) consortium is supported by the National Institutes on Aging (NIA) grant U19AG060909. Study data were generated from postmortem brain tissue donated to the University of Washington BioRepository and Integrated Neuropathology (BRaIN) laboratory and Precision Neuropathology Core, which is supported by the UW Alzheimer’s Disease Research Center (NIA P30AG066509, previously P50AG005136), the Adult Changes in Thought (ACT) Study (NIA U19AG066567). ACT Data collection for this work was additionally supported, in part, by prior funding from NIA (U01AG006781), and the Nancy and Buster Alvord Endowment (to C.D.K.). We thank the participants of the ADRC and the ACT study for the data they have provided and the many ADRC and ACT investigators and staff who steward that data. You can learn more about UW ADRC at https://depts.washington.edu/mbwc/adrc and ACT at https://actagingstudy.org/. We thank AMP-AD, Sage Bionetworks and the Open Data Registry on AWS for hosting various datasets from this study. We thank the Rush Alzheimer’s Disease Center (RADC) for sharing donor metadata from the ROS/MAP studies.

The results published here are in part based on data obtained from the AD Knowledge Portal (https://adknowledgeportal.org). This includes data from:

Mathys et. al., 2019: Samples for this study were provided by the Rush Alzheimer’s Disease Center, Rush University Medical Center, Chicago. Data collection was supported through funding by NIA grants P30AG10161, R01AG15819, R01AG17917, R01AG30146, R01AG36836, U01AG32984, U01AG46152, the Illinois Department of Public Health, and the Translational Genomics Research Institute. Zhou et. al., 2020 Cain et. al., 2022, Olah et.al., 2020, Green et. al., 2023: Study data were provided by the Rush Alzheimer’s Disease Center, Rush University Medical Center, Chicago. Data collection was supported through funding by NIA grants P30AG10161 (ROS), R01AG15819 (ROSMAP; genomics and RNAseq), R01AG17917 (MAP), R01AG30146, R01AG36042 (5hC methylation, ATACseq), RC2AG036547 (H3K9Ac), R01AG36836 (RNAseq), R01AG48015 (monocyte RNAseq) RF1AG57473 (single nucleus RNAseq), U01AG32984 (genomic and whole exome sequencing), U01AG46152 (ROSMAP AMP-AD, targeted proteomics), U01AG46161(TMT proteomics), U01AG61356 (whole genome sequencing, targeted proteomics, ROSMAP AMP-AD), the Illinois Department of Public Health (ROSMAP), and the Translational Genomics Research Institute (genomic). Additional phenotypic data can be requested at www.radc.rush.edu. snRNAseq Data: Study data were generated from postmortem brain tissue provided by the Religious Orders Study and Rush Memory and Aging Project (ROSMAP) cohort at Rush Alzheimer’s Disease Center, Rush University Medical Center, Chicago. This work was funded by NIH grants U01AG061356 (De Jager/Bennett), RF1AG057473 (De Jager/Bennett), and U01AG046152 (De Jager/Bennett) as part of the AMP-AD consortium, as well as NIH grants R01AG066831 (Menon) and U01AG072572 (De Jager/St George-Hyslop).

Mathys et.al., 2023: Study data were generated from postmortem brain tissue provided by the Religious Orders Study and Rush Memory and Aging Project (ROSMAP) cohort at Rush Alzheimer’s Disease Center, Rush University Medical Center, Chicago. This work was supported in part by the Cure Alzheimer’s Fund, NIH grants AG058002, AG062377, NS110453, NS115064, AG062335, AG074003, NS127187, MH119509, HG008155 (M.K.), RF1AG062377, RF1 AG054321, RO1 AG054012 (L.-H.T.) and the NIH training grant GM087237 (to C.A.B.). ROSMAP is supported by P30AG10161, P30AG72975, R01AG15819, R01AG17917. U01AG46152, U01AG61356.

## Author contributions

Study design: B.P.L., C.D.K., E.B.L., E.S.L., J.A.M., J.L.C., K.A.S., K.J.T., M.H., M.I.G., M.W., P.K.C., R.D.H., S.J., T.J.G. Data generation: A.G., A.K., A.L.N., A.M.W., A.R., A.T., B.E.K., B.P.L., B.R.L., C.A.P., C.L.M.D., C.R., C.S.L., D.H., D.M., E.B.L., E.J.M., E.M., J.A., J.C., J.G., J.G., J.G., J.L.C., J.M., J.T.M., K.A.S., K.B., L.M.K., M.C., M.D., M.H., M.K., M.K., M.L., M.M., M.P., M.T., N.D., N.G., N.J.V.C., N.M., N.P., N.P., N.V.S., P.A.O., R.C., R.D.H., R.F., R.M., S.A.S., S.D., S.J., S.K., S.T.B., T.C., T.E., T.P., W.H., Z.M. Data analysis: A.A., B.E.K., B.L., B.R.L., C.S.L., E.G., G.A.S., G.M., J.A., J.A.M., J.C., J.L.C., J.T., K.A.S., K.J.T., L.N., M.I.G., M.K., O.K., R.C., R.D., S.A.S., S.M., S.W., T.J., V.M.R., X.Z., Y.D. Data interpretation: A.A., B.E.K., B.L., B.P.L., B.R.L., C.D.K., C.L.M.D., D.H., E.B.L., E.S.L., J.L.C., J.T., J.W., K.A.S., K.J.T., L.N., M.I.G., O.K., P.K.C., R.D., R.D.H., S.M., S.,W., V.M.R., X.Z., Y.D. Project management: A.L.N., A.M.S., C.M.P., E.J.M., E.S.K., E.S.L., J.K.N., J.W., K.A.S., K.P.M., L.E., L.E.F., M.S., N.G., S.M.S., T.T. Creating web product: A.J.C., B.S., C.T., E.S.K., E.S.L., J.A.M., J.C., K.J.T., L.N., M.H., M.S., M.W., N.J., R.G., S.M., T.D., T.M., T.T. Writing manuscript: B.E.K., B.L., C.D.K., E.S.K., E.S.L., J.A.M., J.L.C., K.J.T., M.H., M.I.G., O.K., P.K.C., R.D.H., S.M., V.M.R.

## Extended Data Figure Legends

**Extended Data Figure 1:**
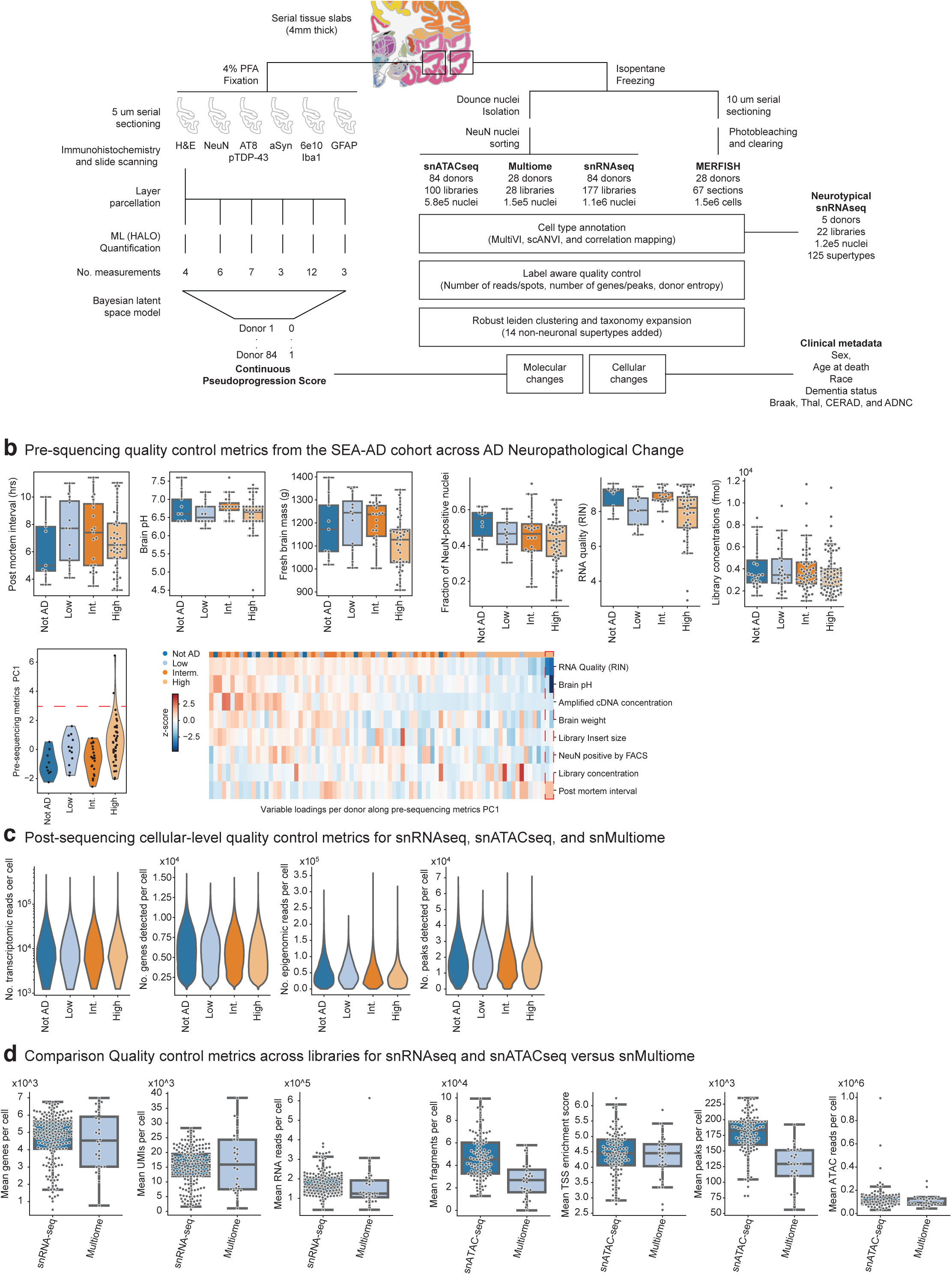
SEA-AD Brain Cell Atlas study design. A) Schematic detailing experimental design for applying quantitative neuropathology, single nucleus RNA sequencing (snRNAseq), single nucleus ATAC sequencing (snATAC-seq), single nucleus Multiome (Multiome), and multiplexed error robust fluorescence in situ hybridization (MERFISH) to middle temporal gyrus (MTG) of SEA-AD donors as well as the analysis plan for construction of a pseudo-progression score from quantitative neuropathology, integration across -omics data modalities, common cell type mapping to the BRAIN initiative reference, and use of demographic and clinical metadata to identify cellular and molecular changes in AD. B) Top, boxplots showing pre-sequencing quality control metrics for donor tissue (e.g. PMI, RIN, brain pH and mass) and single nucleus preparations (e.g. fraction of NeuN positive nuclei and library concentration) organized by AD Neuropathological Change (ADNC). Bottom, A donor by metric matrix was constructed for the values indicated, using a simple average for variables that had multiple values per donor (e.g. multiple sequencing library concentrations). Principle component analysis (PCA) was then run on the matrix. Bottom and left, Violin plot showing the eigenvalues for each donor along the first principle component organized by ADNC. Bottom and right, heatmap showing z-scores of the pre-sequencing quality control metrics (rows) in each donor (columns). Donors and metrics are ordered based on the first principle component eigenvalues and eigenvectors. Red dashed box, two outlier values along first principle component for two donors that were driven by low RIN and brain pH. C) Violin plots showing cellular-level post-sequencing quality control metrics for single nucleus transcriptomics, chromatin accessibility and multiome data organized by ADNC. Significant p-values: NeuN Fraction Not AD versus High=0.05. D) Violin plots comparing library-level post-sequencing quality control metrics of snRNA-seq to snMultiome (left) and snATAC-seq to snMultiome (right).

**Extended Data Figure 2:**
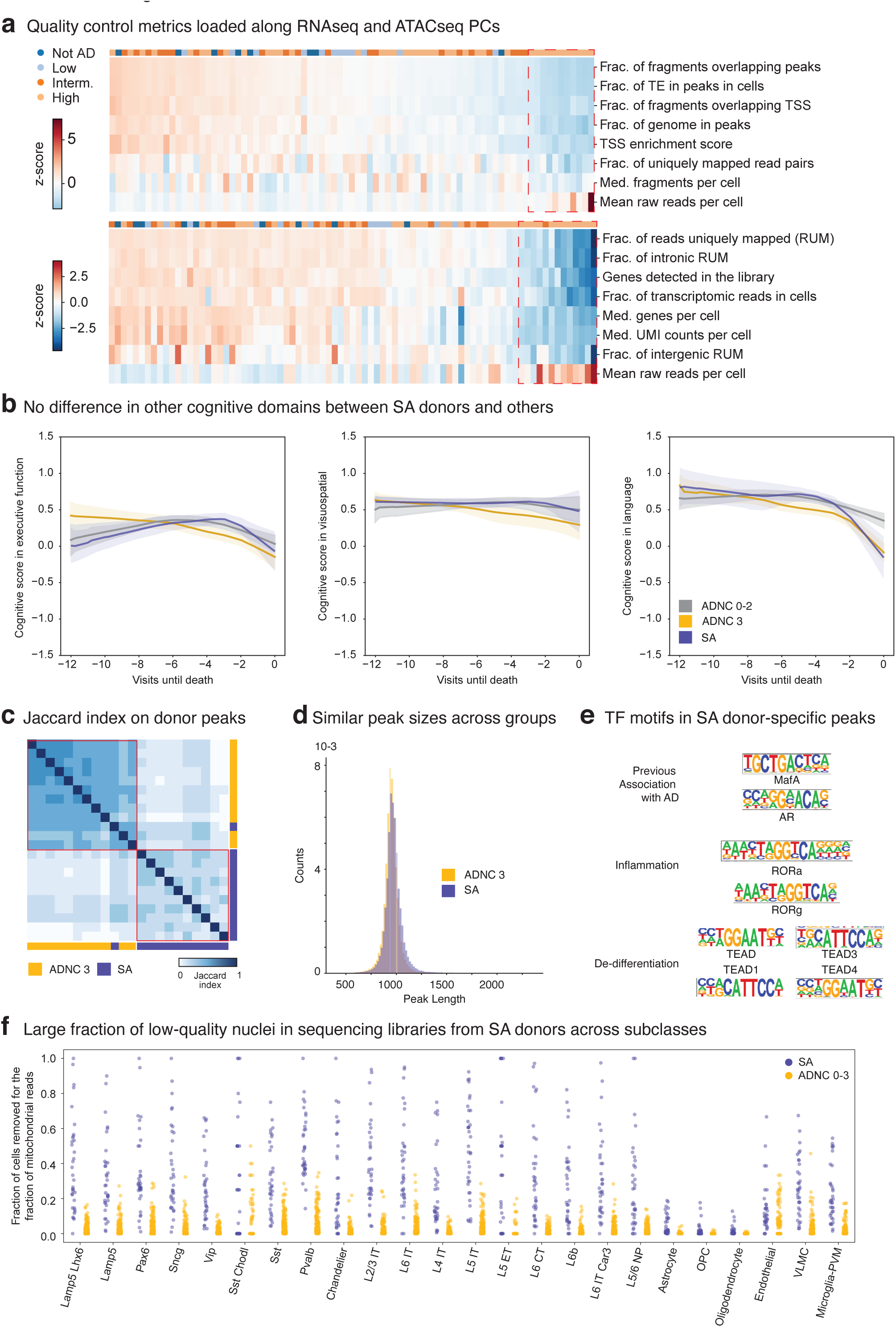
Altered multimodal metrics within severely affected donors. A) Donor by metric matrices were constructed for the library-level post-sequencing quality control values indicated, using a simple average when multiple libraries were sequenced per donor. Principle component analysis (PCA) was then run on each matrix. Heatmaps showing z-scores of snATAC-seq (top) and snRNA-seq (bottom) metrics (rows) in each donor (columns). Donors and metrics are ordered based on the first principle component eigenvalues and eigenvectors. Red dashed boxes, donors with outlier eigenvalues along each PC. B) LOESS regression on longitudinal cognitive scores in the executive, visuospatial, and language domain across ADNC 0-2 (Not AD to Intermediate) in grey, ADNC 3 donors that were not severely affected in gold, and ADNC 3 donors that were in purple. Uncertainty represents the standard error from 1000 LOESS fits with 80% of the data randomly selected in each iteration. Significant p-values for cognitive decline: SA donors versus ADNC 0-2=0.009, Other ADNC 3 versus ADNC 0-2=0.021. C) Heatmap showing the pairwise jaccard distances based on the peak universes from 11 randomly selected ADNC 3 donors (yellow) and all 11 severely affected donors (purple) hierarchically ordered. Red boxes, two clusters within the hierarchy that largely correspond to the separation between ADNC3 and SA donors. D) Histogram showing the distribution of peak lengths of accessible regions in ADNC 3 (yellow) and severely affected donors (purple). E) Transcription factors binding sites enriched in chromatin accessible regions uniquely found in severely affected donors organized by their gene ontology category. Transcription factors that bind to them are indicated. F) Stripplot showing the fraction of cells removed from each library for having too many mitochondrial reads during quality control organized by subclass and by severely affected donors (purple) and ADNC 0-3 donors (yellow).

**Extended Data Figure 3:**
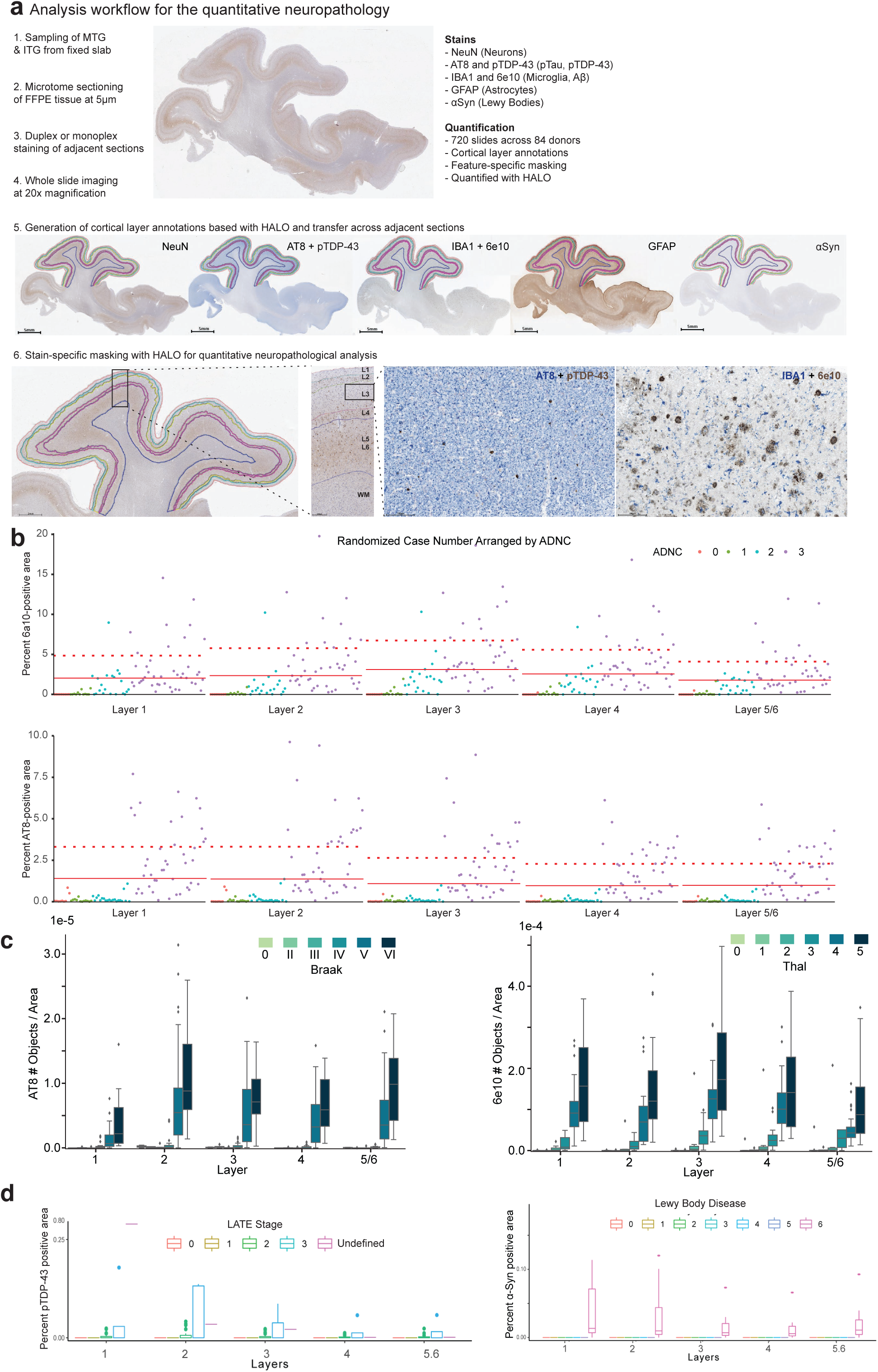
Human MTG neuropathological stains track brain-wide pathological states. A) Schematic depicting neuropathological data acquisition pipeline (ordered 1 to 6) B) Scatterplots showing the percent of 6e10 (Aβ)-positive (top) and percent of AT8 (pTau)-positive voxels (bottom) across donors, stratified by layer and color-coded/organized by ADNC. Within a particular ADNC group, donors are randomized. Red solid line, mean value across donors. Red dashed line, mean plus 1 standard deviation C) Boxplots showing the number of pTau-bearing cells per unit area organized by Braak stage (left) and number of Aβ plaques per unit area organized by Thal phase (right) across donors. Note, in later stages there is considerable variability in plaque and tangle number, underscoring limitations in classical staging. D) Boxplots showing the percent of pTDP-43-positive voxels (left) and percent of α-Syn-positive (α-Synuclein) voxels across donors organized by to LATE-NC stage (left) and Lewy Body Disease stage (right). Lewy Body Disease is coded numerically (0 = Not or Incompletely Assessed, 1=Not Identified, 2=Amygdala-predominant, 3=Brainstem-predominant, 4=Limbic (Transitional), 5=Olfactory bulb only, 6=Neocortical). Note, only donors in later stages have large accumulation of co-pathology.

**Extended Data Figure 4:**
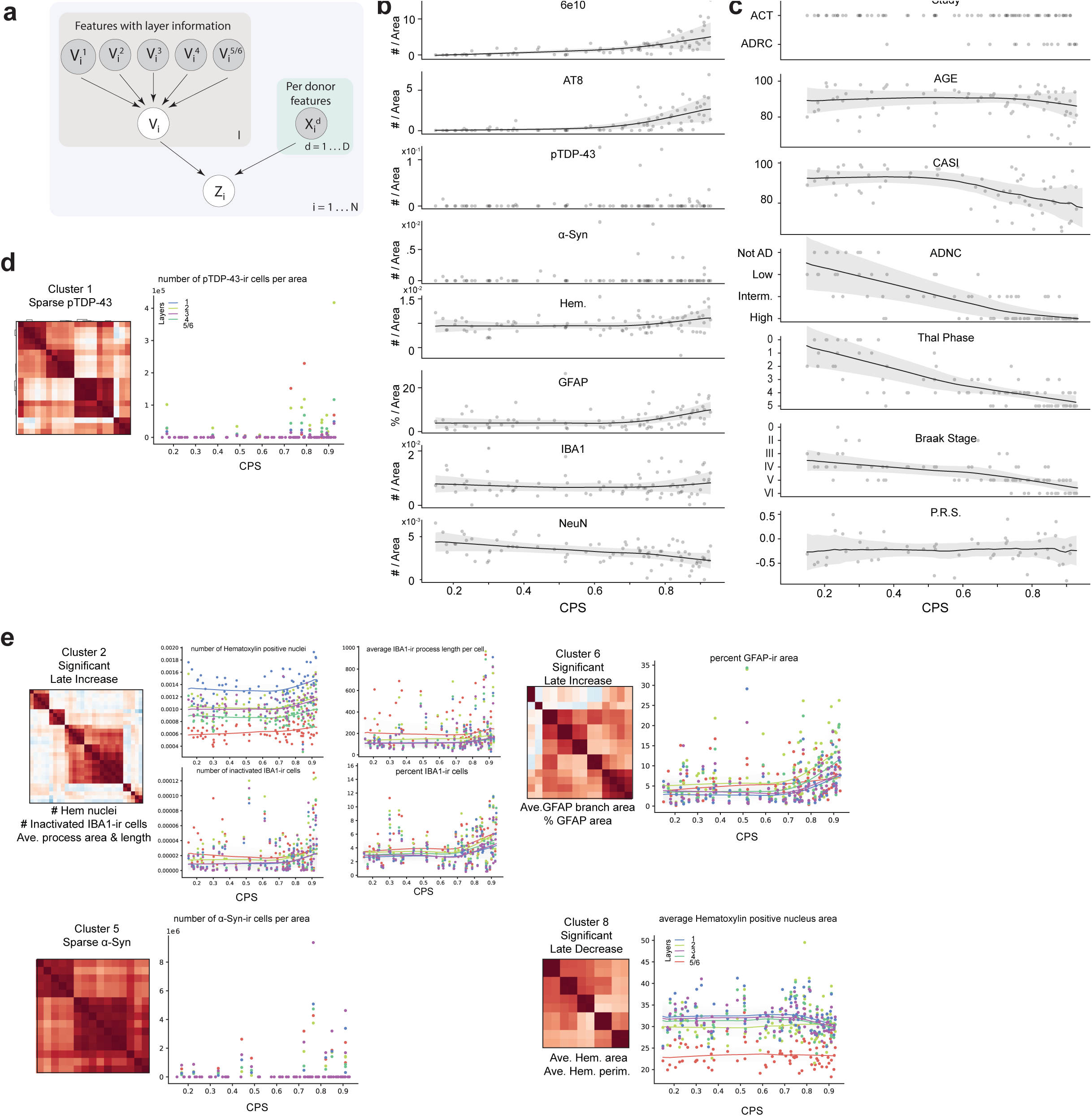
MTG pseudo-progression scores orders quantitative neuropathological variables following increasing disease severity. A) Graphical model used to infer the continuous pseudo-progression score (CPS). B, C) LOESS regression plots relating mean quantitative neuropathological (QNP) variables across layers (B) and demographic/clinical metadata (C) indicated to CPS. Dots represent individual donor values. Uncertainty in each line represents the standard error from 1000 LOESS fits with 80% of the data randomly selected in each iteration. Note, variables from (C) were not used to construct the model. CASI, Cognitive Abilities Screening Instrument; ADNC, AD Neuropathological Change; PRS, Polygenic Risk Score. D) Left, Subset of heatmap from Figure 2c showing co-correlation of QNP variables in cluster 1. Right, Scatterplot showing how the QNP variable number of pTDP-43 positive cells per unit area, which is within correlation cluster 1, relates to CPS. Dots represent values from each donor in the cortical layer indicated, lines are LOESS regressions for measurements across donors within each layer. E) Same plots as in (D) but for clusters 2, 6, 5, and 8.

**Extended Data Figure 5:**
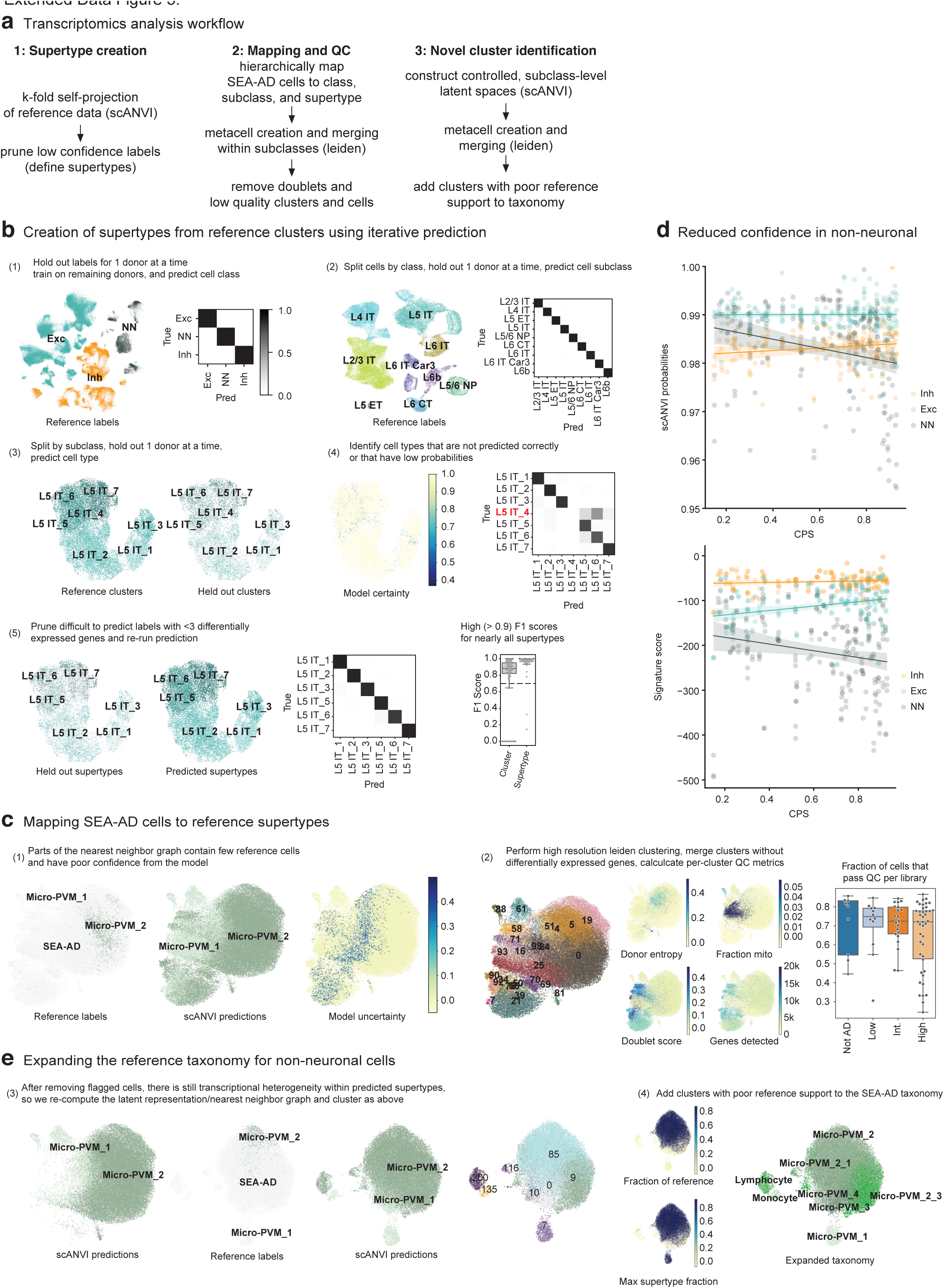
Pipeline for the creation of the SEA-AD MTG taxonomy. A) Schematic showing steps involved in supertype creation from snRNA-seq data in neurotypical reference donors, mapping and quality control on SEA-AD snRNA-seq and snMultiome data, and expansion of the BRAIN Initiative taxonomy to include novel types and states found in SEA-AD. B) Hierarchical procedure for the creation of robustly mappable cell types, termed “supertypes”. Labels from 1 of 5 reference donors was systematically held out and predicted using a deep generative model (DGM) trained on the remaining 4 donors. Steps 1 to 3 represent mapping cells to one of three classes, splitting each class and mapping to one of 24 subclasses, splitting each subclass and mapping to one of 151 clusters from the original BRAIN initiative taxonomy. In each step, we include scatterplots with UMAP coordinates for exemplar populations from references donors along with heatmaps showing confusion matrices. Note, near perfect mapping is achieved to the subclass level and clusters are occasionally missed a the cluster level. In step 4, we identified these hard-to-map clusters (example L5 IT_4 cluster shown in red) and removed labels for these cells in the training data. 26 of 151 clusters were pruned, mostly representing cell types that were intermediates of others. Finally, in step 5 we repeat mapping with the 125 highly mappable supertypes and show consistently high F1 scores across them (box and whisker plot). C) After hierarchically mapping SEA-AD nuclei to supertypes using the same approach as above, we filtered low quality nuclei within subclasses (The microglia subclass is shown as an example). Left, scatterplots showing the UMAP coordinates of all SEA-AD and reference nuclei within the microglia subclass. In the first plot, reference nuclei are labeled and colored and SEA-AD nuclei are in light grey. In the second and third plots, we show the supertype predictions for each nucleus from the DGM as well as the uncertainty in the prediction (darker nuclei are more uncertain). In the fourth plot we show robust, high resolution Leiden clusters and color them by their quality control metrics (i.e. donor entropy, mean fraction of mitochondrial reads, mean doublet score, and mean number of genes detected). Clusters were flagged and removed based on these metrics. The far right box and whisker lot shows the fraction of nuclei removed per library organized by AD Neuropathological Change (ADNC). D) Scatterplots showing scANVI probabilities (top) and supertype signature scores (bottom) organized by cell classes. Lines represent linear regressions. Note, decreasing probabilities and signature scores for non-neuronal supertypes, but not others. E) After removing low quality nuclei new latent representations were learned with DGMs, which were then underwent robust Leiden clustering. Clusters with low fractions of nuclei from neurotypical reference donors (<10%) were added to the taxonomy. The first scatterplot shows the UMAP coordinates of nuclei that passed quality control filtering in (C) from the same latent representation. The next three scatterplots show the UMAP coordinates of the same cells based on the new latent representation colored by reference versus SEA-AD nuclei (light grey), scANVI predictions, and robust Leiden clusters. The next two scatterplots show the fraction of all reference nuclei per cluster (top) and the max fraction of any supertype per cluster (bottom). The final scatterplot is colored by the final SEA-AD taxonomy with the new clusters that had poor reference support added.

**Extended Data Figure 6:**
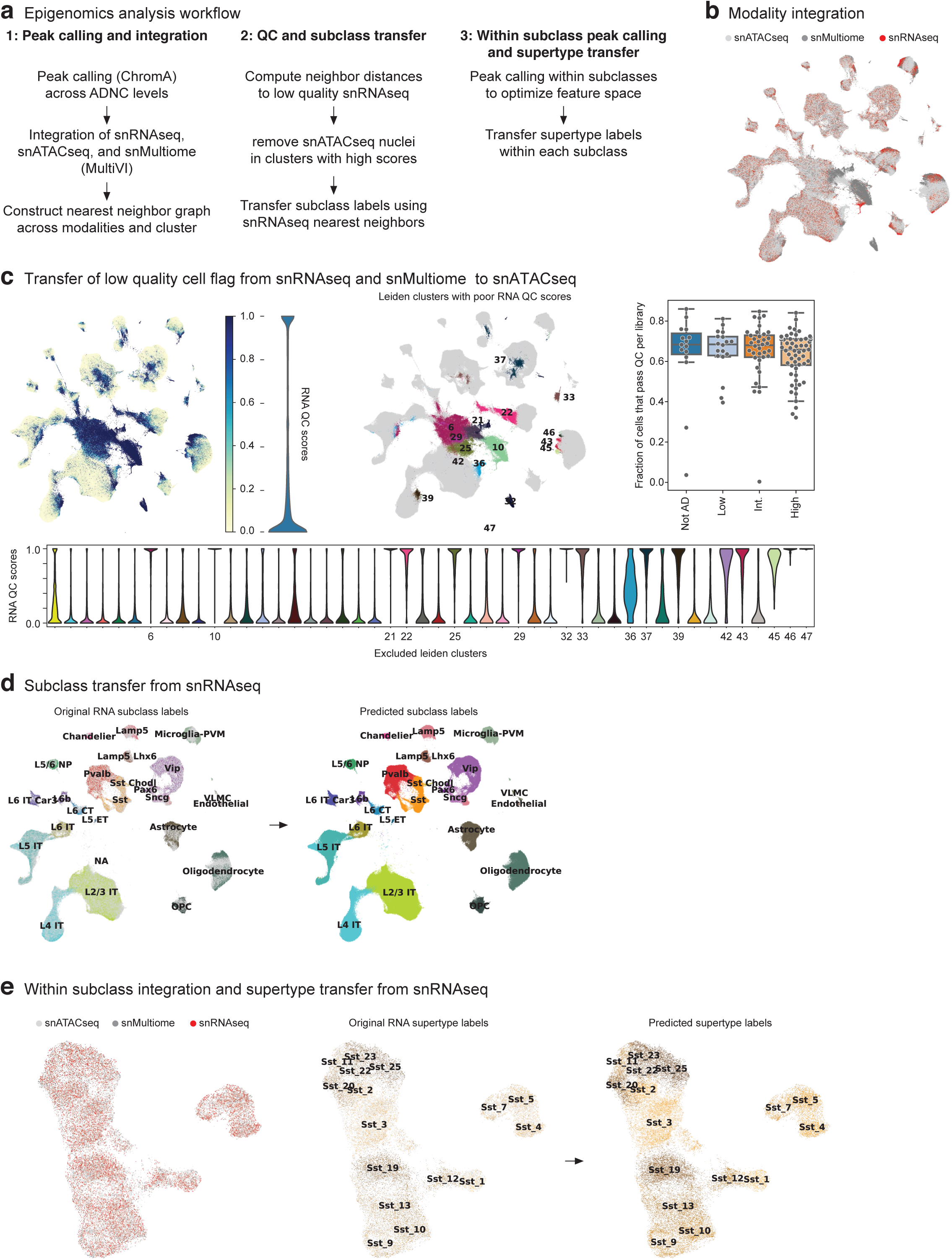
Pipeline for the annotation of chromatin accessibility data sets. A) Schematic showing steps involved in processing the SEA-AD snATAC-seq data, which include global peak calling and modality integration, quality control filtering and subclass mapping, and within subclass peak calling and supertype mapping. B) Scatterplot showing the UMAP coordinates of all nuclei profiled in the middle temporal gyrus (MTG) color coded by indicated data modalities. C) Top and left, Same scatterplot as in (B) but color coded by low quality cell score (left) and (right) by Leiden clusters with mean low quality cell scores greater than 0.5. Violin plot to the right of the first plot shows the binary distribution of the low quality cell scores (RNA QC score). Bottom, violin plots showing the distribution of the low quality cell score per Leiden cluster, with the number of those that were flagged indicated. Top and right, box and whisker plot showing the fraction of cells in each snATAC-seq library that were filtered during quality control. D) Scatterplots showing the UMAP coordinates from (B) of only the high quality nuclei colored by neurotypical reference subclasses versus SEA-AD in light grey (left) and by predicted subclass (right). E) Scatterplots showing UMAP coordinates of nuclei from 1 example subclass (Sst) based on integrated space constructed with subclass-specific peaks. Plots are color coded by modality (left), by reference supertypes versus SEA-AD in light grey (middle) and by predicted supertype (right).

**Extended Data Figure 7:**
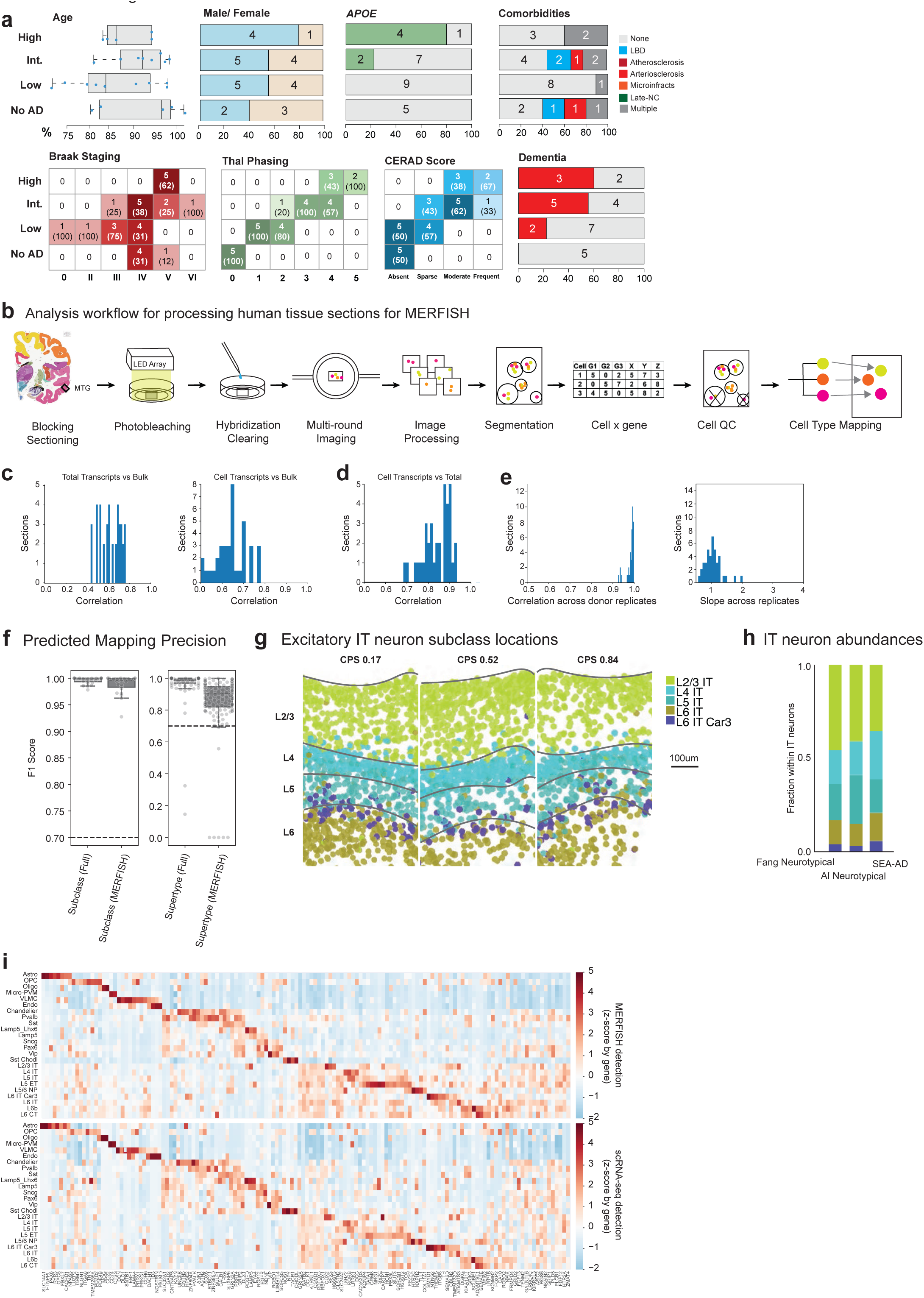
Pipeline for the acquisition of high quality spatial transcriptomic data in the human MTG. A) Top, SEA-AD MERFISH cohort demographics stratified by AD neuropathological change (ADNC) score. Age at death is represented by box-and-whisker plots. The median is indicated by the solid line within the box and donor values are represented as points. The fraction of donors that have a biological sex of male or female, that have an APOE4 allele, and that have a co-morbidity are shown as bar charts per ADNC level, with the number of donors in each group indicated. Bottom, SEA-AD MERFISH cohort composition stratified by ADNC on the left versus Braak stage (measuring distribution of neurofibrillary tangles across the brain), Thal Phase (measuring distribution of amyloid beta plaques across the brain), and CERAD score (measuring the distribution of neuritic plaques across the brain) as heatmaps. The number of donors in each box are indicated with the fraction in Braak, Thal, and CERAD stages in parentheses. Darker colors represent a higher fraction of donors. The fraction of donors across ADNC that have dementia (red) or not (grey) are shown as bar plots. Numbers indicate the number of donors in each group. B) Schematic showing the Spatial Transcriptomics workflow. Tissue blocks for profiling are cut from a frozen donor slab and cryosectioned. Sections are photobleached to reduce autofluorescence, hybridized with the probe panel, digested to clear light scattering elements, then processed through multi-round imaging on the MERSCOPE. Stitched and processed images are segmented to provide cell boundaries and a cell x gene table with cell locations is created. Cells are mapped to their transcriptomic type and assigned to their location within the tissue to create cell maps. C) Histograms showing the correlation between total slide transcripts (left) or transcripts within cells (right) and bulk RNAseq across sections. D) Histogram showing the correlation between total slide transcripts and transcripts in cells. E) Left, histogram showing the correlation in total slide transcripts across sections from the same donor. Right, Histogram showing the slope from a linear regression comparing total slide transcripts across sections from the same donor. F) Box and whisker plot showing F1 scores for subclasses (left) and supertypes (right) from the procedure where 1 donor was systematically held out at a time in neurotypical reference snRNA-seq data where the model could use all genes (Full) or only the 140 genes in the MERFISH panel (MERFISH). Note, when the mapping is limited to the 140 genes in the MERFISH panel, F1 scores decreased in both taxonomic levels but were still above 0.7 for the vast majority of supertypes. G) Scatterplots showing the positions of excitatory intratelencephalic (IT) neurons as dots from example sections from donors with an early (0.17), middle (0.52) and late (0.84) continuous pseudo-progression score (CPS) color coded by their subclass. The spatial positions of these cells is consistent with their mapped identity across CPS. H) Barplot showing the relative abundance of excitatory IT neurons across data collection efforts in neurotypical specimens from previous studies compared to SEA-AD data. I) Heatmaps showing the average gene expression levels of genes included in the 140 gene MERFISH panel at the subclass level in snRNA-seq (top) and MERFISH (bottom) data from the middle temporal gyrus (MTG). Note, highly similar expression patterns across modalities.

**Extended Data Figure 8:**
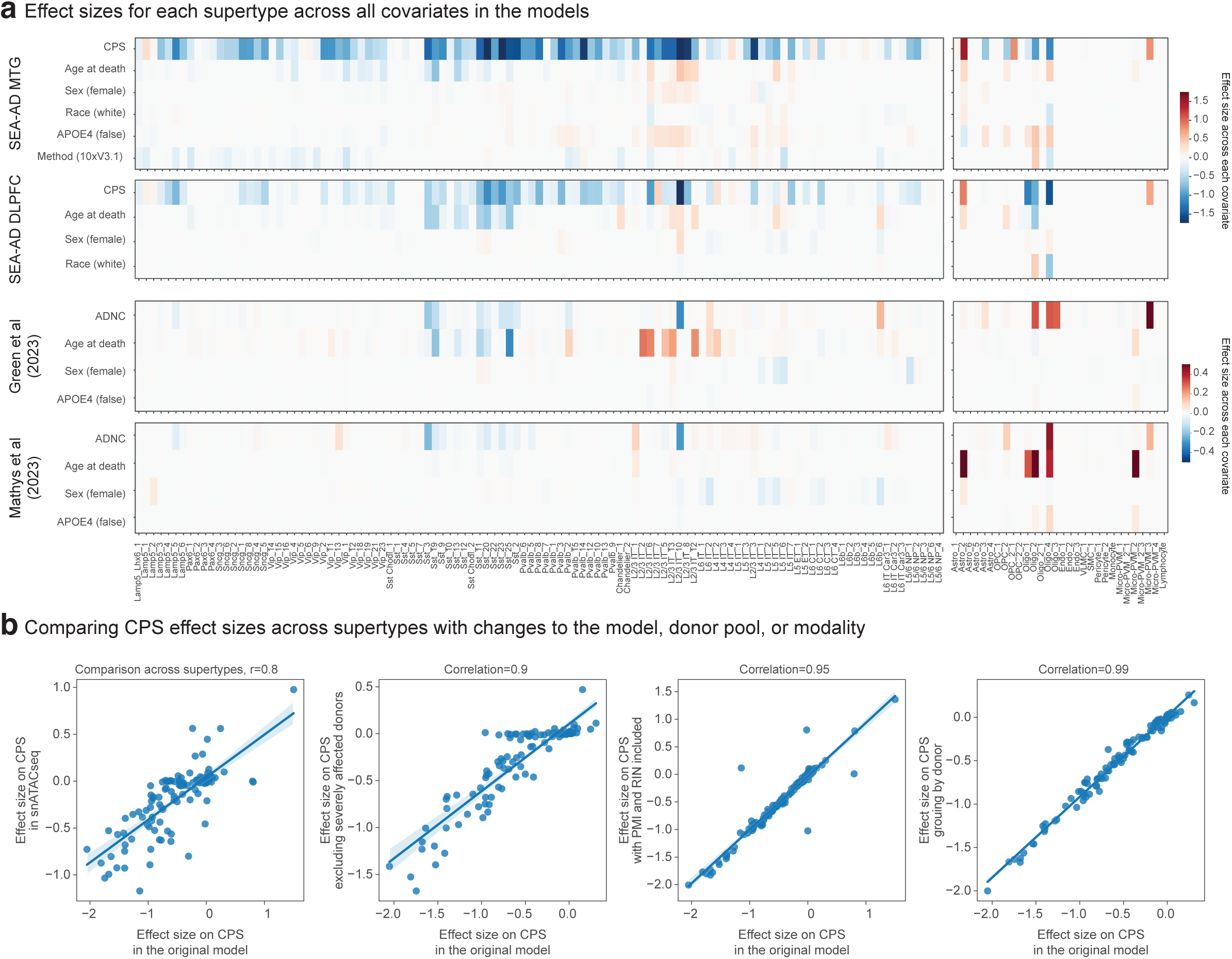
scCODA model covariates. A) Heatmaps showing the effect sizes of relative abundance changes along each covariate from neuronal (left) and non-neuronal (right) scCODA models in the SEA-AD snRNA-seq and snMultiome MTG dataset (first), the SEA-AD snRNA-seq DLPFC dataset (second), Green et al_2023 snRNA-seq dataset (third), and Mathys et al_2023 snRNA-seq dataset (fourth). B) Scatterplots relating the effect sizes of each supertype along CPS from the scCODA model on the SEA-AD snRNA-seq and snMultiome MTG dataset to a similar model run on the SEA-AD snATAC-seq MTG dataset (first), to a model run without the severely affected donors (second), to a model that included post mortem interval (PMI) and RIN as covariates (third), and to a model that grouped data by donor instead of by library (fourth). Linear regressions and their R values are shown. Note strong agreement across all tests.

**Extended Data Figure 9:**
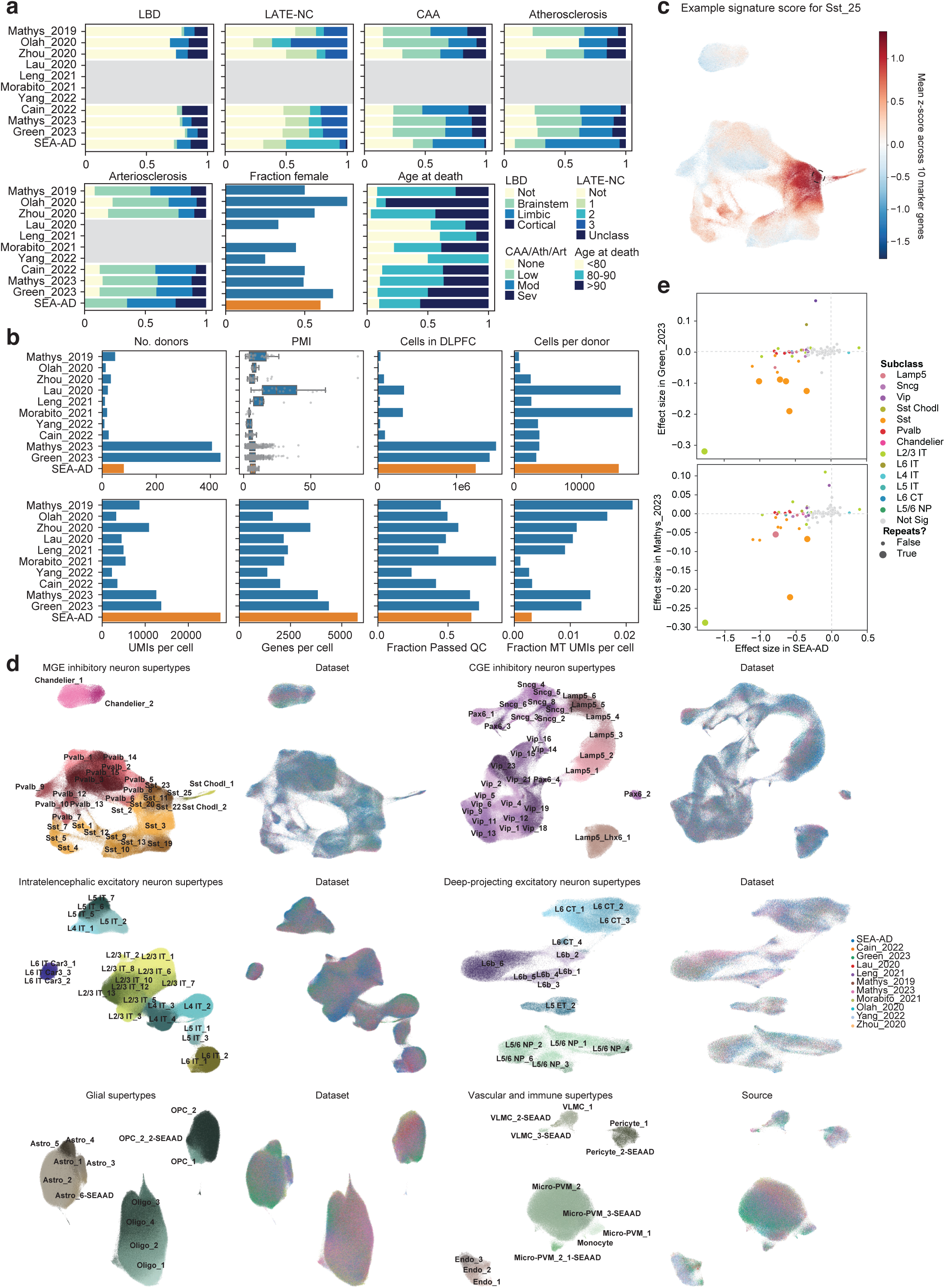
Integration of publicly available snRNA-seq datasets. A) Barplots showing the fraction of donors in each of the publicly available snRNA-seq datasets that we harmonized metadata for and integrated classified in co-pathology neuropathological stages (LBD, Lewy Body Disease; LATE-NC, limbic-predominant age-related TDP-43 encephalopathy neuropathologic changes; CAA, Cerebral amyloid angiopathy; Ath, Atherosclerosis; Art, Arteriosclerosis), that were female, or were in defined age groups. Grey boxes, metadata that was unavailable. B) Box and whisker or barplots showing quality control metrics across each of the publicly available datasets. Metrics for the SEA-AD DLPFC snRNA-seq dataset are shown at bottom in orange for comparison. C) Scatterplot showing UMAP coordinates for MGE-derived inhibitory interneuron supertypes across all publicly available and the SEA-AD DLPFC dataset. Nuclei or cells are colored based on the signature score for Sst_25, which are indicated with the black dashed circle. D) Scatterplots showing UMAP coordinates of all supertypes within their cellular neighborhoods (i.e. MGE-derived inhibitory neurons, CGE-derived inhibitory neurons, Intratelencephalic excitatory neurons, Deep-projecting excitatory neurons, glial cells, and vascular and immune cells. In each neighborhood on left are nuclei and cells colored by supertype and on right cells are colored by dataset. E) Scatterplots relating the effect size for the change in relative abundance across supertypes in the SEA-AD DLPFC dataset to those observed in the Green_2023 (top) and Mathys_2023 (bottom) datasets. Each point is a supertype colored by their subclass and supertypes that are significant in both datasets have bigger circles. Dashed grey lines are at 0. Note, several Sst, 1 L2/3 IT and 1 Lamp5 supertypes that have significant negative effect sizes in both datasets.

**Extended Data Figure 10:**
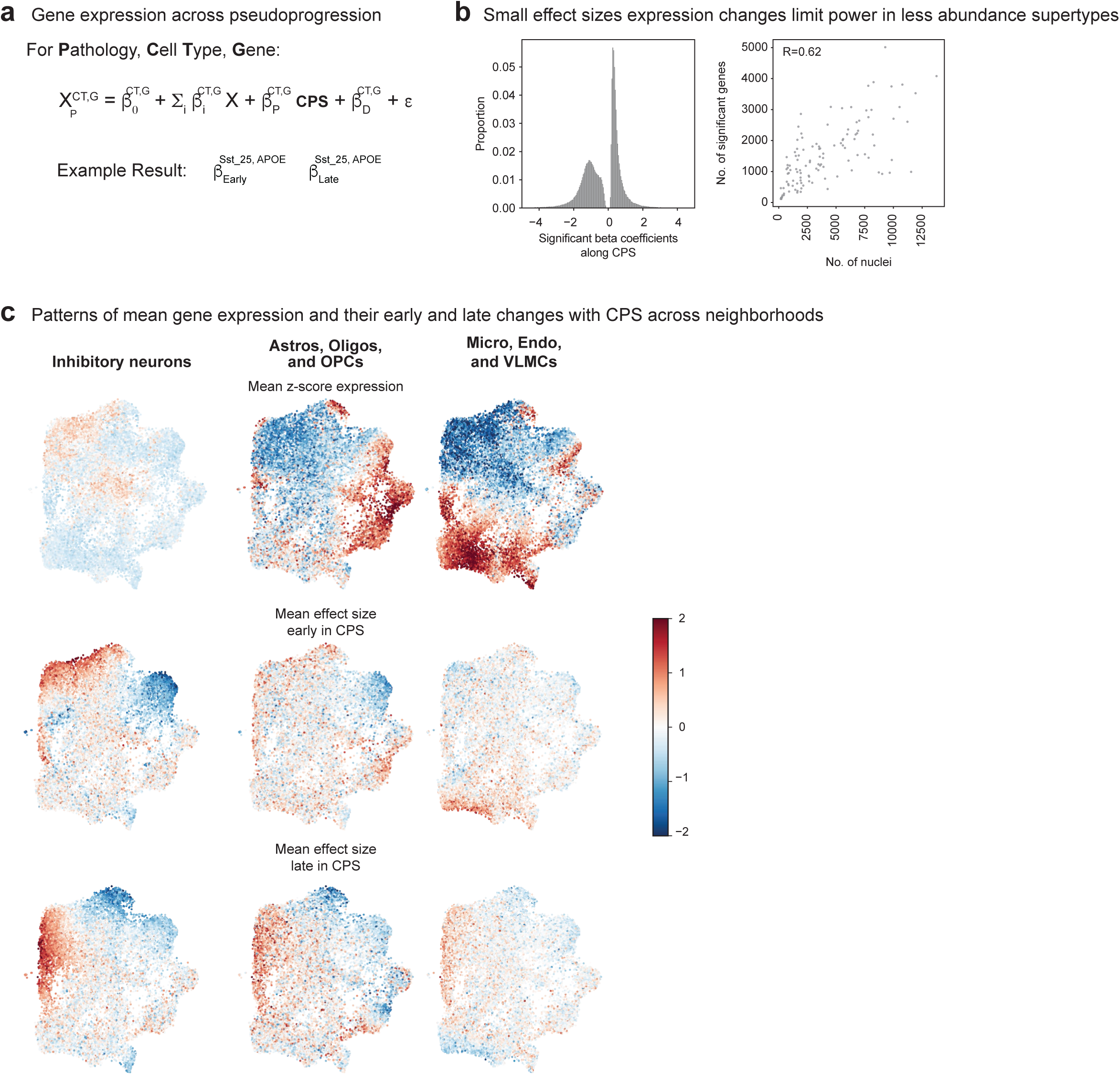
Construction of the “gene-dynamic space”. A) Schematic for identifying differentially expressed genes in each disease epoch along CPS using a generalized linear mixed model. B) Left, histogram showing the effect sizes across all supertypes of significantly changed along CPS. Note, many significant changes had relatively small effect sizes. Right, Scatterplot showing a weak (but present, R=0.62) correlation between the number of nuclei and number of genes called as significantly changed along CPS. This suggests SEA-AD may lack the statistical power necessary to identify all of the genes changed in supertypes that are less abundant in our dataset, such as endothelial and VLMC supertypes. C) Scatterplots of the “gene-dynamic space” in Figure 5 colored by the mean z-scored expression, effect size in the early epoch of CPS and effect size in the late epoch of CPS across the supertypes in the cellular neighborhoods indicated. These plots provide context for the qualitative annotation in Figure 5.

**Extended Data Figure 11:**
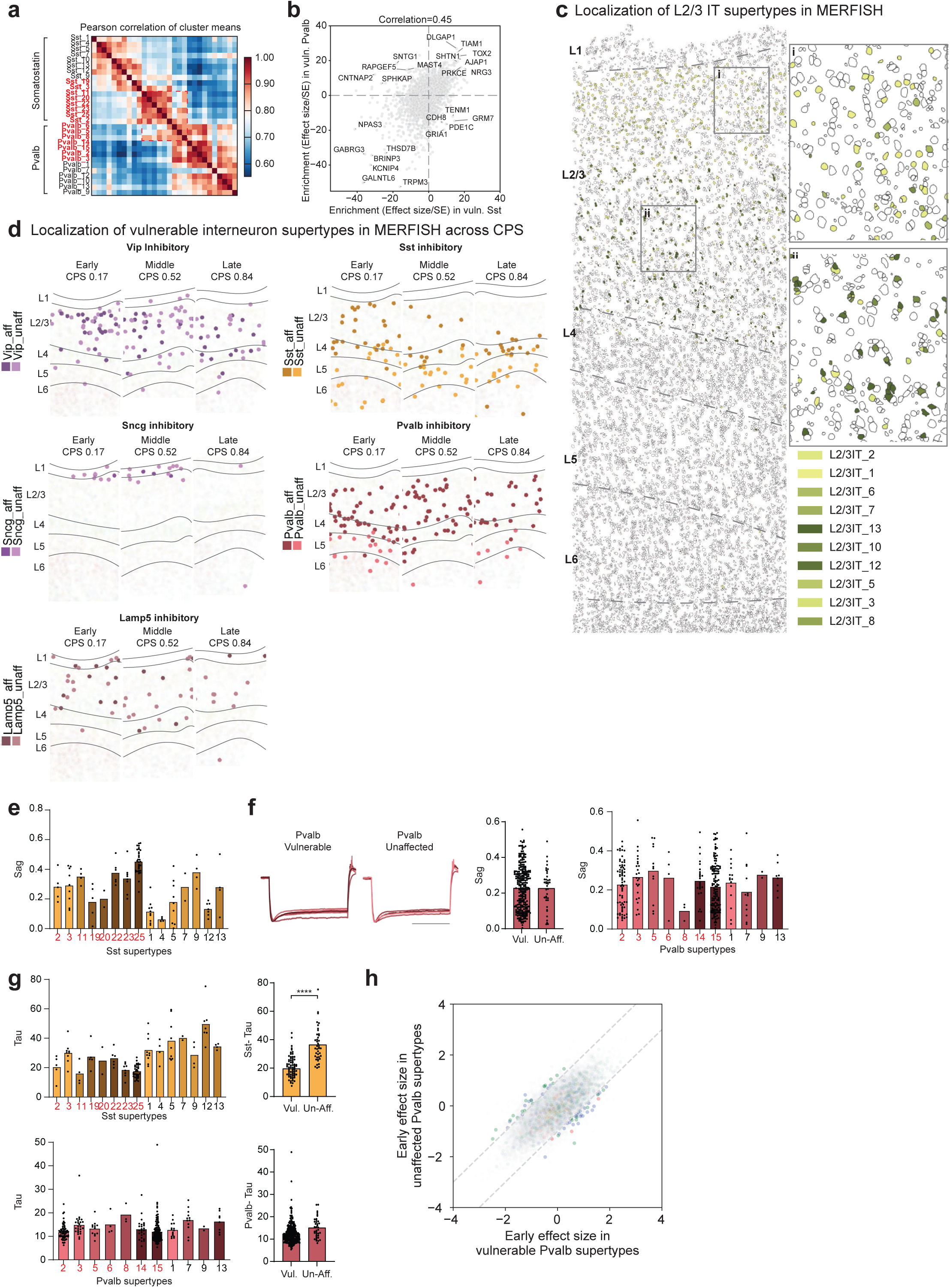
Characteristics of vulnerable neuronal supertypes. A) Heatmap showing the pairwise correlations of the mean expression of all genes across the MGE-derived supertypes indicated. Red labels are vulnerable supertypes; Red dashed box shows high co-correlation among vulnerable Sst and Pvalb supertypes despite them coming from distinct subclasses. B) Scatterplot relating the mean enrichment (defined as the effect size divided by its standard error (SE) from NEBULA) of each gene in vulnerable (vuln) Sst and Pvalb supertypes compared to unaffected types in their respective subclasses. Note many common marker genes in top right corner, but also several subclass specific genes (top left and bottom right corners). C) MERFISH-profiled brain slice in early CPS donor (CPS=0.23) showing each cells location and boundaries defined by the cell segmentation, with cortical layers indicated (L1-L6) and separated by dashed grey lines. Vulnerable L2/3 intratelencephalic (IT) neurons are color-coded. Insets: i) L2/3 IT supertypes have characteristic depths within layers 2 and 3. Supertypes that are known to be in layer 2 are found here, as expected. ii) Supertypes that are known to be found in layer 3 are found here, also as expected. D) Scatterplots showing the spatial locations of individual cells of the inhibitory neuron subclasses indicated from representative MERFISH sections in donors at increasing CPS stages. Vulnerable supertypes (aff) are shown in darker colors and unaffected supertypes (unaff) in lighter ones. Cortical layers (L1-L6) are indicated and separated by solid grey lines. Note, superficial localization of all vulnerable inhibitory neurons. E) Bar and swarm plot showing the Sag values for Sst supertypes from PatchSeq data on non-AD donors. Vulnerable supertypes are colored in red. F) Left, electrophysiological traces showing post-spike hyperpolarization of membrane potential (y-axis) over time in almost all Pvalb neurons from tissue of non-AD human donors that underwent surgical resection. Middle, bar and swarm plot showing sag distributions in individual vulnerable (Vul) and unaffected (Unaff) Pvalb neurons. They are not significantly different. Right, Bar and swarm plot showing the Sag values for Pvalb supertypes from PatchSeq data on non-AD donors. Vulnerable supertypes are colored in red. G) Left top and bottom, Bar and swarm plot showing the Tau apparent membrane time constant values for Sst (top) and Pvalb (bottom) supertypes from PatchSeq data on non-AD donors. Vulnerable supertypes are colored in red. Middle top and bottom, Bar and swarm plots for data on left grouped by vulnerable (vul) and unaffected (un-aff) Sst (top) and Pvalb (bottom) supertypes. P-values for all differential electrophysiological features are in **Supplementary Table 8**. H) Scatterplot relating the mean early effect size of each gene (dots) in vulnerable versus unaffected Pvalb supertypes. The vast majority of genes fall within dashed grey lines bounding the space where effect sizes are within 1 unit of each other. No gene sets were significantly enriched outside of this area (in contrast to Sst supertypes).

## Supplementary Tables

**Supplementary Table 1.** Index of donors, clinical metadata, and modalities profiled

**Supplementary Table 2.** Quantitative neuropathology values

**Supplementary Table 3.** Gene panel for spatial transcriptomic profiling

**Supplementary Table 4.** Model output from scCODA for cellular abundance changes along CPS

**Supplementary Table 5.** Model output from NEBULA for transcriptional changes along CPS

**Supplementary Table 6.** Curated gene sets

**Supplementary Table 7.** Genes differentially expressed by affected Sst and Pvalb supertypes

**Supplementary Table 8.** Differential electrophysiological properties in affected versus unaffected supertypes

## Methods

### SEA-AD cohort selection and brain tissue collection

Brain specimens were donated for research to the University of Washington (UW) BioRepository and Integrated Neuropathology (BRaIN) laboratory from participants in the Adult Changes in Thought (ACT) Study and the University of Washington Alzheimer’s Disease Research Center (ADRC). The study cohort was selected based solely on donor brains undergoing precision rapid procedure (optimized tissue collection, slicing, and freezing) during an inclusion time period at the start of the SEA-AD study, with exclusion of those with a diagnosis of frontotemporal lobar degeneration (FTLD), Down’s syndrome, amyotrophic lateral sclerosis (ALS) or other confounding degenerative disorder (not including Lewy body disease, limbic-predominant TDP-43 encephalopathy, or microvascular brain injury). The cohort was chosen in this manner to represent the full spectrum of Alzheimer’s disease neuropathology, with or without common comorbid age-related pathologies.

The ACT study is a community cohort study of older adults from Kaiser Permanente Washington (KPW), formerly Group Health, in partnership with the UW. The ACT study seeks to understand the various conditions and life-long medical history that can contribute to neurodegeneration and dementia and has been continuously running since 1994, making it the longest running study of its kind. In 2004, ACT began continuous enrollment with the same methods to replace attrition from dementia, dropout, and death, ensuring a consistent cohort of ≥2,000 at risk for dementia. Total enrollment is nearing 6,000, with over 1,000 incident dementia cases; more than 900 have had autopsies to date with an average rate of approximately 45-55 per year. The study completeness of the follow up index is between 95 to 97%. Subjects aged 65 or older without dementia are invited to enroll by random selection from the greater Seattle area patient population of KPW Seattle and undergo bi-annual study visits for physical and mental examinations. In addition to this study data, the full medical record is available for research through KPW. Approximately 25% of ACT autopsies are from people with no MCI or dementia at their last evaluation; roughly 30% meet criteria for MCI, and roughly 45% meet criteria for dementia. Thus, the ACT study provides an outstanding cohort of well-characterized subjects with a range of mixed pathologies including many controls appropriate for this study. Approximately 30% of the ACT cohort consents to research brain donation upon death, and tissue collection is coordinated by the UW BRaIN lab, which preserves brain tissue for fixed, frozen, and fresh preparations (described below), as well as performing a full post-mortem neuropathological examination and diagnosis by Board-certified neuropathologists using the NIA-AA and other relevant, current guidelines.

The UW Alzheimer’s Disease Research Center (ADRC) has been continuously funded by NIH since 1984. It is part of a nationwide network of ADRCs funded through the NIA and contributes uniquely to this premier program through its vision of precision medicine for AD: comprehensive investigation of an individual’s risk, surveillance with accurate and early detection of pathophysiologic processes while still preclinical, and interventions tailored to an individual’s molecular drivers of disease. Participants enrolled in the UW ADRC Clinical Core undergo annual study visits, including mental and physical exams, donations of biospecimens including blood and serum, and family interviews. The UW ADRC is advancing understanding of clinical and mechanistic heterogeneity of Alzheimer’s disease, developing pre-clinical biomarkers, and, in close collaboration with the ACT study, contributing to the state of the art in neuropathological description of the disease. For participants who consent to brain donation, tissue is also collected by the UW BRaIN lab, and is preserved and treated with the same full post-mortem diagnosis and neuropathological work up as described above.

Human brain tissue was collected at rapid autopsy (postmortem interval <12 hours, mean close to 7 hours, **Extended Data Fig. 1a**). One hemisphere (randomly selected) was embedded in alginate for uniform coronal slicing (4mm), with alternating slabs fixed in 10% neutral buffered formalin or frozen in a dry ice isopentane slurry on Teflon-coated plates^30,31^. Superior and Middle Temporal Gyrus (STG-MTG) for quantitative neuropathology was sampled from fixed slabs and subjected to standard processing, embedding in paraffin (**Extended Data Fig. 1b**).

### Single and duplex-IHC for quantitative neuropathology

The STG-MTG tissue blocks were sectioned (cut at 5 µm), deparaffinized by immersion in xylene for 3 minutes, 3 times. Then, rehydrated in graded ethanol (100%, 3x, 96%, 70% and 50% for 3 minutes each) and washed with TBST (Tris Buffered Saline with 0.25% Tween) twice for 3 minutes. The slides were immersed in Diva Decloaker 1x solution (Biocare Medical, DV2004) for heat-induced epitope retrieval (HIER) using the Decloaking Chamber at 110C for 15 minutes for most of the antibodies. For the alpha-Synuclein protein detection, enzymatic antigen retrieval with protein kinase is used. After the HIER is completed, the slides are cooled for 20 minutes at RT. Afterward, the slides are washed with TBST for 5 minutes, twice.

Chromogenic staining was performed using the fully automated BioCare Medical intelliPATH®. Blocking with 3% hydrogen peroxide, Bloxall (Vector Labs), Background punisher (BioCare Medical), and levamisole (Vector labs) is performed to avoid any cross-reactivity and background. The following primary antibodies are used for the first target protein at the dilutions indicated: NeuN (1:500, A60, Mouse, Millipore MAB5374), pTDP43 (1:1000, Ser409/Ser410, ID3, Rat, Biolegend, 829901), Beta Amyloid (1:1000, 6e10, Mouse, Biolegend 80303), Alpha-Synuclein (1:200, LB509, Mouse, Invitrogen 180215) and GFAP (1:1000, Rabbit, DAKO, Z033401-2). Following primary antibody incubation sections were washed 4×2 minutes with TBST and stained with species-appropriate secondary antibody conjugated to a Horseradish Peroxidase (HRP, MACH3-Mouse (M3530), and MACH-Rabbit (M3R531), BioCare Medical). Sections were washed 2×2 minutes with TBST and the antibody complex is then visualized by HRP-mediated oxidation of 3,3’-diaminobenzidine (DAB) by HRP (brown precipitate). Counterstaining is done with hematoxylin after the DAB reaction.

In the case of a duplex IHC (6e10 and pTDP43), the slides were washed 18×2 minutes in TBST and then incubated with primary antibodies at the dilutions indicated after the DAB reaction: IBA1 (1:1000, Rabbit, Wako, 019-19741) and PHF-TAU (1:1000, AT8, Mouse, Thermofisher, MN1020), washed as above and stained with species-appropriate secondary antibodies conjugated to an Alkaline Phosphatase (AP, MACH3-Mouse (M3R532) MACH3-Rabbit (M3R533), Biocare Medical). The complex was then visualized with the intelliPATH® Ferangi Blue reaction kit (IPK5027, Biocare Medical) (blue precipitate). Once staining is completed, the slides were removed from the automated stainer and immersed in TBST, 3 minutes, then dehydrated in graded ethanol (70%, 96%, 100%, 2x) for 3 minutes and xylene (or xylene substitute in the case of double IHC), 3 times each for 3 minutes. Finally, coverslipping is carried out with a Tissue-Tek automated cover slipper (Sakura).

### Image acquisition of whole slide images

To analyze the different slides obtained from the MTG tissue samples processed for IHC, the slides were scanned on the Aperio AT2 digital scanner (Leica), which captures sequential images of a 20x field of view, using slide settings optimized for our IHC protocols which are subsequently assembled or stitched into whole slide images (WSIs) to be exact replicas of the glass slides. All images are scanned at 20x magnification and using the same gain, brightness and exposure times to avoid image to image variations (**Extended Data Fig. 2a**)

### Quantification of whole slide images

The quantitative pathological assessment for the WSIs obtained from the MTG region were analyzed using the HALO® v.3.4.2986 (Indica labs, Albuquerque, New Mexico, USA).

First, DenseNet^194^, a deep learning convolutional neural network was trained to segment MTG cortical ribbon into cortical layers. The DenseNet network is a minimally pretrained classifier developed to recognize patterns in the tissue structure provided by Halo. Training data was created by manually annotating cortical layers labelled with NeuN in 10 cases. Based on the cellular architecture and the relative position withing the cortical ribbon the following layers were annotated: Layer1 (molecular layer), layer 2 (external granular layer), layer 3 (external pyramidal layer), layer 4 (Internal granular layer) and layers 5-6 (internal pyramidal and multiform layers) (Figure 1). Then the trained classifier was applied to the NeuN-labelled sides from all donors. All results of the automatic segmentation were examined by a scientist trained in cortical neuroanatomy and adjusted when necessary. Manual adjustment of the annotations also included removal of staining artifacts and non-parenchymal structures, such as large blood vessels by drawing exclusion areas around them.

Second, using the Serial Section registration tool, all 5 WSIs belonging to the same case (labelled with NeuN, GFAP, α-Syn, Aβ combined with Iba1, and pTau combined with pTDP-43) were registered to each other in order to establish anatomical correspondence between all 5 tissue sections, and the cortical annotations from the NeuN-labelled slide were copied to the other 4 IHC stained slides (noted above). We then applied different algorithms and approaches to obtain stain-specific metrics from all the slides for each cortical layer. Area quantification algorithm (Area Quantification module) was used for determining the area of positive staining for all proteins of interest (NeuN, GFAP, Iba1, Aβ, pTau, α-Syn, and pTDP-43). Multiplex IHC module for used to determine the number of cells displaying positive labelling for NeuN, pTau, α-Syn, and pTDP-43). For the double labelled slides, Multiplex IHC module was used to estimate the area of co-localization of pTau with pTDP-43, and Aβ with Iba1. Microglia Activation module was used to determine the number of cells positive for Iba1, measure the cell process area and length, as well as to classify the cells according to the activation state (activated vs not, based on the process thickness). The same module was adapted to estimate the process length and process area for cells positive for GFAP. In the slides double-labelled for Aβ and Iba1 the Object Colocalization module was used to determine the number of abet Aβ-positive objects (amyloid plaques), the average object area, median object diameter, and the number of objects that were double-positive for Aβ and Iba1.

Development, optimization, and testing of all analysis algorithms was done by a scientist trained in neuropathology. The final quantitative neuropathology dataset includes raw measurements (absolute values) and metrics normalized to the unit area. (**Supplementary Table 2**).

### Creation of Pseudo-progression Score

Our quantitative neuropathological data, *X^m,l^_d_*, is measured in d = 1 … D = number of donors, in l=1 … L cortical layers and m=1 … M distinct neuropathological measurements. To estimate a continuous pseudoprogression score (CPS) of pathological severity in MTG for each brain donor, we created a latent Bayesian statistical model. We assign to each donor a latent variable, termed 𝑡_*d*_ ∈ [0,1], representing CPS. In addition, we propose to infer the most likely donor permutation 𝜋, to facilitate latent space exploration. As described in the main text, the observation model has a mean value dictated by the exponential biophysical dynamics 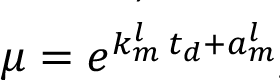, where *k^l^_m_* and *a^l^_m_* are per layer and per QNP measurement dynamic parameters representing rise time and initial condition respectively. We assume that our data is corrupted with observational noise described with a Poisson distribution. We impose Bayesian priors on this model and obtain the following hierarchical Bayesian generative statistical model:

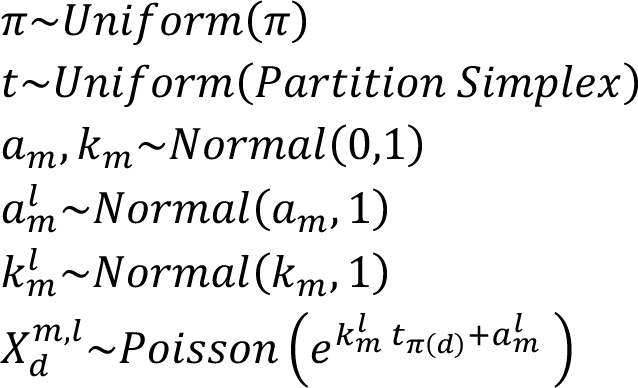

in which the symbol “∼” represents that we are taking draws from a distribution. The hierarchical nature of this model enables the ‘borrowing of information’ across layers and manifests in the fact that, for each measurement, layer specific parameters *k^l^_m_* and *a^l^_m_* are sampled from their population parameters 𝑘_*m*_ and 𝑎_*m*_.

We perform approximate Bayesian inference in this model to obtain draws from an approximate posterior distribution given the model and the underlying priors for a, k, π and t. Our inferential strategy is based on a Gibbs block coordinate sampler where we iteratively sample from each block of variables (t, π or (a,k)) conditioned on the others being fixed. To sample an element t of the simplex that we unequivocally associate with an increasing sequence of times fixed we use the sampler described in^195^. To sample permutations π we resorted to the parametric Gumbel-Sinkhorn family of distributions over permutations^196^ to approximate the otherwise intractable conditional distribution (and hence, our method is approximate). Finally, to sample model parameters (a, k) we used Stan (version 2.34) with 1000 burn out iterations and collect samples from multiple chains ^197^. After initial burned out samples, we iterate through this procedure.

### Tissue processing for single nucleus isolations

Cortical areas of interest were identified on tissue slab photographs taken at the time of autopsy and at the time of dissection using the Allen Human Reference Atlas as a guide for region localization. MTG was sampled at the level of first appearance of the lateral geniculate nucleus corresponding to the intermediate subdivision of area (A) 21. DLPFC was sampled in tissue slabs anterior to the first appearance of the corpus callosum within the superior frontal gyrus corresponding to the rostrodorsal portion of DLPFC (A9^198^). Tissue blocks encompassed the full height of the cortex from pia to white matter (∼5mm) and were ∼2-3 mm wide and 4mm thick. To dissect regions of interest, tissue slabs were removed from storage at –80C, briefly transferred to a –20C freezer to prevent tissue shattering during dissection, and then handled on a custom cold table maintained –20C during dissection. Dissections were performed using dry ice cooled razor blades or scalpels to prevent warming of tissues. Photographs were taken before and after each dissection to document the precise location of each resected tissue block. Dissected tissue samples were then transferred to vacuum seal bags, sealed, and stored at -80C until the time of use. Single nucleus suspensions were generated using a previously described standard procedure (https://www.protocols.io/view/isolation-of-nuclei-from-adult-human-brain-tissue-ewov149p7vr2/v2). Briefly, after tissue homogenization, isolated nuclei were stained with a primary antibody against NeuN (FCMAB317PE, Millipore-Sigma) to label neuronal nuclei. Nuclei samples were analyzed using a BD FACS Aria flow cytometer and nuclei were sorted using a standard gating strategy to exclude multiplets^30^. A defined mixture of neuronal (70% from the NeuN positive gate) and non-neuronal (30% from the NeuN negative) nuclei was sorted for each sample. Nuclei isolated for 10x Genomics v3.1 snRNA-seq were concentrated by centrifugation after FANS and were frozen and stored at –80C until later chip loading. Nuclei isolated for 10x Genomics Multiome and 10x Genomics Single Cell ATAC v1.1 were concentrated by centrifugation after FANS and were immediately processed for chip loading.

### Isolation of RNA and determination of RNA Integrity Number (RIN) from frozen human brain tissue

To assess RNA quality, three tissue samples (roughly 50mg each) were collected from the tissue slab corresponding to the frontal pole of each donor brain. Tissue samples were collected from three different regions of the tissue slab to assess within-slab variability in RNA quality. Dissected tissues were stored in 1.5 mL microcentrifuge tubes on dry ice or in the -80C until the time of RNA isolation. Tissue samples were homogenized using a sterile Takara BioMasher (Takara, 9791A). RNA isolation was performed using either a Qiagen RNeasy Plus Mini Kit (Qiagen, 74134) or a Takara NucleoSpin RNA Plus kit (Takara, 740984) following the manufacturer’s protocol. RNA integrity (RIN) values for each sample were determined using the Agilent RNA 6000 Nano chip kit (Agilent, 5067-1511) and an Agilent Bioanalyzer 2100 instrument following the manufacturer’s protocol.

### 10x genomics sample processing

10x Genomics chip loading and post-processing of the emulsions to sequencing libraries were done with the Chromium Next GEM Single Cell 3’ Gene Expression v3.1, Chromium Next GEM Single Cell ATAC v1.1, and Chromium Next GEM Single Cell Multiome ATAC+Gene Expression kits according to the manufacturer’s guidelines. Nuclei concentration was calculated either manually using a disposable hemocytometer (InCyto, DHC-NO1) or using the NC3000 NucleoCounter.

### 10x sequencing and pre-processing

All 10x libraries were sequenced per manufacturer’s specifications on a NovaSeq 6000 using either a NovaSeq-X or S4 flow cell. Reads were demultiplexed to fastq files using BCL Convert (version 4.2.7) for libraries run on NovaSeq-X flow cells and bcl2fastq (version 2-20-0) for libraries run on S4 flow cells. Reads from snRNA-seq libraries were mapped to 10x Genomics’ official human reference (“Human reference (GRCh38) – 2020-A”) and unique molecular identifiers (UMIs) counted per gene using the cellranger (version 6.1.1) pipeline with the “—include—introns” parameter included. Reads from snATAC-seq and snMultiome libraries were mapped to the same reference using cellranger-atac (version 2.0.0) and cellranger-arc (version 2.0.0) pipelines with default parameters, respectively.

### Identification of donors with low quality tissue

To identify donors with correlated poor tissue-level and pre-sequencing metrics (brain pH, brain weight, postmortem interval, RIN, cDNA amplification concentration, and snRNA-seq library insert size) we constructed an AnnData object using the scanpy^199^ python package (version 1.9.1). Each donor was treated as an “observation” and each quality control metric above as a “variable” instead of the typical cell by gene construction. We then centered and scaled each metric with the scanpy.pp.scale function with default parameters and performed principal component analysis on the matrix with the scanpy.pp.pca function also with default parameters. Two donors were outliers on the first principal component (e.g. had values beyond 1.5 times the interquartile range centered at the median), which were driven by severely low RIN scores and brain pH. These donors were excluded from downstream analyses.

### Identification of severely affected donors

To identify donors with systematically lower data quality by *post-sequencing metrics,* we repeated the above procedure with library-level snRNA-seq and snATAC-seq metrics from the cellranger and cellranger-atac pipelines. The metrics included mean raw reads per cell, median UMIs per cell, median genes detected per cell, fraction of reads mapped to the genome, fraction of reads mapped uniquely to the genome, fraction of reads mapped to intronic regions, fraction of reads mapped to exonic regions, fraction of reads mapped to intergenic regions, fraction of reads mapped antisense, fraction of reads mapped to the transcriptome, fraction of transcriptomic reads in cells, and total genes detected across the library for snRNA-seq and mean raw reads per cell, median fragments detected per cell, fraction of uniquely mapped reads, fraction of the genome in peaks, fraction of fragments overlapping peaks, fraction of fragments overlapping transcription start site (TSS), TSS enrichment, and fraction of transposition events in peaks in cells for snATAC-seq. The first principle component for the snRNA-seq and snATAC-seq metrics explained 71.4% and 65.5% of the variance, respectively, so were taken as composite scores of these highly correlated library level metrics. 11 donors were outliers along the first principle component from both modalities (again defined as values beyond 1.5 times the interquartile range centered at the median), so were flagged as having systematically lower data quality. We classified these donors as “severely affected” based on differences in the steepness of their memory decline in life compared to other donors with similar pathology (see methods section “Testing for differential cognitive slopes in severely affected donors” below and Results section “SEA-AD: Multimodal profiling Alzheimer’s disease progression across wide pathological stages” above). There were no severely affected donors in the snMultiome dataset.

### Testing for differential cognitive slopes in severely affected donors

Cognitive testing was previously co-calibrated and harmonized into cognitive composites for memory, executive functioning, visuospatial recognition, and language for all ACT and UW ADRC study participants using confirmatory factor analyses ^200^. To test for differences in the decline of these domains between severely affected and other high pathology donors, each of the composite scores was standardized to a normal distribution (i.e. *N(0, 1)*) across the broader cohort they came from at baseline. We calculated slopes of memory decline over time using a mixed-effects models using the mixed function from Stata (version 18), where time was parameterized as years before death. We also created a multinomial outcome variable with a reference level consisting of lower pathology donors (i.e. donors with ADNC score Not AD, Low, or Intermediate) and two distinct test groups. The first included donors who were ADNC high, but were not outliers based on library level metrics and the second included donors who were outliers. We then ran a multinomial logistic regression for our outcome variable using the mlogit function from Stata on the slope of each cognitive domain separately, adjusting for age at death and sex and study site (i.e. ACT or UW ADRC). P-values from the models were Bonferroni corrected with the number of cognitive domains tested and called as significant with an alpha value of 0.05.

### Testing for differential quantitative neuropathological features in severely affected donors

We tested for differences in each quantitative neuropathology variable (described above in “Quantification of whole slide images”) using a similar model implemented in the python scikit-learn package (version 1.1.1) using the sklearn.linear_model.LogisticRegression function with default parameters. The multinomial outcome variable was identical to that above and all models adjusted for age at death and sex. Covariates were adjusted with either the minmax_scale (for age at death) or the OneHotEncoder (for sex) functions in sklearn.preprocessing. We then fit the model using the sklearn.linear_model.LogisticRegression.fit function. P-values from the models were Bonferroni corrected with the number of quantitative neuropathology features tested and called as significant with an alpha value of 0.05.

### Comparing peak universes of severely affected donors to other high pathology donors

We used the ChromA ^201^ python package (https://github.com/marianogabitto/ChromA, version 2.1.2) with default parameters on fragment files from each donor individually to call a set of donor specific peaks. As part of this procedure, peaks are filtered by whitelisted regions existing in 10x Cellranger ATAC.

All peak sets were then combined by concatenation. They were then subjected to a fusion condition, namely if 2 peaks shared a 10% overlap their coordinates would be merged using the default bedtools^202^ merge mode. ChromA was used to then compute counts by peaks matrices for each donors using the peak set defined above using fragment and peak files as inputs.

### Transcription factor motif enrichment in peaks specific to severely affected donors

To explore the relationship between Transcription factors (TF) and regions of accessible chromatin specific to severely affected donors, we used the human TF motif list from HOCOMOCO^203^ as an input to FIMO^204^. We collected the list of peaks unique to severely affected donors (cf. Testing for differential peaks in severely affected donors) and retrieved fasta sequences using BED2FASTA. Finally, we run fimo with the following paramters: fimo --oc seq.fa --verbosity 1 --bgfile --nrdb----thresh 1.0E-4 H12CORE_meme_format.meme seq. The results were then filtered on q-value <=.05 and sum aggregated to identify the TF with top represented motifs.

### Creation of “supertypes” in neurotypical reference data

We defined “supertypes” as a set of fine-grained cell type annotations for single nucleus expression data that could be reliably predicted on held-out neurotypical reference data (where “ground truth” labels were assigned as described above) using state-of-the-art machine learning approaches^82,83^ in the scvi-tools python package (version 0.14.3). From 5 neurotypical donors in a related study with roughly 140K nuclei captured with 10x snRNA-seq^26^ we systematically held out labels from 1 donor and used scVI to compute joint latent space then scANVI to iteratively and probabilistically predict their class (3 labels), subclass (24 labels), and then cluster (151 labels). When predicting each nucleus’ class, we selected the top 2,000 highly variable genes (using the scanpy.pp.highly_variable_genes function implemented in the scanpy^199^ python package, version 1.9.1, with the flavor parameter set to “seurat_v3”, n_top_genes parameter set to “2000”) along with the top 500 differentially expressed genes unique to each class (calculated from the reference cells from donors that had their labels retained using a Wilcoxon rank sum test implemented in scanpy.tl.rank_gene_groups with the method parameter set to “wilcoxon”, tie_correct parameter set to “True”, pts parameter set to “True”) to use as features in training the model and specified the donor’s ID and number of genes detected as categorical and continuous covariates, respectively, in the setup_anndata function implemented in scvi.model.SCVI and scvi.model.SCANVI. 2 hidden layers were used for all models, specified by setting n_layer to “2” when initializing the model. The scVI model was then trained using the scvi.model.SCVI.train function with max_epochs set to “200” and passed to scANVI with the scvi.model.SCANVI.from_scvi_model function. The scANVI model was then trained for an additional 20 epochs using the scvi.model.SCANVI.train function. We then obtained the latent representation from the scANVI model with the scvi.model.SCANVI.get_latent_representation function and label predictions with the scvi.model.SCANVI.predict function where the soft parameter was set to “True” to export probabilities. Nuclei were then separated by their predicted class and features were re-selected with the same criteria to predict subclasses and again in predicting clusters. A differential expression test (same Wilcoxon test and parameters as above) was run on clusters with an F1 score below 0.7, and those without 3 positive markers when compared against nuclei from their constituent subclass (cutoff paramters: corrected p-value <0.05, fraction of in-group expression >0.7, fraction out of group expression <0.3) were dropped from the taxonomy, with the remaining clusters representing supertypes.

### Mapping transcriptomic SEA-AD nuclei to reference supertypes

SEA-AD nuclei with transcriptomic data (from either snRNA-seq or snMultiome) with fewer than 500 genes detected were removed upstream of supertype mapping. After defining supertypes in neurotypical donors, we iteratively and probabilistically predicted each SEA-AD nucleus’s class, subclass, and supertype using scANVI^83^, as above. Briefly, each SEA-AD nucleus’ class was predicted after projecting them into a shared latent space with reference nuclei using models trained with 2000 highly variable genes and 500 differentially expressed genes per class (from reference data, where donor name and number of genes were passed as categorical and continuous covariates, respectively). Nuclei were then split by predicted class, projected into a new class-specific latent space where subclass was predicted, and again for supertype. The subclass-specific latent spaces were then used to construct a nearest neighbor graph with the scanpy.pp.neighbors function with default settings and represented with a two-dimensional uniform manifold approximation and projection (UMAP) computed with scanpy.tl.umap with default settings. Predictions from scANVI were evaluated by probabilities from the model and by known marker gene expression (signature scores were computed by summing the absolute value of the t-statistic between reference and SEA-AD nuclei for the top 50 differentially expressed genes for each supertype computed from reference nuclei using the same Wilcoxon test as above). Areas of the nearest neighbor graph with few reference nuclei could represent droplets with ambient RNA, multiplet nuclei, dying cells, or transcriptional states missing from the reference, unique to a donor, or found only in aging or disease. To assess these possibilities, we fractured the graph into tens to hundreds of clusters (called “metacells”) using high resolution Leiden clustering implemented in the scanpy.tl.leiden function with the resolution parameter set to 5 and then merged them based on differential gene expression using the merge_clusters function in the transcriptomics_clustering python package from the Allen Institute (https://github.com/AllenInstitute/transcriptomic_clustering [will be made public upon publication of this manuscript]) with default merging thresholds for gene expression and cluster size. Clusters and metacells were then flagged and removed if they had poor group doublet scores^30^, fraction of mitochondrial reads, number of genes detected, or donor entropy (computed with scipy.stats.entropy with default parameters, version 1.8.1), with cutoffs adjusted for each subclass based on their distributions (to account for dramatically different RNA content found across cell types).

The NIH Brain initiative snRNA-seq dataset generated from the DLPFC of neurotypical reference donors^25^ was mapped to the MTG cellular taxonomy using the same iterative procedure. DLPFC snRNA-seq data from SEA-AD donors were then mapped to the DLPFC neurotypical reference dataset with the predicted MTG labels to ensure a common cellular taxonomy using the same procedure. All downstream quality control steps were also performed identically to those done for the SEA-AD dataset.

### Expanding the reference taxonomy for non-neuronal cells

After removing common technical axes of variation, we next identified nuclei that were transcriptionally distinct from the reference and added them to our supertype taxonomy. To do so, we constructed a new latent space for each subclass using scVI, where the model was passed the supertype predictions as cell labels; gene dispersion was allowed to vary per supertype; sex, race and 10x technology (multiome versus singlome) were included as categorical covariates; and the number of genes detected in each nucleus and the donor age at death were passed as continuous covariates. We then d trained the scVI model and obtained the latent representation using the same functions and parameters described above. Using the neighborhood graph computed from this latent representation, we clustered the nuclei into tens to hundreds of groups and merged them based on differential gene expression using the transcriptomics_clustering package, as above. We defined merged clusters with fewer than 10% of all reference cells or of any single supertype as having poor reference support and added them to the taxonomy (systematically named Subclass_Number-SEAAD). In cases where more than 90% of SEA-AD nuclei within these poorly supported groups were predicted to be one supertype, their new label reflected that assignment (e.g., Subclass_SupertypeNumber_Number-SEAAD). These cell type assignments are used as baseline for the analyses, plots, and tools developed for the web product and this manuscript.

### Mapping epigenomic SEA-AD nuclei to supertypes

To define the peak universe used across all nuclei with epigenomic data (from snATAC-seq and snMulitome), we first separated the 84 donors by their AD neuropathological changes into 4 groups (Not AD, Low, Intermediate, and High) and randomly selected 5 donors from each group (excluding severely affected donors). In each group, we identified group-specific peaks using the atac function in the ChromA^201^ python package (https://github.com/marianogabitto/ChromA, version 2.1.2) with default parameters. We created a union peak set across the 4 groups using the version 2.3.11 bedtoolsmerge function. We then used the count function in ChromA with default parameters to quantify the number of UMIs within each peak to construct a nucleus by peak matrix. We next integrated the snRNA-seq, snATAC-seq, and snMultiome datasets using MultiVI^84^ (from scvi-tools 0.14.3), with modality (e.g. snRNA-seq, snATAC-seq, or snMulitome) set as the batch_key, and donor ID and sex passed to the model as categorical covariates in the scvi.model.MULTIVI.setup_anndata function. After training the model using the scvi.model.MULTIVI.train function with default parameters, we obtained the joint latent representation with the scvi.model.MULTIVI.get_latent_representation function and constructed the nearest neighbor graph across modalities as above and clustered the nuclei using the leiden algorithm implemented in scanpy.tl.leiden with default settings (not high-resolution clustering). We performed quality control on snATAC-seq nuclei based on those from snRNA-seq and snMulitome data in this integrated space. Briefly, we calculated a quality control score for each snATAC-seq nucleus by computing the fraction of its neighbors that were flagged as low-quality snRNA-seq and snMultiome nuclei. snATAC-seq nuclei in leiden clusters with scores greater than 0.5 were removed. We then transferred the subclass labels to snATAC-seq nuclei by labeling them based on what the majority snRNA-seq and snMultiome nearest neighbors. We separated the epigenomic nuclei based on each subclass and called peaks (as above) within subclasses from 5 randomly selected non-SA donors using ChromA to optimize the feature space. Finally, we integrated the multiple modalities data and transferred supertype labels with each subclass using MultiVI, as above.

### Common reprocessing, integration and mapping of publicly available datasets

We obtained raw sequencing reads from 10 publicly available datasets^9–17,98^ that performed single cell or single nucleus RNA-seq on or near the DLPFC of human cohorts that included sporadic AD donors. These included datasets from the AD Knowledge Portal hosted on Synapse: Mathys et al (2019)^9^ (Accession syn18485175, stated brain region “prefrontal cortex/Brodmann area 10”), Zhou et al (2020) (Accession syn21670836, stated brain region “dorsolateral prefrontal cortex”), Olah et al (2020) (Accession syn21438358, stated brain region “dorsolateral prefrontal cortex”), Cain et al (2022) (Accession syn16780177, stated brain region “dorsolateral prefrontal cortex”), Green et al (2023) (Accession syn31512863, stated brain region “dorsolateral prefrontal cortex/Brodmann area 9”), and Mathys et all (2023) (Accession syn52293417, stated brain region dorsolateral prefrontal cortex”). It also included datasets from the Sequencing Read Archive (SRA): Lau et al (2020) (Accession PRJNA662923, stated brain region “prefrontal cortex”), Leng et al (Accession PRJNA615180, stated brain region “superior frontal gyrus”), Morabito et al (2021) (Accession PRJNA729525, stated brain region “prefrontal cortex”), and Yang et al (2022) (Accession PRJNA686798, stated brain region “superior frontal cortex”). From each of these repositories, personal communications with authors, and separate data use agreements with the Rush Alzheimer’s Disease Research Center (for donors from the ROSMAP cohort) we also obtained clinical metadata and harmonized it to a standardized schema included below. The harmonization was done reproducibly, using python code to read in source files, make necessary alterations such as renaming “Braak” 1 to “Braak I”, and write out finalized files.

Reads from each snRNA-seq library were mapped to the same human reference noted above using the same cellranger pipeline as was used for SEA-AD snRNA-seq data. Mapping of nuclei to the SEA-AD cellular taxonomy was done separately from each dataset using the same iterative scVI and scANVI procedure described above to map SEA-AD nuclei from the DLPFC to the neurotypical BRAIN initiative reference dataset. Flagging of low quality nuclei, doublets, and donor-specific nuclei and identification of cell types not in the SEA-AD cellular taxonomy was also done identically to above except neighborhood graphs, leiden clustering, and UMAP visualizations were computed with the GPU-accelerated rapids_singlecell python package (version 0.9.2) with the drop-in replacement functions rapids_singlecell.pp.neighbors, rapids_singlecell.tl.leiden, and rapids_singlecell.tl.umap using default parameters. 3 cell types were added (all perivascular cell types from VINE-seq in Yang et al (2022), as expected), which were found in SEA-AD datasets but in levels too low to drive cluster separation. Mapping results were validated two ways: (1) the probabilities from scANVI for each supertype across each dataset and (2) a signature score computed for each supertype. The top 10 marker genes for each supertype within the SEA-AD dataset compared to all other supertypes within its cellular neighborhood (see **Extended Data Figure 9d**) were identified using the same Wilcoxon test described above. Next, we computed each supertypes’ signature score for each nucleus by transforming the log-normalized expression values (i.e. natural log of UMIs per 10,000 plus 1, computed with the scanpy.pp.normalize_total and scanpy.pp.log1p functions with default parameters) for each of these marker genes to z-scores using the scanpy.pp.scale function (with default parameters) and then taking their mean (**Extended Data Figure 9c**). Closely related supertypes could have similar signature scores, but would retain the same rank order across datasets, if correctly mapped (e.g. Sst_20, Sst_22, Sst_25, Sst_23 and Sst_11 nuclei would all have a high Sst_25 signature score on average, but the order from highest to lowest should be retained across datasets). To test this, we computed the spearman correlation of each supertypes signature score across all supertypes within a cellular neighborhood between the SEA-AD dataset and every other publicly available dataset using the scipy.stats.spearmanr function (scipy version 1.8.1).

#### Common metadata format/specification for every library

- library_prep -(Required) str
- Donor ID - (Required) str
- Brain Region - (Reguired) literal ’MTG’ or ’DLPFC’
- Method - (Required) literal “3’ 10x v2”, “3’ 10x v3”, “3’ 10x v3.1”, “3’ 10x Multiome”, or “5’ 10x v1”
- RIN - float
- barcode – str

#### Common metadata format/specification for every donor

- Donor ID - (Required) str
- Publication - (Required) str
- Primary Study Name - (Required) str
- Age at Death (Required) - int, float or literal ’90+’
- Sex - (Required) Male or Female
- Race (choice=White) - bool
- Race (choice=Black/ African American) - bool
- Race (choice=Asian) - bool
- Race (choice=American Indian/ Alaska Native) - bool
- Race (choice=Native Hawaiian or Pacific Islander) - bool
- Race (choice=Unknown or unreported) - bool
- Race (choice=Other) - bool
- Hispanic/Latino - bool
- Years of education - int or float
- APOE4 Status - (Required) literal ’Yes’ or ’No’
- PMI - (Required) float
- Fresh Brain Weight - float
- Brain pH - float
- Overall AD neuropathological Change - literal “Not AD”, “Low”, “Intermediate”, or “High”
- Thal - literal “Thal 0”, “Thal 1”, “Thal 2”, “Thal 3”, “Thal 4”, or “Thal 5”
- Braak - (Required) literal “Braak 0”, “Braak I”, “Braak II”, “Braak III”, “Braak IV”, “Braak V”, or “Braak VI”
- CERAD score - literal ’Absent’, ’Sparse’, ’Moderate’, or ’Frequent’
- Cognitive Status - (Required) literal ’No dementia’ or ’Dementia’
- Highest Lewy Body Disease - literal ’Not Identified (olfactory bulb not assessed)’, ’Not Identified (olfactory bulb assessed)’, ’Olfactory bulb only’, ’Amygdala-predominant’, ’Brainstem-predominant’, ’Limbic (Transitional)’, or ’Neocortical (Diffuse)’
- LATE - literal ’Unclassifiable’, ’Not Identified’, ’LATE Stage 1’, ’LATE Stage 2’, ’LATE Stage 3’
- Overall CAA Score - literal ’Not identified’, ’Mild’, ’Moderate’, ’Severe’
- Atherosclerosis - literal ’None’, ’Mild’, ’Moderate’, ’Severe’
- Arteriolosclerosis - literal ’None’, ’Mild’, ’Moderate’, ’Severe’

### Spatial transcriptomics gene panel selection

The 140 gene human cortical panel was selected using a combination of manual and algorithmic based strategies requiring a reference single cell/nucleus RNA-seq data set from the same tissue, in this case the human MTG snRNA-seq dataset and resulting taxonomy^30^. First, an initial set of 40 high-confidence marker genes are selected through a combination of literature search and analysis of the reference data. These genes are used as input for a greedy algorithm (detailed below). Second, the reference RNA-seq data set is filtered to only include genes compatible with mFISH. Retained genes need to be 1) long enough to allow probe design (>960 base pairs); 2) expressed highly enough to be detected (FPKM >=10 in at least one cell type cluster), but not so high as to overcrowd the signal of other genes in a cell (FPKM <500 across all cell type clusters); 3) expressed with low expression in off-target cells (FPKM <50 in non-neuronal cells); and 4) differentially expressed between cell types (top 500 remaining genes by marker score, see code below). To sample each cell type more evenly, the reference data set is also filtered to include a maximum of 50 cells per cluster.

The computational step of gene selection uses a greedy algorithm to iteratively add genes to the initial set. To do this, each cell in the filtered reference data set is mapped to a cell type by taking the Pearson correlation of its expression levels with each cluster median using the initial gene set of size n, and the cluster corresponding to the maximum value is defined as the “mapped cluster”. The “mapping distance” is then defined as the average cluster distance between the mapped cluster and the originally assigned cluster for each cell. In this case a weighted cluster distance, defined as one minus the Pearson correlation between cluster medians calculated across all filtered genes, is used to penalize cases where cells are mapped to very different types, but an unweighted distance, defined as the fraction of cells that do not map to their assigned cluster, could also be used. This mapping step is repeated for every possible n+1 gene set in the filtered reference data set, and the set with minimum cluster distance is retained as the new gene set. These steps are repeated using the new get set (of size n+1) until a gene panel of the desired size is attained. Code for reproducing this gene selection strategy is available as part of the mfishtools R library (https://github.com/AllenInstitute/mfishtools).

### Spatial transcriptomics data collection

Human postmortem frozen brain tissue was embedded in Optimum Cutting Temperature medium (VWR 25608-930) and sectioned on a Leica cryostat at -17C at 10 μm onto Vizgen MERSCOPE coverslips. These sections were then processed for MERSCOPE imaging according to the manufacturer’s instructions. Briefly: sections were allowed to adhere to these coverslips at room temperature for 10 minutes prior to a 1 minute wash in nuclease-free phosphate buffered saline (PBS) and fixation for 15 minutes in 4% paraformaldehyde in PBS. Fixation was followed by 3×5 minute washes in PBS prior to a 1 minute wash in 70% ethanol. Fixed sections were then stored in 70% ethanol at 4C prior to use and for up to one month. Human sections were photobleached using a 240W LED array for 72 hours at 4C (with temperature monitoring to keep samples below 17C) prior to hybridization then washed in 5 mL Sample Prep Wash Buffer (VIZGEN 20300001) in a 5 cm petri dish. Sections were then incubated in 5 mL Formamide Wash Buffer (VIZGEN 20300002) at 37C for 30 min. Sections were hybridized by placing 50 μL of VIZGEN-supplied Gene Panel Mix onto the section, covering with parafilm and incubating at 37 C for 36-48 hours in a humidified hybridization oven. Following hybridization, sections were washed twice in 5 mL Formamide Wash Buffer for 30 minutes at 47C. Sections were then embedded in acrylamide by polymerizing VIZGEN Embedding Premix (VIZGEN 20300004) according to the manufacturer’s instructions. Sections were embedded by inverting sections onto 110 μL of Embedding Premix and 10% Ammonium Persulfate (Sigma A3678) and TEMED (BioRad 161-0800) solution applied to a Gel Slick (Lonza 50640) treated 2×3 inch glass slide. The coverslips were pressed gently onto the acrylamide solution and allowed to polymerize for 1.5 hours. Following embedding, sections were cleared for 24-48 hours with a mixture of VIZGEN Clearing Solution (VIZGEN 20300003) and Proteinase K (New England Biolabs P8107S) according to the manufacturer’s instructions. Following clearing, sections were washed 2×5 minutes in Sample Prep Wash Buffer (PN 20300001). VIZGEN DAPI and PolyT Stain (PN 20300021) was applied to each section for 15 minutes followed by a 10 minutes wash in Formamide Wash Buffer. Formamide Wash Buffer was removed and replaced with Sample Prep Wash Buffer during MERSCOPE set up. 100 μL of RNAse Inhibitor (New England BioLabs M0314L) was added to 250 μL of Imaging Buffer Activator (PN 203000015) and this mixture was added via the cartridge activation port to a pre-thawed and mixed MERSCOPE Imaging cartridge (VIZGEN PN1040004). 15 mL mineral oil (Millipore-Sigma m5904-6X500ML) was added to the activation port and the MERSCOPE fluidics system was primed according to VIZGEN instructions. The flow chamber was assembled with the hybridized and cleared section coverslip according to VIZGEN specifications and the imaging session was initiated after collection of a 10X mosaic DAPI image and selection of the imaging area. Specimens were imaged and automatically decoded into transcript location data and a cell by gene table. All postprocessing and segmentation was completed using the vizgen-postprocessing docker container version 0.0.5 (https://github.com/Vizgen/vizgen-postprocessing). For each section, segmentation was run on a single z plane (z=z3). Segmentation was a combination of cellpose-cyto2 2d segmentation (with clahe, Contrast Limited Adaptive Histogram Equalization, normalized DAPI and PolyT images as inputs) and cellpose nuclei-only segmentation (using clahe normalized DAPI images only). Results were then fused using the “harmonize” strategy and returned as cell metadata summary files and parquet mosaic geometry files. If segmentation failed on the z=z3 image plane, z=z4 image data was used instead.

### Spatial transcriptomics data quality control and mapping

Resulting transcript location data and cell by gene tables were assessed for quality by comparing total transcript counts across specimens. A rectangular region was selected in each section to encompass a region spanning pia to white matter with uniform layer thickness and minimal in-plane cortical curvature. Transcript counts within these regions were summed to create a spatial transcriptomics pseudo-bulk profile. This pseudo-bulk profile was consistent with the bulk RNASeq measurements summed across 10 donors (Pearson correlation 0.69). Two sections with total transcript correlation less than 0.6 to the spatial transcriptomic pseudo-bulk were eliminated, along with two sections that measured unusually high counts of one gene (HS3ST2). Within the cortical selections, layers were annotated manually based on excitatory subclass annotations and cellular density. After these steps, selected cells from 69 sections from 27 donors formed our spatial dataset for subsequent analysis. Cells were eliminated from further analysis if they fell outside the following criteria: >15 genes detected, 30-4000 total transcripts detected, 100-4000 um^3^ total cell volume. Cells in this dataset had a mean of 210.9 detected transcripts, and mean volume of 1292 μm^3^.Cells in the spatial transcriptomics dataset were mapped to the integrated taxonomy at the supertype level by finding the supertype whose mean gene expression within the supertype best matched to each cell. Specifically, we used a bootstrapped Pearson correlation in the map_cells_knn_bs function in the scrattch-mapping R package version 0.16.

### Compositional analysis of supertypes

To model changes in the composition of cell types as a function of CPS and other covariates we used the Bayesian method scCODA^96^ (version 0.1.7). We tested compositional changes in neuronal and non-neuronal nuclei separately, as they were sorted to have a defined ratio (70% neuronal nuclei, 30% non-neuronal nuclei in each donor). To do this, we created separate AnnData objects of neuronal and non-neuronal nuclei with supertype annotations, sequencing library IDs and relevant donor-level covariate information (noted below) for all snRNA-seq and snMultiome nuclei formatted per https://sccoda.readthedocs.io/en/latest/data.html using the sccoda.util.cell_composition_data function with cell_type_identifier set to supertype and sample_identifier set to the sequencing library ID. As we did not know which supertypes would be affected by AD, we ran models with each supertype set as the unchanged reference population, as recommended by scCODA’s authors. We setup an ensemble of models to test whether supertypes were credibly affected across cognitive status (No dementia [0] versus Dementia [1]), ADNC (Not AD [0], Low [1/3], Intermediate [2/3], High [1]), and CPS (Interval [0,1]) using the scconda.util.comp_ana.CompositionalAnalysis function with formula set to “Sex + Age at death + Race + 10x Chemistry + APOE4 Status + [disease covariate]” with each supertype as the reference population (yielding 417 models total) and obtained posterior estimates for each parameter with a Markov chain Monte Carlo (MCMC) process implemented in the sample_hmc function with default parameters. The sampling occasionally stayed at fixed points, so we re-ran models with fewer than 60% accepted epochs. We defined credibly affected supertypes as those that had a mean inclusion probability across models >0.8. The same approach was used for testing compositional changes across CPS in snRNA-seq data from SEA-AD DLPFC dataset using the formula “Sex + Age at death + Race + [disease covariate]” and across ADNC in snRNA-seq data from Green et al (2023)^17^ and Mathys et al (2023)^98^ using the formula formula “Sex + Age at death + APOE4 Status + [disease covariate]”, with the sample_identifier set to Donor ID as there was not a one-to-one or one-to-many relationship between donors and libraries across these datasets.

### Gene expression changes along CPS

To model gene expression changes along CPS while adjusting for other covariates and pseudo-replication within donors we used a general linear mixed effects model implemented in the NEBULA R package^205^ (version 1.2.0) accessed in python via the rpy2 package (version 3.5.2). We used objects with all nuclei and with nuclei divided into the first (<0.55, “early”) and second (>0.45, “late”) disease phase along CPS. Briefly, we split CPS into two phases: (1) A period where donors had normal cognition and relatively low levels of plaque and tangle pathology (but changes in other quantitative neuropathology variables) and (2) a period where donors had markedly increased levels of plaque and tangle pathology and cognitive decline. To delineate the cutoffs for these phases, we interrogated the general additive models used to relate the number of amyloid plaques and tau tangles to CPS. Significant coefficients were first observed for splines starting at 0.4 and 0.6, depending on the variable and layer. We chose CPS=0.5 as the intermediary cut point but added a small amount of overlap to account for uncertainty precisely when the transition occurs. For each supertype, we constructed a model matrix from relevant metadata with the base model.matrix function in R with the formula “Sex + Age at death + Race + 10x Chemistry + CPS + Number of genes detected” after standardizing numerical values to a [0,1] interval. We randomly added single pseudocounts to 3 nuclei to features that had zero values across all nuclei within a supertype in the metadata groupings, which would have prevented the model from properly fitting coefficients (e.g. the X-chromosome gene **XIST** had zero counts across all nuclei from **male** donors, so 1 pseudocount for XIST was added to 3 random male nuclei). We then grouped raw count and model matrices with the group_cell function in NEBULA, passing the counts matrix to count, the model matrix to pred, the number of UMIs detected in each nucleus to offset, and the donor IDs as the random effect to id. To fit the model, we then ran the nebula function using the output of group_cells. We filtered genes with fewer than 0.005 counts per nucleus (as recommended) which resulted in coefficients for roughly 14,000 genes being fit in each supertype. We further restricted the results to genes with convergences equal to 1. We determined the number of significant genes from the resulting p-values in each supertype with the Benjamini-Hochberg procedure with an alpha threshold of 0.01.

### Construction of gene dynamic space

To identify patterns in gene expression dynamics, we constructed a matrix spanning all genes on one axis and their corresponding normalized early and late beta coefficients (divided by their standard errors) from NEBULA (see above) as well as z-scores of the mean expression (capped at a magnitude of 2) for each supertype along the other axis. Genes that were not tested in a particular supertype because of low counts per cell were assigned a beta coefficient of 0. We then computed a nearest neighbor graph across all genes using Euclidian distances with the scanpy.pp.neighbors function with use_rep set to “X” and n_neighbors set to 15. To visualize the resulting graph, we computed a low dimensional UMAP representation with the scanpy.ul.umap function with default parameters and computed mean normalized beta coefficient and z-score values across classes and subclasses for visualization purposes.

### Curation of gene sets

We constructed 31 gene sets that encompass molecular processes important for neurons or implicated in AD from various databases and literature sources noted below. The gene lists are compiled in **Supplementary Table 6**.

- Electron transport chain, based on Gene Ontology (GO) 0022900
- Glycolysis, based on BIOCYC Pathway PWY66-400
- Cholesterol biosynthesis, BIOCYC Pathways PWY66-341, PWY66-3, PWY66-4
- Steroid metabolism, based on UniuProt Keyword KW-0753 (reviewed only)
- Fatty acid metabolism, based on UniProt Keyword KW-0276 (reviewed only)
- Phospholipid metabolism, based on UniProt Keyword KW-1208 (reviewed only)
- Sphingolipid metabolism, based on UniProt Keyword KW-0746 (reviewed only)
- Ribosomal proteins, based on GO 0006412
- Eukaryotic initiation, elongation and termination factors, based on GO 0006412
- Transcriptional machinery, based on GO 0006351
- Ubiquitin machinery (split by Category), based on Unibet 2.0^206^
- Kinases (split by group), based on KinHub^207^
- Voltage gated ion channels, based on the Guide to Pharmacology (GtP)^144^
- Ligand gated ion channels, based on the Guide to Pharmacology (GtP)
- Nuclear hormone receptors, based on the Guide to Pharmacology (GtP)
- Transcription factors (split by Family) based on the Guide to Pharmacology (GtP)
- Genes identified by Genome Wide Association Studies (GWAS)^208^
- Cell adhesion, based on GO 0007155
- Actin-Spectrin-Spetin cytoskeleon, based on GO 0005200, 0003779, 0030507, and 0031105
- Microtubule cytoskeleton, based on GO 0005200, 0015630, 0008017
- Molecular motors used in vesicle trafficking, based on Hirokawa et al^209^
- Cargo adaptors used in vesicle trafficking, based on Hirokawa et al^209^ and GO 0030705 and 0016192
- Axonal guidance cues, based on GO 0097485
- Neuropeptides, based on Smith et al^210^
- Neuropeptide receptors, based on Smith et al^210^
- Myelin components, based on Morell et al^211^
- OPC differentiation and re-myleination program, based on Long et al^161^, Nakatani et al^159^, Wang et al^155^ Genoud et al^156^, Zhang et al^158^, and Tomassy et al^160^
- Fc receptors, based on Owen et al^212^
- Major histocompatibility complex class II, based on Jones et al^213^
- Human plaque induced genes, based on Chen et al^20^
- Interferon stimulated genes, based on Schneider et al^214^

### Gene set enrichment analysis

We employed a bootstrapping procedure to test for significant enrichment of each gene set in the early or late AD epoch along CPS, in specific cell types. Briefly, for each gene set we randomly selected the same number of genes within it 1000 times with the numpy.random.choice (version 1.22.4) function with replace set to “False”. For each iteration, we computed the mean early and late beta coefficients for the randomly chosen set of genes to create a background distribution for each AD epoch. We then computed a z-score for the actual gene set by computing the mean early and late beta coefficients for the genes within the set, subtracting the mean from the null distributions from them and dividing them by the standard deviation from the null distributions. We computed p-values for these z-scores using the scipy.stats.norm.cdf python function. We applied a Bonferroni correction for the number of gene lists tested and set an alpha threshold of 0.05.

### Identification of marker genes in subclasses with vulnerable and disease associated supertypes

We used the same general linear mixed effects model (NEBULA^205^) to test for supertype specific expression within subclass. All parameters were the same for the test across CPS, except we constructed a model matrix from relevant metadata with the base model.matrix function in R using the formula “Cell type + Sex + Age at death + Race + 10x Chemistry + Number of genes detected” after standardizing numerical values to a [0,1] interval. Cell type was encoded as 1 for the supertype being tested and 0 for all other supertypes within the subclass.

### Gene regulatory networks

To compute gene regulatory networks (GRNs) within non-neuronal subclasses, we used the SCENIC+^215^ python package (version 1.0.1.dev3+g3741a4b). Briefly, we first created a fragment file that contained data from all SEA-AD nuclei within each non-neuronal subclass to call subclass-specific peaks using the MACS2^216^ package, as recommended by SCENIC+. We constructed an ArchR^217^ R object (version 1.0.1) from the fragments file and called peaks within each subclass using MACS2 implemented in the addGroupCoverages and addReproduciblePeakSet functions in ArchR with the groupBy parameter set to the subclass labels. We then exported these cell by peak matrices and created pycisTopic objects (version 1.0.3.dev18+ge563fb6). We used the pycisTopic.cistopic_class.run_cgs_models function with n_topics set to “[2,4,10,16,32,48]”, n_cpu set to “32”, and n_iter set to “500” to determine the appropriate number of topics and settled on 16 based on results from the pycisTopic.lda_models.evaluate_models function with select_model set to “16”, return_model set to “True” and the SCENIC+ usage guide. To select candidate enhancer elements, we then binarized the topics using the pycisTopic.topic_binarization.binarize_topics function once with method set to “otsu” and once with method set to “ntop” and ntop set to “3000”. We also identified differential features with the pycisTopic.diff_features.find_highly_variable_features and pycisTopic.diff_features.find_diff_features functions with default parameters after imputing and normalizing the cisTopic object with the pycisTopic.diff_features.impute_accessibility and pycisTopic.diff_features.normalize_scores functions with scale_factor set to “10**6” and “10**4”, respectively. These features were used to define region sets that were associated with transcription factors with the scenicplus.wrappers.run_pycistarget function with species set to “homo_sapiens”, ctx_db_path set to the “hg38_screen_v10_clust.regions_vs_motifs.rankings.feather” file obtained from the SCENIC+ guide, dem_db_path set to the “hg38_screen_v10_clust.regions_vs_motifs.scores.feather” file also obtained from the SCENIC+ guide, path_to_motif_annotations set to the “hg38_screen_v10_clust.regions_vs_motifs.scores.feather” file also obtained from the SCENIC+ guide, run_without_promoters set to “True”, n_cpu set to “32”, and the annotation_version set to “v10nr_clust”. We then passed the final pycisTopic object along with the snRNA-seq data for the non-neuronal subclasses to SCENIC+ with the scenicplus.scenicplus_class.create_SCENICPLUS_object function with the key_to_group_by set to the nuclei subclasses and nr_cells_per_metacells set to “5”. We then identified GRNs with the scenicplus.wrappers.run_scenicplus function with variable set to the subclass labels, species set to “hsapiens”, assembly set to “hg38”, tf_file set to the “utoronto_human_tfs_v_1.01.txt” file obtained from the University of Toronto Human Transcription Factor database, upstream set to “[1000, 150000]”, downstream set to “[1000, 150000]”, calculate_TF_eGRN_correlation set to “True”, calculate_TF_eGRN_correlation set to “False”, and n_cpu set to “32”. Finally, only gene regulatory networks with a rho value for “TF_cistrome_correlation” greater than 0.7 or less than -0.7 were retained, per the SCENIC+ guidelines. We then identified transcription factors within the GRNs that had both subclass-specific expression (z-score mean gene expression greater than 2) and increased in the early AD epoch (mean NEBULA early beta coefficient greater than 1) and identified predicted downstream target genes that were common across all of them.

### Identifying differential electrophysiological features in vulnerable neurons from PatchSeq data

We obtained publicly available PatchSeq^29,38,39^ data from 2,602 cells, originating from slices from 401 donors. These cells were recorded in healthy tissue extracted in surgical resection due to cancer pathology or epilepsy (95% of cases) and hydrocephalus, encephalomyelitis, aneurism, and ventriculoperitoneal shunt (5%). We subsetted the dataset to include only cells obtained from the MTG and mapped the snRNA-seq data from them to the SEA-AD MTG cellular taxonomy using the same iterative scVI and scANVI approach described above for SEA-AD nuclei and the publicly available datasets.

We tested for differences in each electrophysiological feature between vulnerable and unaffected neurons in the Sst and Pvalb using a logistic regression implemented in the python scikit-learn package (version 1.1.1) using the sklearn.linear_model.LogisticRegression function with default parameters. The outcome variable was 0 for unaffected supertypes and 1 for affected ones and all models adjusted for age at death, sex, and whether the slices were cultured or not. Covariates were adjusted with either the minmax_scale (for age at death) or the OneHotEncoder (for sex and culture status) functions in sklearn.preprocessing. We then fit the model using the sklearn.linear_model.LogisticRegression.fit function. P-values from the models were Bonferroni corrected with the number of quantitative neuropathology features tested and called as significant with an alpha value of 0.05.

## Code availability

We will release the collection of scripts used to annotate SEA-AD and publicly available data and perform analysis on the Allen Institute GitHub page.

## Data availability

FASTQs containing sequencing data from snRNA-seq, snATAC-seq, and snMultiome assays are available through controlled access at Sage Bionetworks (accession: <u>syn26223298</u>). Nuclei by gene matrices with counts and normalized expression values from snRNA-seq and snMultiome assays are available through the <u>Open Data Registry</u> in <u>AWS bucket</u> as AnnData objects (h5ad), and viewable on the <u>cellxgene platform</u> and <u>ABC Atlas</u>. Nuclei by peak matrices for the snATAC-seq data (with peaks called across all nuclei) are in the same bucket. Cell by gene matrices containing spatial coordinates from MERFISH data are in a <u>separate bucket</u>. MERFISH data is also viewable on the <u>ABC Atlas</u>. Donor, library, and cell-level metadata are available in these objects and also on SEA-AD.org. Raw images from the quantitative neuropathology data are available on the Open Data Registry on AWS in yet another <u>bucket</u> and the variables derived from HALO on SEA-AD.org. We are working to obtain approval to share the publicly available datasets under the data use agreements that govern them. In the meantime, we have placed the cell type annotations on <u>AWS</u> without gene expression data or donor metadata.

